# 5S-IGS rDNA in wind-pollinated trees (*Fagus* L.) encapsulates 55 million years of reticulate evolution and hybrid origins of modern species

**DOI:** 10.1101/2021.02.26.433057

**Authors:** Simone Cardoni, Roberta Piredda, Thomas Denk, Guido W. Grimm, Aristotelis C. Papageorgiou, Ernst-Detlef Schulze, Anna Scoppola, Parvin Salehi Shanjani, Yoshihisa Suyama, Nobuhiro Tomaru, James R.P. Worth, Marco Cosimo Simeone

## Abstract

Standard models of plant speciation assume strictly dichotomous genealogies in which a species, the ancestor, is replaced by two offspring species. The reality in wind-pollinated trees with long evolutionary histories is more complex: species evolve from other species through isolation when genetic drift exceeds gene flow; lineage mixing can give rise to new species (hybrid taxa such as nothospecies and allopolyploids). The multi-copy, potentially multi-locus 5S rDNA is one of few gene regions conserving signal from dichotomous and reticulate evolutionary processes down to the level of intra-genomic recombination. Therefore, it can provide unique insights into the dynamic speciation processes of lineages that diversified tens of millions of years ago. Here, we provide the first high-throughput sequencing (HTS) of the 5S intergenic spacers (5S-IGS) for a lineage of wind-pollinated subtropical to temperate trees, the *Fagus crenata – F. sylvatica* s.l. lineage, and its distant relative *F. japonica.* The observed 4,963 unique 5S-IGS variants reflect a complex history of hybrid origins, lineage sorting, mixing via secondary gene flow, and intra-genomic competition between two or more paralogous-homoeologous 5S rDNA lineages. We show that modern species are genetic mosaics and represent a striking case of ongoing reticulate evolution during the past 55 million years.

**Significance statement:** The evolution of extra-tropical wind-pollinated tree genera involves dynamic speciation processes. High-throughput sequencing of the multi-copy, potentially multi-locus 5S rDNA reveals a complex history of hybrid origins, lineage sorting and mixing, and intra-genomic competition between paralogous-homeologous loci in the core group of Eurasian beech trees (genus *Fagus*) and their distant relative, *F. japonica*. The modern species are genetic mosaics and represent a striking case of at least 55 million years of ongoing reticulate evolution.

## Introduction

Wind-pollination is a derived condition in most angiosperms and likely evolved in response to changing environmental conditions, with the grass family (Poaceae) being the most prominent example of a strictly wind-pollinated group (Friedman & Barrett, 2008). One of the most diverse and ecologically important groups of wind-pollinated woody angiosperms is the order Fagales including walnuts, birches, oaks and beech trees. Fagales lineages repeatedly shifted from biotic insect- to abiotic wind-pollination (Endress, 1977; Manos et al., 2001; Friis et al., 2011). Adaptations to wind-pollination were accompanied by spectacular changes in reproductive biology. Major innovations included spring flowering, the simplification of flowers and specialization of inflorescences, large stigmatic surfaces, lax male inflorescences, increase in stamen numbers, large anthers, high pollen production, and thin-walled pollen grains (Endress, 1977; Denk & Tekleva, 2014). The most striking innovation in the life history of many wind-pollinated Fagales groups is the delay of ovule development in relation to pollination (Endress, 1977). The absence of leaves facilitates pollination, whereas the beginning of the growing season favours fruit development. The general view is that the shift to wind-pollination in the Fagales promoted the group’s rise to dominance in temperate forest ecosystems (Endress, 1977). However, how this costly change in reproductive biology affected speciation is not well understood (Friedman & Barrett, 2008).

Assessing processes involved in the origin and diversification of species is a central goal of taxonomy, ecology, biodiversity and phylogeny research (De Queiroz, 2007; Mallet, 2013; Scornavacca et al., 2020). In plants, these processes include cross-lineage genomic admixture, hybridization events and ancestral gene flow between allopatric and highly disjunct modern taxa (Willyard et al., 2009; Klein & Kadereit, 2016; Marques et al., 2016; Suarez-Gonzalez et al., 2016; Folk et al., 2017, 2018; McVay et al., 2017). Especially in trees, characterized by long generation times and large effective population size, wind-pollination acts as a natural catalyst for reticulate evolutionary processes such as chloroplast capture, introgression and increased interpopulation gene flow. Notably, plastid DNA variation is sorted by geography rather than speciation in all broadly sampled groups of Fagales studied so far (Fagaceae: Simeone et al., 2016, Yan et al., 2019; Nothofagaceae: Acosta & Premoli, 2010).

*Fagus* L. (Fagaceae) provides an exemplary case. It is a small but ecologically and economically important genus of about ten monoecious tree species occurring in three isolated regions of the Northern Hemisphere: East Asia, western Eurasia, and eastern North America (Shen, 1992; Peters, 1997; Denk, 2003; Fang & Lechowicz, 2006). The genus probably originated at high latitudes during the Paleocene (western Greenland, northeast Asia; Fradkina et al., 2005; Denk & Grimm, 2009; Grímsson et al., 2016). It is subdivided into two informal subgenera (Shen, 1992; Denk et al., 2005). “Subgenus Engleriana” includes three (North-)East Asian species: *Fagus engleriana* Seemen ex Diels*, F. multinervis* Nakai, and *F. japonica* Maxim. “Subgenus Fagus” includes five or more Eurasian species: *Fagus sylvatica* L. s.l. (including *F. orientalis* Lipsky) in western Eurasia; *F. crenata* Blume in Japan; *F. lucida* Rehder & E.H.Wilson*, F. longipetiolata* Seemen and *F. hayatae* Palib. (incl. *F. pashanica* C.C.Yang) in central and southern China and Taiwan, and a single North American species, *F. grandifolia* Ehrh. (including *F. mexicana* [Martínez] A.E.Murray). These two lineages diverged by the early Eocene (53 [62–43] Ma; Renner et al., 2016). While the lineage leading to the modern genus is at least 82–81 myrs old (Grímsson et al., 2016), extant species are the product of ~50 myrs of trans-continental range expansion and phases of fragmentation leading to diversification (Denk, 2004; Denk & Grimm, 2009; Renner et al., 2016). These dynamic migration and speciation histories left multifarious morphological and molecular imprints on modern members of the genus (Fig. 1).

**Fig. 1.**
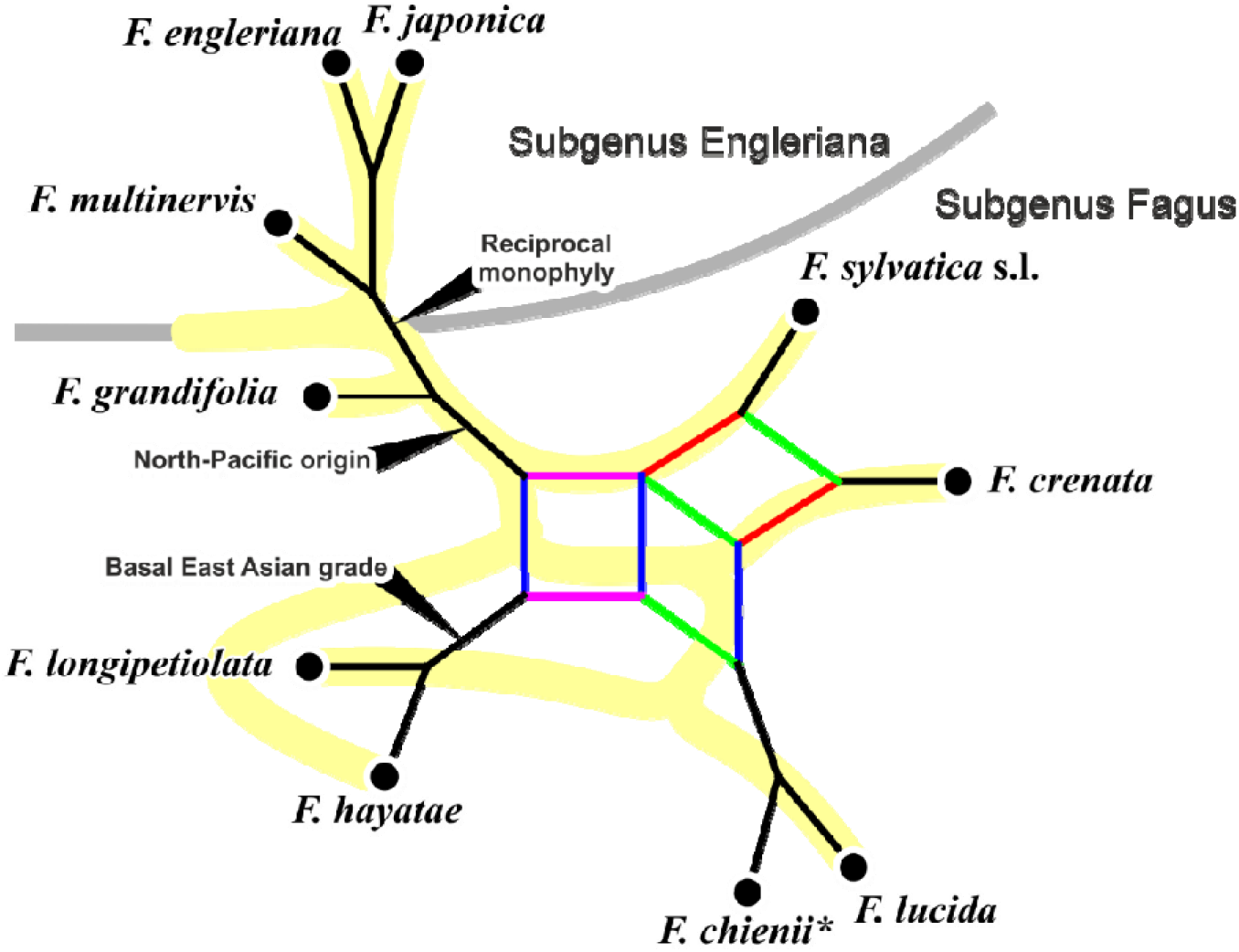
Synoptical consensus network depicting known species relationships in *Fagus* (modified from Denk & Grimm, 2009, fig. 2). Alternative rooting scenarios (discussed in Denk & Grimm, 2009; Renner et al., 2016) indicated by black arrows. Black edges – non-conflicting phylogenetic splits; red – *crenata-sylvatica* lineage (within a clade including *F. lucida,* purple edge bundle) is supported by morphology/fossil record (Denk, 2003; Denk & Grimm, 2009), ITS data capturing substantial intra-individual polymorphism (Denk et al., 2005; Göker & Grimm, 2008), and when ITS individual consensus sequences are combined with *LEAFY* intron data (Renner et al., 2016); green – a closer relationship between *F. crenata* and *F. lucida* is indicated by intra-individual ITS polymorphic sequence patterns (Grimm et al., 2007); blue – a (semi-)continental East Asian clade (China, Taiwan) agrees well with morphology (Denk, 2003) and *LEAFY* intron data (Oh et al., 2016; Renner et al., 2016); yellow branches – species tree inferred by Jiang et al. (2021). *, there are no molecular data for *F. chienii* morphotypes; the species is probably a synonym of *F. lucida.*

Intra- and inter-specific phylogenetic relationships within *Fagus* have been difficult to resolve (Denk et al., 2002, 2005; Renner et al., 2016). In western Eurasia, for instance, where beech is the dominant climax species in mid-altitude forests, the number and rank of several taxa is still controversial (e.g. Gömöry et al., 2018). Difficulties arise from a low inter-specific but high intra-specific morphological diversity (e.g. Denk, 1999a, b) and equally complex inter- and intra-specific genetic differentiation in both the nucleome and plastome of beeches (Denk et al., 2002; Okaura & Harada, 2002; Grimm et al., 2007; Gömöry & Paule, 2010; Hatziskakis et al., 2009; Lei et al., 2012; Zhang et al., 2013). Most recently, Jiang et al. (2021) combined the data of 28 nuclear genes to infer a fully resolved (sub-)species tree using reconstructed haplotypes. This tree poorly reflects the signal in their gene sample indicating multiple events of gene duplication, incomplete lineage sorting below and above the species-level, and gene flow via secondary contact. Accordingly, ancient and reiterated hybridization/ introgression events were inferred for *Fagus* based on all assembled morphological, fossil and molecular data, and with respect to its biogeographic history and ecology (Denk et al., 2005; Denk & Grimm, 2009; Oh et al., 2016; Renner et al., 2016). Thus, *Fagus* constitutes a perfect example to study genetic complexity in a wind-pollinated genus of dominant forest trees.

Nuclear ribosomal DNA spacers, organized in multi-copy and potentially multi-loci arrays (Symonová, 2011), have great potential to detect past and recent reticulation events (e.g. Grimm & Denk, 2008, 2010). For *Fagus sylvatica*, Ribeiro et al. (2011) found two paralogous 5S rDNA loci, suggesting chromosomal rearrangements during its evolution. Although not studied in *Fagus*, intra- and inter-array variation can thus be expected. Availability of High-Throughput Sequencing (HTS) approaches is now boosting new efforts into research questions that were previously considered too expensive and labour-intensive (Babik et al., 2009; Glenn, 2011). In a recent study, Piredda et al. (2021) generated HTS amplicon data of the non-transcribed intergenic spacers of the 5S rDNA in *Quercus* (Fagaceae) and demonstrated the great value of this approach to provide phylogenetic information for inspecting range-wide evolutionary patterns while circumventing previous technical limitations (e.g. cloning, abundant samples, special methodological frameworks; Denk & Grimm, 2010; Simeone et al., 2018).

In the present study, we explored the applicability of the same experimental pathway in six geographic samples of the core group of Subgenus Fagus: the *F. crenata – F. sylvatica* s.l. lineage, a dominant element of temperate mesophytic forests of Eurasia. The two species involved were resolved as sister species in previous phylogenetic studies (Denk et al. 2005; Renner et al., 2016; but see Fig. 1); they share a number of morphological key features such as leaf-like cupule appendages and rhombic-crenate leaves, and their modern distribution is highly disjunct at the western and eastern margins of the Eurasian continent. Hence, they are ideal to investigate evolutionary and speciation processes in wind-pollinated forest-forming trees. Our objectives were to test the utility of 5S-IGS HTS data for delineating beech species, assessing intra- and inter-specific diversity, and gaining deeper insights into an evolutionary history that involved reticulation. Further, we discuss differences in speciation processes and success in wind- and insect-pollinated trees.

## Results

Pre-processing and removal of rare HTS reads (abundance < 4) yielded 145,643 sequences, corresponding to 4,693 representative 5S-IGS variants (work details, statistics and sequence structural features reported in Data S1, S2). Of these, 686 had an abundance ≥ 25 and were included in the phylogenetic analyses leaving 4007 variants to be classified using the Evolutionary Placement Algorithm (Data S2, S3). Analyses of 38 selected sequences further clarified the evolutionary relationships among the detected 5S-IGS lineages (Data S4).

### 686-tip 5S-IGS backbone phylogeny

We first categorized the 4,693 5S-IGS sequences into nine sequence classes based only on their sample distribution (Fig. 2). Variants were labelled as “specific” when exclusively found in one (or two, in case of *F. sylvatica* s.str.) sample of a specific taxon: “japonica”, “crenata”, “Iranian orientalis”, “Greek orientalis”, and “sylvatica”. In addition, we identified four “ambiguous” classes, i.e. sequences shared among different taxa.

**Fig. 2.**
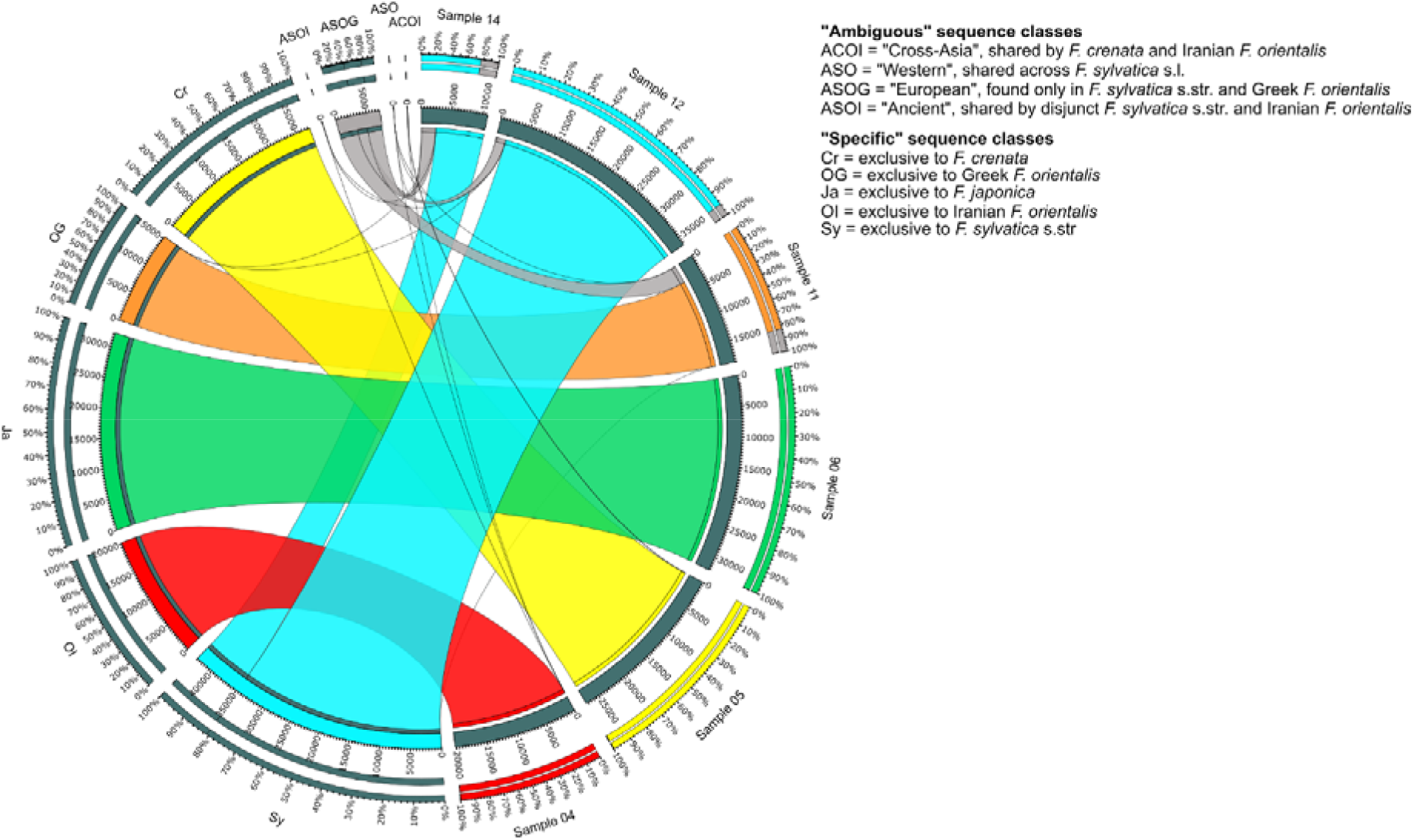
Map of 5S-IGS sequence class versus samples (DNA extraction pools). Class “Ambiguous” refers to unique sequence variants shared between samples of different taxonomic identity (Iranian and Greek *F. orientalis* treated as different taxa); class “Specific” refers to variants exclusively found in a single taxon sample (or two samples, in the case of *F. sylvatica* s.str.).

In a second step, five main 5S-IGS lineages (A, B, X, I, and O) were defined based on the phylogenetic analysis of 686 sequences with abundance ≥ 25 (Fig. 3).

**Fig. 3.**
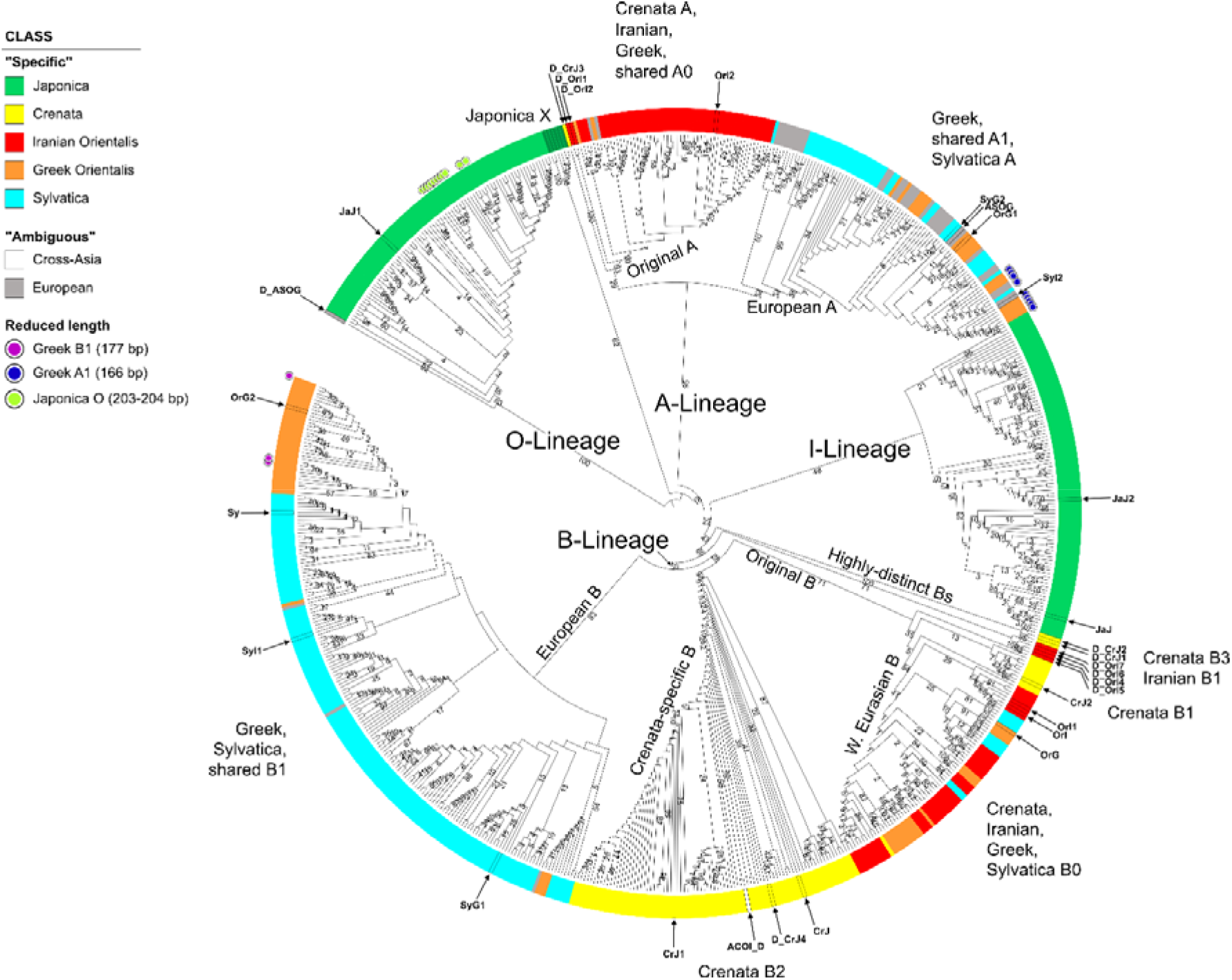
Circular cladogram based on maximum likelihood (ML) tree inference and bootstrap (BS) analysis (550 BS pseudoreplicates) of the 686-sequence matrix including all 5S-IGS variants with a total abundance ≥ 25. The tree is rooted on the genetically most distinct O-type lineage; numbers at branches give ML-BS support. Colours indicate sequence class based on their sample distribution (Fig. 2); very short variants and sequences selected for the 38-sequence matrix (cf Fig. 5) are highlighted.

The two ingroup 5S-IGS main types, ‘A-Lineage’ and ‘B-Lineage’, formed distinct clades (bootstrap [BS] support of 95/47) predating speciation within the *F. crenata* – *F. sylvatica* s.l. lineage (short: *crenata-sylvatica* lineage).

The Maximum Likelihood (ML) tree and Neighbour-Net (NNet) network (Fig. 4) show the duplicity of the five “specific” sequence classes. Most “ambiguous” variants are part of the ‘European A’ clade (BS = 34), while Iranian A-Lineage variants are exclusively found in the sister clade (‘Original A’; BS = 53, BS = 29 when including ‘Crenata A’ variant). The lower support for the B-root (Fig. 3) relates to the higher diversity in the B-Lineage (Fig. 4; Table 1), which comprises two genetically coherent but diverse clades (‘European B’ and ‘Crenata B2’) and a poorly sorted clade ‘Original B’. The ancestor of the *crenata-sylvatica* lineage was polymorphic and the modern pattern affected by incomplete lineage sorting. The Iranian *F. orientalis* individuals represent a now genetically isolated sub-sample of the original variation found in the western range of the *crenata-sylvatica* lineage. In contrast, the Greek *F. orientalis + F. sylvatica* s.str. and *F. crenata* are better sorted.

**Fig. 4.**
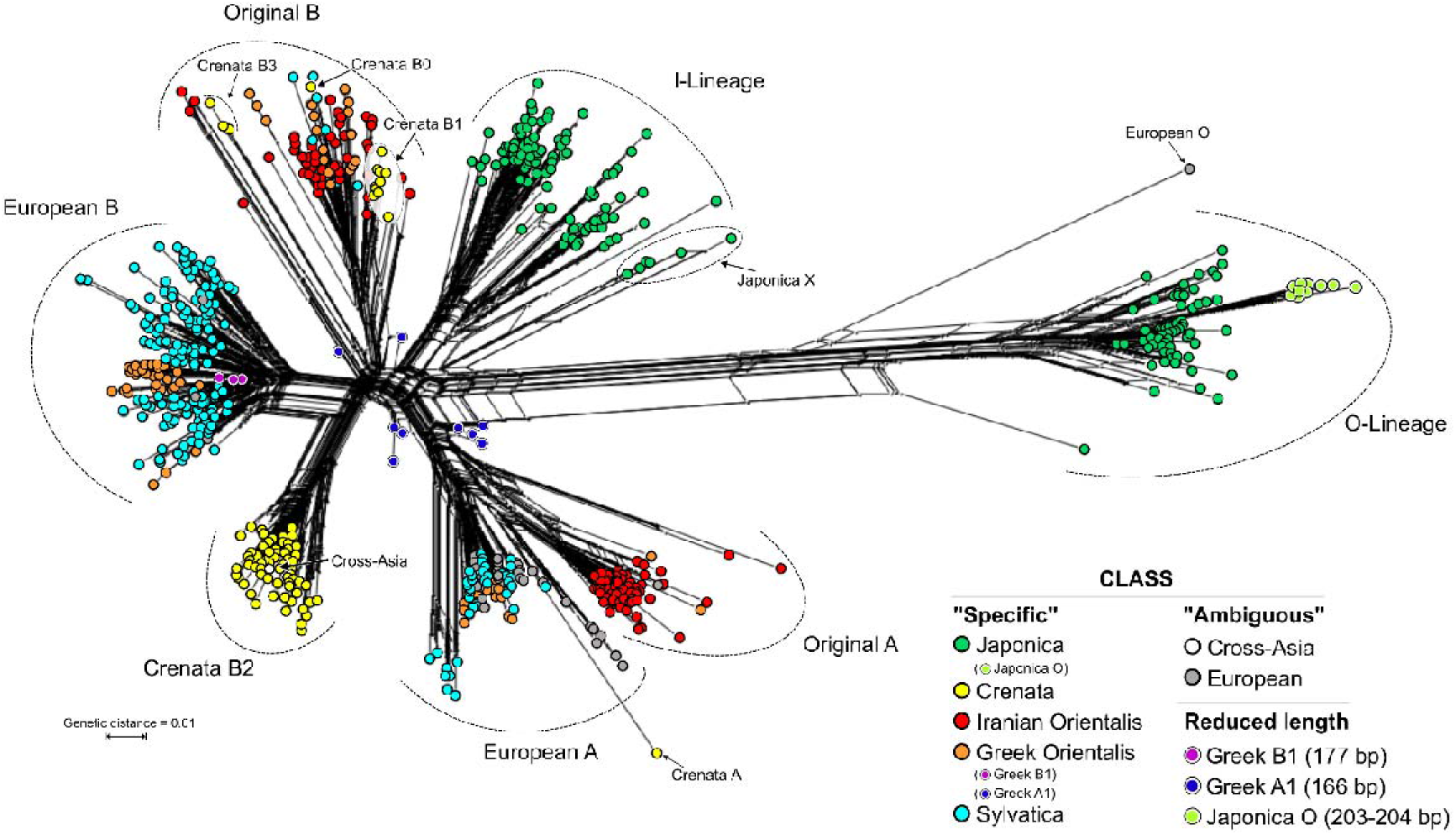
Neighbour-net for the 686-sequence matrix, inferred from uncorrected *(p-)* pairwise distances. Neighbourhoods defined by well-defined interior “trunks” relate to prominent sorting events (bottleneck situations; evolutionary jumps) leading to coherent 5S-IGS lineages with high (near-unambiguous) root branch support in Fig.3; “fans” represent poor sorting of more ancient 5S-IGS variants forming incoherent lineages with ambiguous root branch support. Note the absolute genetic distance between the assumed outgroup, ‘Japonica O’type, and the ingroup types including the ‘Japonica I’ type. Colouration as in Fig. 3.

**Table 1.**
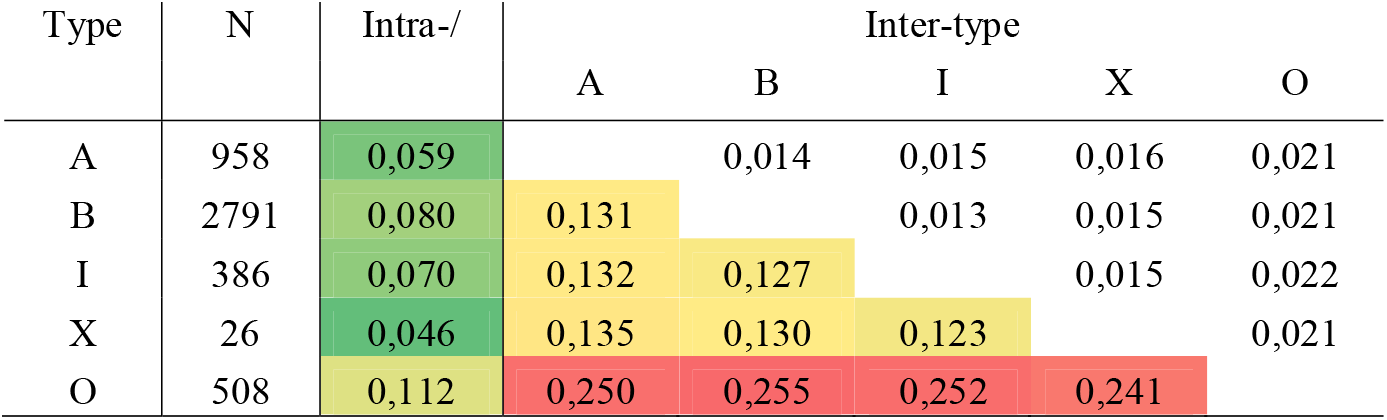
Mean intra- and inter-type 5S-IGS divergence estimates. (standard error shown above the diagonal). N = sample size (number of variants)

Our data do not show ongoing lineage mixing in Japan between *F. japonica* (Subgenus Engleriana) and *F. crenata* (Subgenus Fagus). However, one of the *F. japonica* types, ‘I-Lineage’, is substantially more similar to both ingroup main types (A- and B-Lineage) than the other dominant *F. japonica* type (‘O-Lineage’; Fig. 4). One ambiguous variant shared by western Eurasian beeches (‘European O’) is part of the O-Lineage; a small clade of *F. japonica-exclusive* variants (‘Japonica X’) nests between the A-B-I and the O-Lineage clades. *Fagus crenata* shares a relatively recent common origin with the western Eurasian beeches, represented by the ‘Original B’-type (BS = 71) with two subgroups. Its ‘Western Eurasian B’ subclade (BS = 54; including one *F. crenata* B-variant: ‘Crenata B0’) is poorly sorted. Potentially ancient variants closest to ‘Crenata B1’ sequences (grade in Fig. 3; neighbourhood in Fig. 4) persisted in Italian, Greek and Iranian populations. In contrast to its western Eurasian relatives, *Fagus crenata* lacks non-deteriorated A-Lineage variants (Table 2; Data S1, section 4.2) with most variants falling within a highly supported, *F. crenata*-exclusive B-type clade (‘Crenata B2’; BS = 98); this *F. crenata*-specific lineage also includes the ‘Cross-Asia’ shared variant found as a singleton in Iranian *F. orientalis.* The remainder (‘Crenata B3’) is placed between the I-Lineage and the core group of the B-lineage, together with sequentially highly derived Iranian B-type variants (‘Iranian B1’). One sequence (‘Crenata A’) is placed within the ‘Original A’ clade, as sister to all other variants (Fig. 3; possibly a tree-branching artefact, cf. Fig. 4).

**Table 2.** Main 5S-IGS sequence types observed and their evolutionary interpretation.

| Type | Taxonomic coverage | Evolutionary interpretation |
| --- | --- | --- |
| O | *F. japonica*, co-dominant, mostly inconspicuous<br>*Crenata-sylvatica* lineage, very rare, ± deteriorated <sup>a</sup> | Evidences ancient hybridization/ polyploidization event |
| I | *F. japonica*, co-dominant, inconspicuous | Reflecting shared ancestry with 'Subgenus *Fagus*' |
| A | *F. sylvatica* s.l., frequent, mostly inconspicuous<br>*F. crenata*, very rare, deteriorated | Shared ancestral type, most diverse in Iranian *orientalis* |
| B | *Crenata-sylvatica* lineage, frequent, mostly inconspicuous | Lineage-specific types, diverse and ± sorted |
| Original B<sup>b</sup> | Relatively common, rare in *F. sylvatica* s.str. | Reflecting divergence between East Asia and western Eurasia |
| Iranian B1 | Iranian *F. orientalis*, uncommon, mostly inconspicuous | Strongly diverged types, reflecting ancient genetic drift |
| Crenata B3 | *F. crenata*, uncommon, mostly inconspicuous |  |
| Crenata B2 | *F. crenata*, most frequent, *inconspicuous* | Species-specific types |
| European B | *F. sylvatica* s.str., Greek *F. orientalis*, *inconspicuous* |  |
| Relict lineage | Western Eurasia, very rare; ± deteriorated | Reminiscent of ancient speciation processes or reticulation |
| X | *F. japonica*, rare, inconspicuous | Unclear; potentially linked to non-sampled Chinese spp. of 'Subgenus *Fagus*' |

<sup>a</sup> Signs of sequence deterioration include increased AT-content, accumulation of aberrant sequence features, and rare additional deletions.
<sup>b</sup> Two subtypes: Crenata B1, Western Eurasian B (incl. Crenata B0)

The only other subtype showing the same level of genetic coherence as ‘Crenata B2’ is the ‘European B’ type (BS = 93; Fig. 4), shared by *F. sylvatica* s.str. and Greek *F. orientalis* (Table 2). Given its distinctness, the ‘European B’ type might reflect most-recent sorting and speciation events that involved *F. sylvatica* and western (Greek) *F. orientalis* but not their eastern (Iranian) relatives. In general, the western Eurasian beeches form a genetic continuum characterized by several, partially incomplete sorting events (‘Iranian A’ vs. ‘European A’; ‘Original B’ vs. ‘European B’). Within this continuum, Iranian *F. orientalis* appears to be most isolated and ancient/ ancestral with respect to *F. crenata* and the I-Lineage of *F. japonica.*

Very short ‘Greek orientalis’ variants are deeply embedded within the ‘European A’ and ‘European B’ clades; all short ‘Japonica’ variants are O-Lineage (Data S1, section 4.1). The distance-based NNet placed all short variants next to the centre of the graph (Fig. 4). Thus, they are sequentially undiagnostic lacking more than 100 bp from the 5’ or central part of the spacer but also inconspicuous within the larger ingroup (*crenata-sylvatica* A-, B- and *F. japonica* I-Lineage).

### Framework phylogeny

The ML trees for the 38 selected sequences representing the most abundant variants within each sample, strongly deviating variants, and “ambiguous” variants, show the deep split between outgroup (O-Lineage) and ingroup 5S-IGS variants (I-, A- and B-Lineage; Fig. 5). The mid-point root corresponds to this deep split, indicating an early (ancient) duplication of 5S rDNA arrays. Tip-pruning and elimination of the sequentially indiscriminative *F. japonica* X-Lineage (Figs 3, 4; for details see Data S1) led to a marked increase in backbone branch support: the divergence between *F. japonica* ingroup variants (I-Lineage) and B-Lineage clades occurred after the isolation of the A-Lineage (BS = 99/84). A group of sequentially unique, rare (abundances < 25), partly shared variants form a distinct clade, a sixth lineage. This “Relict Lineage” is placed between the outgroup (O-Lineage) and ingroup subtrees (A-B-I clade). Detailed analysis of sequence structure (Data S1, section 4.3) shows that this placement is only partly due to potential ingroup-outgroup long-branch attraction. Both subtrees (O- and Relict Lineage) include variants most different from the ingroup (A-B-I-X) consensus, and branch support is higher when length-polymorphic regions are included. However, these rare shared types also have an increased number of mutational patterns, which appear to be primitive within the entire A-B-I-X lineage. Thus, they may represent relict variants; ancestral copies still found in 5S rDNA arrays that were subsequently replaced and eliminated within the *crenata-sylvatica* lineage (Table 2). Another important observation is that the long-branching A- and B-Lineage variants found exclusively in the Iranian sample comprise strongly divergent, genetically coherent lineages and two *F. crenata-*unique variants with no signs of pseudogeny. While some rare, shared ingroup variants (Relict Lineage types, ‘Crenata A’, some ‘European O’; Fig. 4) show clear signs of sequence deterioration, others are inconspicuous (no pseudogenous mutations in flanking 5S rDNA) and can be highly similar to the most abundant variants.

**Fig. 5.**
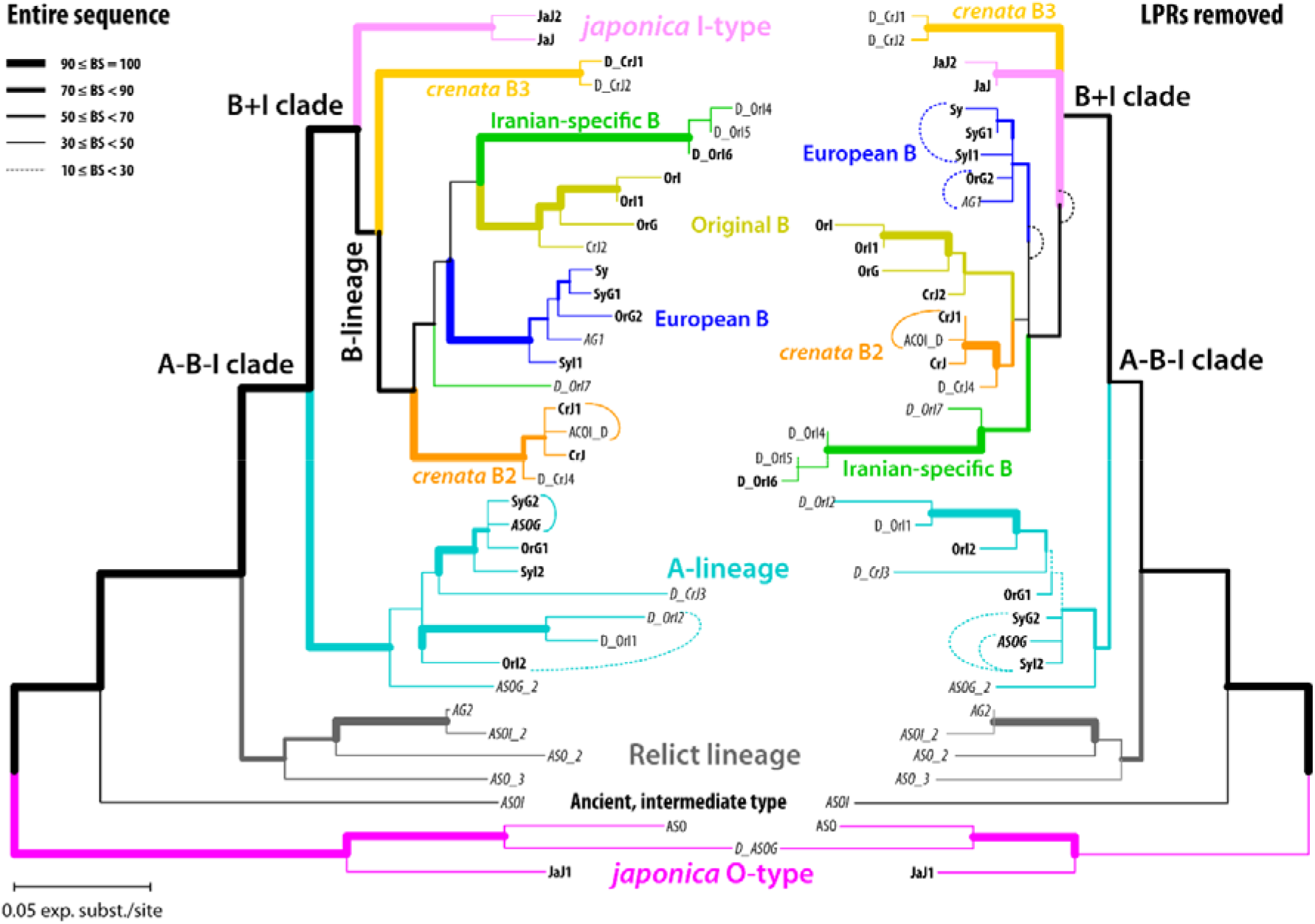
Maximum likelihood phylograms inferred from the selected 38-sequence matrix including the most common variants of each main lineage, and rare, shared variants (labelled as “*A*...”) as well as high-divergent variants *(“D_...* “). Left tree was inferred with generally length-polymorphic regions (LPR) included (as defined in Data S4, *Motives*); right tree with LPR excluded. Line thickness visualises non-parametric bootstrap (BS) support based on 10,000 BS pseudoreplicates.

### Duplicity and differences between 5S-IGS populations in each sample

The unrooted, single-sample ML trees recovered two 5S rDNA clusters (5S main types) with high bootstrap values (71–100) in all studied samples (Fig. 6). This is in accordance with previous cytological results of Ribeiro et al. (2011), who found two functional 5S loci in (diploid) *F. sylvatica* s.str.. The main splits separate I- and O-Lineage variants in the outgroup *F. japonica* and A- and B-Lineage variants in *F. sylvatica* s.l. The difference between the two clusters is most pronounced in *F. japonica;* the least intra-species (intra-sample) divergence is exhibited by *F. crenata,* which largely lacks A-Lineage 5S-IGS. Phylogenetically intermediate variants characterize the western Eurasian samples (long-branched in Iranian *F. orientalis*); strongly modified variants with little affinity to either 5S rDNA cluster, hence, connected to the centre of the graph, are abundant in *F. crenata* (two lineages, one representing a relict ‘Crenata A’-type; see Fig. 4) and *F. sylvatica* s.str. (a single lineage in each population). These intermediate sequences do not show any structural peculiarity, except for the ‘Crenata A’-type variant with a reduced GC content (34.3%; Data S1, section 4.2 and appendix B). The outgroup-type ‘European O’ represents the longest branches in the Greek *F. orientalis* (Greece) and both *F. sylvatica* s.str. samples (Fig. 6, samples 11, 12, 14). As a trend, the variants within the B- and I-Lineage show a higher divergence and phylogenetic structure than found within the A- and O-Lineage subtrees.

**Fig. 6.**
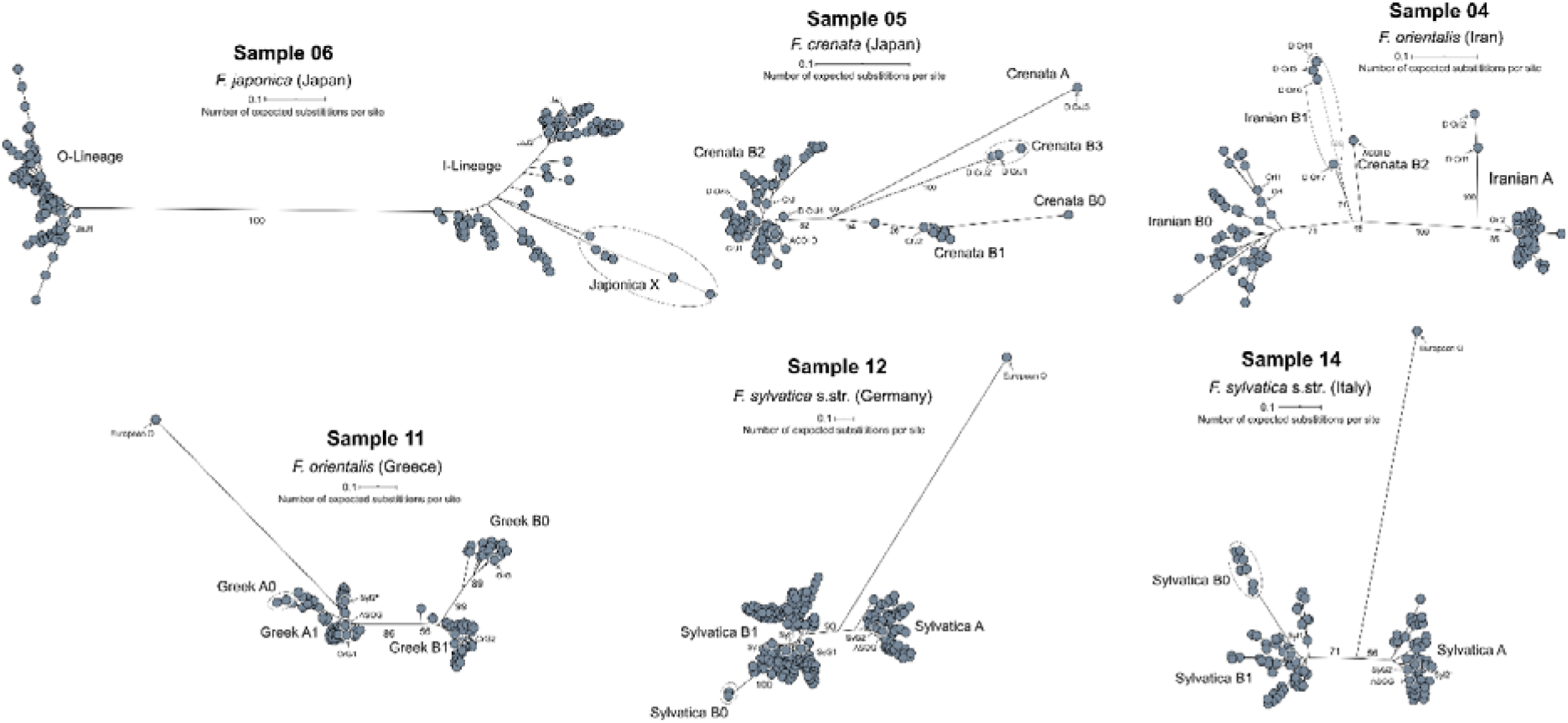
Sample-wise maximum likelihood (ML) trees illustrating the bimodality of the 5S-IGS pool in each sample. Numbers give ML bootstrap support (based on 1000 pseudoreplicates) for selected major phylogenetic splits. Subtree and individual labels refer to lineages and tips introduced in Figs 3–5.

The Evolutionary Placement Algorithm (EPA) assessed the phylogenetic affinity of all variants with an abundance ≥ 4 not included in the 686-tip matrix (Data S2, S3). The A-/O- vs. B-/I-Lineage variants differ not only in GC content and sequence length but also relative abundance (Fig. 7). In the *crenata-sylvatica* lineage, the GC-richer B-Lineage variants are more frequent, while the GC-poorer A-Lineage sequences make up a higher portion of the HTS reads corresponding to rare sequence variants. The B-Lineage successively replaces the A-Lineage along a (south-)east (Iranian *F. orientalis)* to (north-)west (German *F. sylvatica)* gradient. In *F. japonica,* the ratio of I-Lineage, matching the B-Lineage in GC-content and length, to O-Lineage (GC-richest and longest variants) is ~2:1. In general, 5S-IGS arrays showed a high capacity to conserve signal from past reticulations and deep divergences: types placed by EPA in ‘European O’ clade, representing a remnant, deteriorating sister lineage of the *F. japonica* O-Lineage (cf. Table 2; Figs 3, 4), can be found in all samples of the *crenata-sylvatica* lineage (Data S1, appendix A). In contrast, X-Lineage variants are exclusive to *F. japonica*. The single, low-abundant I-Lineage sequence identified by EPA in the *F. sylvatica* s.str. sample from Germany represents a relict variant from the initial radiation within the (A-)I-B clade. The Relict Lineage (cf. Fig. 5) is represented in all samples of the *crenata-sylvatica* lineage as well. Despite showing signs of beginning pseudogeny in the flanking 5S rRNA gene regions, its GC contents (35.1–40.0%) range between the median values of the O-Lineage and the I-B clade. Hence, they match the range seen in the A-Lineage, and are of the same length as most A-and B-Lineage types (Appendix B in Data S1).

**Fig. 7.**
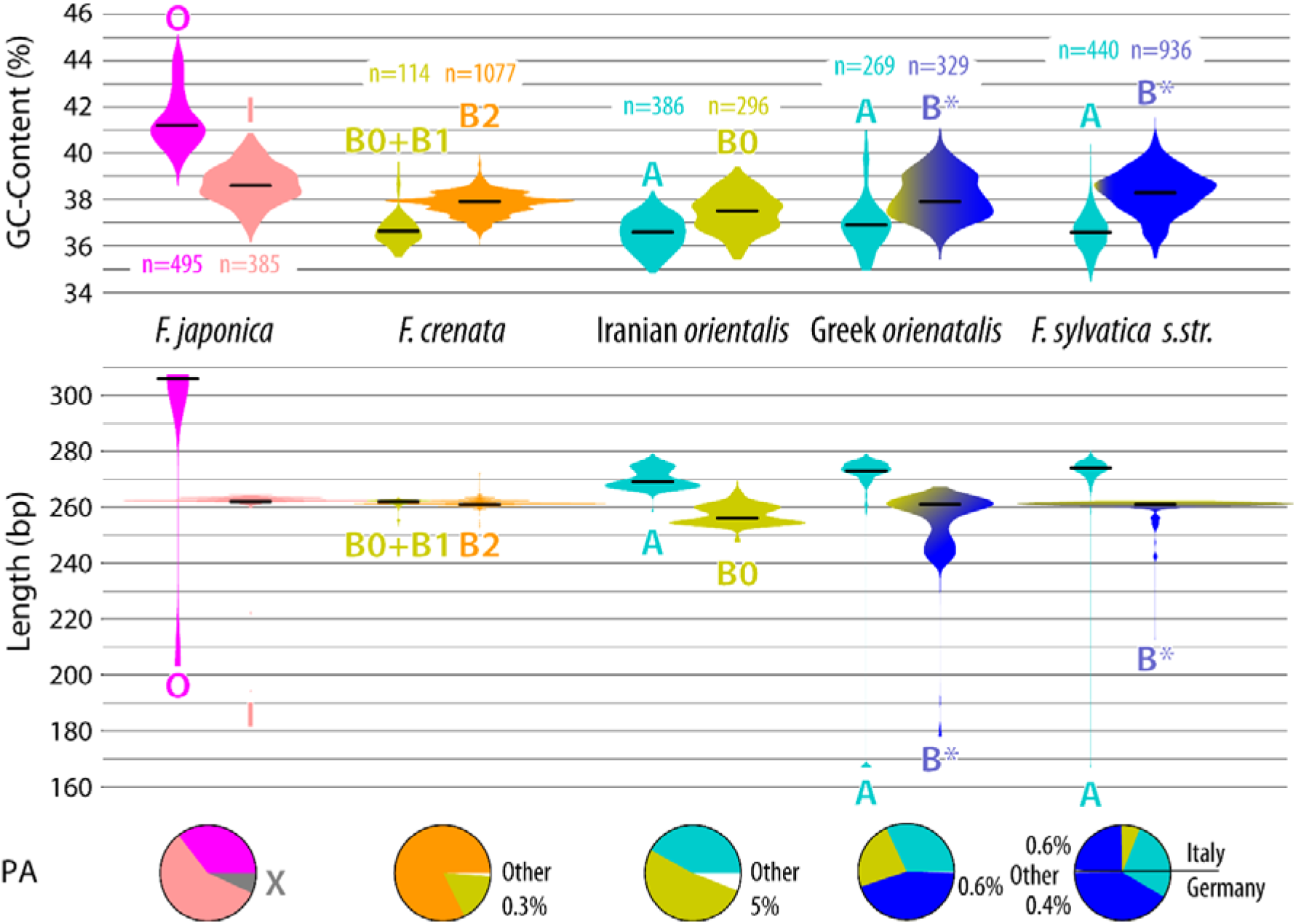
Amplicon GC content and length violin plots for the (co-)dominant lineages/main 5S-IGS types found in each sample. Horizontal black lines give the median value for each main type (based on all variants of the respective lineage). Width of violin plots adjusted to visualise the relative proportion (number of HTS reads) of each type within a sample; *n* gives the plots’ sample number (number of distinct variants). Pie-charts give the proportional abundance (PA) of the plotted types within each sample (see Data S1, appendix A, for absolute numbers). “O”, “I”, “A”, “B0”–”B3” refer to respective (sub)types/ lineages labelled in the 686- and 38-guide trees (Figs 3, 5). Colouration gives affinity to main 5S-IGS lineages (Figs 3–5). *, excluding very rare Iranian *F. orientalis-* and *F. crenata-*specific B types (cf. Fig. 8).

## Discussion

### 5S-IGS data to study complex history of wind-pollinated trees

Fagaceae have shifted at least twice from insect-pollination to wind-pollination during their evolutionary history leading to the rise to dominance of temperate tree genera such as *Fagus* and *Quercus.* These shifts left imprints in the genetic signature of the family recording ancient and recent gene flow between lineages that diverged tens of millions of years ago (e.g. McVay et al., 2017; López-Heredia et al., 2020). These characteristics may have contributed to the success story of these trees, but at the same time complicated phylogenetic reconstructions. The underlying processes can only be represented with a species network; any species tree (combined or coalescent, Jiang et al., 2021, or phylogenomic, Yang et al., 2020, using complete plastomes) will be incomprehensive, irrespective of the obtained support (Sections 4.5, 4.6 in Data S1). *Fagus* species are not only wind-pollinated but also animal-dispersed and have a narrow ecological niche (Peters, 1997). Therefore, beech populations are at risk of becoming isolated during phases of area disruption, which can lead to speciation accompanied by lineage sorting and increased genetic drift. At the same time, species boundaries remain permeable as expected in wind-pollinated plants. During phases of area expansion secondary contacts and lineage mixing occurred. In Subgenus Engleriana, up to three sequentially divergent ITS variants indicated (past) inter-species gene flow; likewise, *F. longipetiolata* and *F. pashanica* shared two distinct ITS lineages (Denk et al., 2005). Recently, the disparate gene trajectories seen in the polymorphic nuclear gene data of Jiang et al. (2021; Section 4.5 in Data S1; Data S5) provided further evidence for lineage mixing. Jiang et al.’s (2021) data indicate past, west-east introgression via the North Atlantic land bridge (cf. Grímsson & Denk, 2007) and suggest multiple secondary contacts shaping the gene pools of modern East Asian species of subgenus Fagus, most pronounced in the species pair *F. longipetiolata/ F. lucida* (Section 4.6 in Data S1). Hence, the objective of our study was to recruit data sets reflecting ancient, between ancestors of modern species, and recent, between modern species, reticulations.

The permeability of species boundaries sets apart wind- and insect-pollinated, common tree genera. A recent study on genetic consequences of pollination mode (Wessinger, 2021) showed that wind pollination leads to reduced population genetic structure. For example, in one insect- and one wind-pollinated species of *Chamaedorea* (Arecaceae) in Mexico, the wind-pollinated species showed less isolation by distance as compared to the insect-pollinated species; the same was true in two common shrubs co-occurring in Mediterranean scrub, *Pistacia lentiscus* (wind-pollinated) and *Myrtus communis* (insect-pollinated). In addition, insect-pollinated, woody angiosperms with limited seed dispersal abundant in humid temperate regions such as *Acer* and *Rhododendron* are genetically coherent down to specieslevel (Grimm et al., 2006, 2007; Chappell et al., 2008; Grimm & Denk, 2014). Pairwise niche comparisons in temperate species of *Acer* have revealed only infrequent niche overlap continentally and globally (Grossman, 2021). In contrast, in the wind-pollinated *Fagus* displays neither niche specificity nor correlation with population sizes and occupied area (Cai et al., 2021). Genetic distances between sections of insect-pollinated *Acer* resemble those of subgenera or genera in wind-pollinated Fagaceae irrespective of the used marker (cf. various sequence data stored in gene banks). In some species (e.g. *Acer ibericum;* Grimm et al. 2007), intra-individual ITS variability can reach similar levels as found in individuals of *Fagus* (and exceeding variation in *Quercus)* but this variation is not shared among morphologically and (otherwise) genetically distinct species. Thus, in species of *Acer* both ITS and 5S-IGS variants, as far as studied, are highly species-diagnostic and sorted to a degree that allows the recognition even of cryptic species (Caucasian *A. orthocampestre,* Grimm & Denk, 2014) within a similar geographic range as western Eurasian *Fagus*. Furthermore, ITS and 5S-IGS data of *Fagus* converge to relationships incongruent with single-/low-copy nuclear genes (cf. Fig. 1; Data S5). In contrast, in *Acer,* the most recent nuclear-phylogenomic species tree (Li et al., 2019, using a 500-gene dataset) is largely congruent with earlier phylogenies using ITS/5S-IGS data/polymorphism. Likewise, species diversification rates in *Rhododendron* are more than ten times higher than in *Fagus* (Renner et al., 2016; Shrestha et al., 2018). The rise to dominance in temperate and boreal forest ecosystems of wind-pollinated Fagales taxa such as *Betula, Fagus,* and *Quercus,* appears to be linked with high within-population gene flow and permeable species boundaries. Consequently, the evolutionary history of such tree genera is different, more complex, and much more difficult to characterize than in insect-pollinated woody angiosperms.

The 5S intergenic spacers are a unique source of information to investigate complex evolutionary processes in wind-pollinated, temperate woody angiosperms. They are the only currently known region that (*i*) is high-divergent and multi-copy, (*ii*) can be easily amplified due to the conservation of the 5S rRNA genes, and (*iii*) can be analysed using High-Throughput Sequencing because of its relatively short length. In this work, the O-Lineage and I/X-Lineage variants found in *F. japonica* indicate hybrid (allopolyploid) origin, and the distribution of A- and B-lineage variants in the *crenata-sylvatica* lineage reflect speciation processes predating the formation of the modern species. Hence, the 5S-IGS captures phylogenetic signal from 55 million years and multiple ancient hybridization events that are not (readily) traceable by other molecular markers (Oh et al., 2016; Yang et al., 2020; Jiang et al., 2021, but see Data S5).

### Inter-species relationships and status of Iranian *F. orientalis*

Our data confirm the close relationship of *F. crenata* with *F. sylvatica* s.l. (including shared identical or highly similar 5S-IGS variants) and the deep split between the two Japanese species representing two subgeneric lineages (Shen 1992; Denk et al., 2005; Renner et al., 2016). The latter has recently been confirmed using single-/low-copy nuclear genes (Jiang et al., 2021). Jiang et al. (2021) suggested that East Asian species form a monophyletic group, setting apart the Japanese *F. crenata* and the western Eurasian *F. sylvatica* s.l. (Fig. 1). Not addressed in the original paper, Jiang et al.’s (2021) gene sample reveals a complex pattern of inter- and intra-species relationships in the East Asian species, partly involving western Eurasian and North American species (Data S5). While three genes (P21, P28, and P54) support an East Asian clade with BS values ≥ 53, nine of the 28 genes reject it. Instead, they support incompatible (conflicting) splits with BS values ranging from 61 (gene P38) to 99.7 (gene F138). The East Asian clade gets its support from the distinctness of western Eurasian and North American species and inconsistently shared, primitive or slightly evolved, genotypes (Fig. S18 in Data S1). Individuals carrying ‘alien’ gene variants (genes P4, P21, P49, P52, P69, and P72) furthermore point to pre- (ancient) or post-speciation (secondary) gene flow between *F. crenata* and *F. lucida* or *F. longipetiolata, F. hayatae* or *F. longipetiolata* and *F. lucida,* and Chinese *F. hayatae (F. pashanica)* and *F. longipetiolata.* Therefore, additional studies are needed to assess to which degree *F. crenata* and *F. japonica* 5S-IGS gene pools are impacted by primary or secondary contact with the predecessors of *F. hayatae, F. longipetiolata* and *F. lucida.* The mutation patterns seen in Jiang et al.’s (2021) data confirm, nonetheless, that the western Eurasian species *(F. sylvatica* s.l.) evolved from an ancestor genetically close to the present-day *F. crenata.* Although not resolved as sister clades in trees based on single-/low-copy nuclear data (Oh et al., 2016; Jiang et al., 2021; but see Section 4.6 in Data S1), *F. crenata* and *F. sylvatica* s.l. are likely sister species.

Regarding the western Eurasian species, our results corroborate previous morphological (Denk, 1999a, b) and population-scale isoenzyme and genetic studies (Gömöry & Paule, 2010; Bijarpasi et al., 2020), which identified a split between disjunct populations traditionally considered as *F. orientalis* in Europe and adjacent Asia Minor, Iran, and the Caucasus. Furthermore, our data confirm the relatively recent contact and mixing between the westernmost *F. orientalis* (here represented by a Greek population) and *F. sylvatica* s.str. (Papageorgiou et al., 2008; Müller et al., 2019).

The Iranian *F. orientalis* must have been isolated for a much longer time and clearly represents a distinct species. According to Denk (1999b), Gömöry & Paule (2010) and Gömöry et al. (2018), the Iranian populations are morphologically and genetically distinct from both the western *F. orientalis* (SE. Bulgarian and NW. Turkish populations) and the Caucasian *F. orientalis*. The high amount of unique 5S-IGS variants can be explained by the geographic isolation and restriction of Iranian beech populations. However, they are not genetically impoverished (see also Erichsen et al., 2018). The overall diversity of 5S-IGS A- and B-Lineage variants points to a complex history of Iranian beech within the limits of its current distribution, characterized by strong topographic heterogeneity (sea-level to close to 3,000 m a.s.l.) and pronounced small and large-scale climatic gradients (Sagheb-Talebi et al., 2014). One possible further source are the Caucasian populations geographically closer to the Iranian, which may have been in contact in the more recent past (via Cis-Caucasia and Azerbaijan). Hence, some of the Iran-specific variants in our study may be shared with Caucasian populations. Additional data from the Caucasus and north-eastern Turkey are needed to quantify the degree of this contact and to clarify the taxonomic status of these populations.

Other genetic imprints in the Iranian populations might be a legacy from extinct beech populations growing to the east and possibly from the disjunct populations of the Nur (Amanos) Mts in south-central Turkey. The fact that *F. crenata* and Iranian *F. orientalis* still carry exclusively shared (‘Cross-Asia Crenata B2’) and related (‘Crenata A’; ‘Crenata B0’) 5S-IGS variants (Figs 3, 4) fits with a more or less continuous range of beech populations from the southern Ural, via Kazakhstan and Siberia, to the Far East until the end of the middle Miocene (see *Formation of 5S-IGS gene pools).*

### Ancient polyploidization and past reticulation in beech

The *Fagus crenata-sylvatica* s.l. lineage represents the most widespread, recently diverged and diversified branch of the genus (Data S4; Denk & Grimm, 2009; Renner et al., 2016). Differentiation patterns observed in the 5S-IGS provide a unique temporal window into the evolution of a wind-pollinated tree genus (Fig. 8): speciation and isolation led to the accumulation of new lineage-specific variants, which were then exchanged or propagated during episodes of favourable climate and massive expansion of beech forests.

**Fig. 8.**
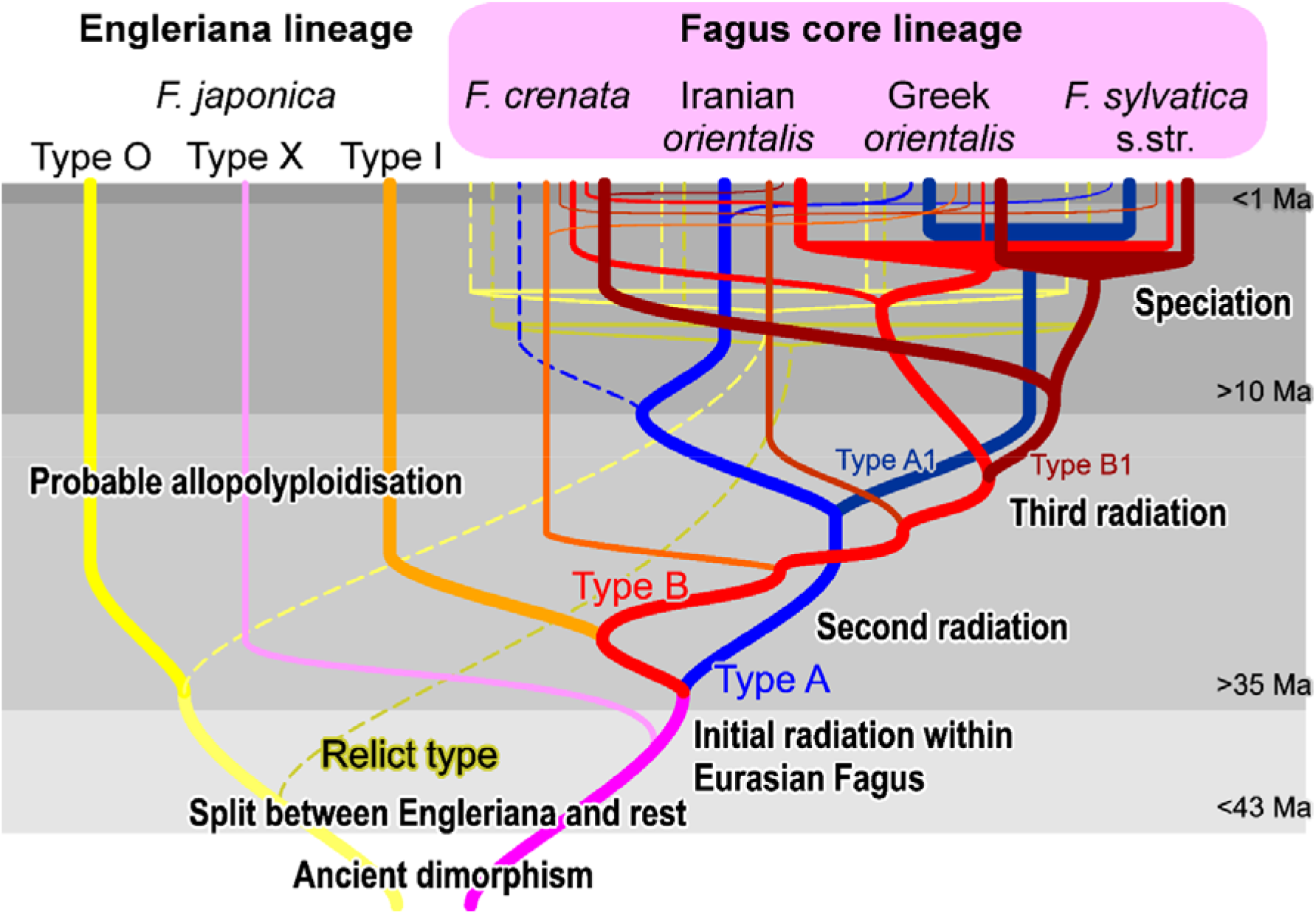
Doodle summarising the totality of our results and their interpretation regarding all available (referenced in the text) information.

New 5S-IGS variants would have been easily dispersed via pollen over long distances, while the genetic characteristics of the mother trees (passed on via ovules) remained geographically constrained. The main dispersal vector for *Fagus* seeds have been sedentary jay birds (Darley-Hill & Johnson, 1981), a group of corvids that evolved in the middle Miocene (Mayr, 2005) coinciding with the highest taxonomic diversity recorded in *Fagus* (Denk & Grimm, 2009). Another important animal seed-dispersal vector are small rodents, with specialized tree squirrels occurring since the early Oligocene (Steppan et al., 2004; Casanovas-Vilar et al., 2018); a time when first major beech radiations took place (Denk & Grimm, 2009; Pavlyutkin et al., 2014).

Additional data from Chinese and Taiwanese species will be needed to assess whether the A-or B-Lineage, or the partly pseudogenic Relict Lineage, represent the original stock of western Eurasian beeches and their Japanese sister species. The A-Lineage is largely lost in *F. crenata* and sequentially equally distant to the B-Lineage and the *F. japonica* ingroup variants (I-Lineage; Fig. 4). Therefore, we can expect that both lineages, A and B+I (and possibly the X-Lineages) are present at least in some of the Chinese species and represent an ancient polymorphism shared by all Eurasian Subgenus Fagus species (Fig. 8; cf. Denk et al., 2005, for ITS). Likewise, our assessment of the 28-gene data of Jiang et al. (2021; Data S5) revealed poor genetic isolation of *F*. *crenata* from all other East Asian species of subgenus Fagus; confirmed by upcoming, densely sampled complete plastome data focussing on Japanese beech (first results accessible on ResearchGate: Worth et al., 2021). This is consistent with a main finding of the present study that *Fagus crenata* and its precursor(s) exchanged genetic material with other East Asian beeches when getting into contact.

The pronounced ancient nuclear polymorphism, also a predominant feature in Jiang et al.’s (2021) data, implies that modern beeches are of hybrid or allopolyploid origin. Our 5S-IGS data clearly indicate a hybrid origin for the species of Subgenus Engleriana, with the *F. japonica* I-Lineage variants representing common ancestry with Subgenus Fagus (Eurasian clade), while the outgroup variant (O-Lineage) represents another, potentially extinct, lineage of high-latitude beeches (Denk & Grimm, 2009; see also Fradkina et al., 2005; Grímsson et al., 2016). Hybrid/ allopolyploid origins would also explain the great divergence observed in the ITS region of *Fagus*, especially Subgenus Engleriana (Denk et al., 2005), a lineage with a poorly sorted nucleome and still sharing ± primitive genotypes with North American and East Asian members of Subgenus Fagus (Data S5). This hypothesis receives strong support from fossil evidence. Oldest unambiguous fossils of *Fagus,* dispersed pollen grains from the early Paleocene of Greenland, of small size and with narrow colpi reaching almost to the poles (Grímsson et al., 2016), resemble Subgenus Engleriana. The subsequent radiation of beeches involved western North America and East Asia, where fossil-taxa combining morphological characteristics of both modern subgenera are known (Denk & Grimm, 2009). For example, the early Eocene *F. langevinii* has branched cupule appendages as exclusively found in extant *F. engleriana* along with Subgenus Engleriana type pollen but resembles Subgenus Fagus in features of leaf and nut morphology (Manchester & Dillhoff, 2005). Dispersed pollen from the late Eocene of South China also resembles modern pollen of Subgenus Engleriana (Hoffman et al., 2019). This might reflect an early phase in the evolution of *Fagus,* during which the modern subgenera were evolving, but not fully differentiated. Consequently, ancient genetic links can be traced in Jiang et al.’s (2021) data as well (Fig. S18, Tables S3, S4 in Data S1). By the early Oligocene (34 Ma), fossil-species can clearly be assigned to either Subgenus Fagus or Engleriana. *Fagus pacifica* from western North America resembles Subgenus Fagus in leaf architecture and the cupule/nut complex (Meyer & Manchester, 1997); cupules and leaves from the Russian Far East can unambiguously be ascribed to Subgenus Engleriana (Pavlyutkin et al., 2014; as *F. evenensis*, *F. palaeojaponica*, *Fagus* sp. 3).

Past reticulation is conceivable for the European species as well. The first beeches arrived in Europe in the Oligocene showing a general morphotype found across continental Eurasia (*F. castaneifolia*; Denk, 2004; Denk & Grimm, 2009). During the Miocene, *F. castaneifolia* was gradually replaced by *F. haidingeri*, the ancestor and precursor of all contemporary western Eurasian beeches (Denk et al., 2005; Denk & Grimm, 2009). Like *F. castaneifolia, F. haidingeri* shows high morphological plasticity. Hence, this fossil-species may represent a species complex rather than a single species. In addition, in southern Europe, gene flow between *F. haidingeri* and another fossil-species, *F. gussonii*, might have occurred. Jiang et al.’s (2021) genes F289, P14, P21, P52, and P54 document past, unilateral introgression via the now unavailable North Atlantic land bridge (Grímsson & Denk, 2007) from the North American into the western Eurasian lineage (Section 4.6 in Data S1). Our data confirm the potential for (sub)recent reticulation and incomplete lineage sorting between and within the morphologically distinct Greek *F. orientalis* and *F. sylvatica* s.str., and similar processes likely occurred repeatedly since the Miocene. Notably, the Greek *orientalis* and *F. sylvatica* s.str. comprise three main 5S-IGS lineages compared to only two in the Iranian sample. In addition to the inherited polymorphism shared with the Iranian *F. orientalis* (relatively similar ‘Original A’ and ‘European A’ types within the A-Lineage; shared ‘Western Eurasian B’), we found a highly abundant, moderately evolved but less diverse B-Lineage variant (European B in Figs 3–6), exclusive to Greek *orientalis* and *F. sylvatica* s.str., likely reflecting the recent common origin of these species.

### Spatio-temporal framework for gene pool formation in the *F. crenata - F. sylvatica* s.str. lineage

With currently available methods, it is impossible to date potentially paralogous-homoeologous multi-copy 5S IGS data. Nevertheless, the 5S IGS captured complex evolutionary signals, likely covering the past 55 million years. During the Eocene (56–33.9 Ma; Cohen et al., 2013, updated), the lineages leading to *F. japonica* and *F. crenata + F. sylvatica* s.str. started to diverge and speciation in the *crenata-sylvatica* lineage probably started in the late Oligocene (Chattian; 27.82–23.03 Ma; see above; Renner et al., 2016). The fossil record indicates that gene flow between the western(-most) and eastern(-most) populations of the *crenata-sylvatica* lineage became impossible after the middle Miocene, when the near-continuous northern Eurasian distribution of beech forests became fragmented at around 15–10 Ma (Denk, 2004; Arkhipov et al., 2005). Because of their geographic closeness, eastern (NE./E. Asian) populations of the *crenata-sylvatica* lineage remained in closer and longer contact with the south-eastern populations from which the modern Chinese and Taiwanese species evolved. Therefore, *F. crenata* may still be genetically closer to *F. lucida* (with possible Central Asian origins, Denk & Grimm, 2009) than *F. sylvatica* s.l. and the most southern species *F. hayatae* and *F. longipetiolata* (see Shen, 1992, for distribution maps). This contact is also reflected in the single-/low-copy genes used by Jiang et al. (2021; cf. Data S5, discussed in Section 4.6 in Data S1) but not in *LEAFY* intron data (Oh et al., 2016; Renner et al., 2016; cf. Fig. 1). Jiang et al.’s (2021) inferred ages for the diversification of Subgenus Fagus (c. 10 Ma) and speciation of *F. crenata* are, however, much too young and at odds with the rich and extensively studied fossil record of *Fagus* (cf. Denk & Grimm, 2009; Grímsson et al., 2016). First, in their dating of the multi-species coalescent (MSC), Jiang et al. (2021) assigned c. 30 and c. 20 myrs too young stem and crown ages for *Fagus*; likewise they assumed a much too young stem age for the outgroup comprising extra-tropical Castaneoideae and members of both oak subgenera (subgen. *Quercus* and *Cerris*). Jiang et al.’s (2021) inferred *Quercus* crown age is at least 25 myrs too young (cf. Hubert et al., 2014; Hipp et al., 2020). The various reticulation events between East Asian species of Subgenus Fagus indicated by Jiang et al.’s (2021) data, may lead to wrong and/or too young MSC estimates unless one infers a MSC network (see simulations in the supplementary information to Wen & Nakhleh, 2018). In addition, asymmetric introgression (a process not modelled by the birth-death model), for instance via long-distance pollen dispersal (e.g. Sofiev et al., 2013), may have quickly homogenized ancestral single-/low-copy gene polymorphism.

Gömöry et al. (2018) used allozymes and approximate Bayesian computation to examine the demographic history of western Eurasian beeches and suggested that *F. sylvatica* s.str. diverged from *F. orientalis* at ~1.2 Ma in the Pleistocene (Calabrian); this scenario is supported by the leaf fossil record (e.g. Follieri, 1958; Denk et al., in press). The diversity seen in Iranian *F. orientalis* (Figs 3, 5) may well represent the original 5S-IGS diversity within the western range of the Oligocene-Miocene precursor of the *crenata-sylvatica* lineage. The poorly sorted ‘Western Eurasian B’ lineage reflects the common origin, the originally shared gene pool, of the western Eurasian beeches and the Japanese *F. crenata* and ongoing gene flow in the Miocene (relict ‘Crenata B0’; ‘Crenata B1’ grade; Fig. 3). Furthermore, *F. crenata,* as the easternmost species, seems to have replaced its A-Lineage 5S-IGS variants with specific ‘Crenata type B2’ sequences. Originally, *Fagus* had a (near)continuous range from Japan via central Asia to Europe, and it is therefore possible that within this continuous area (extinct) *Fagus* populations acquired 5S-IGS variants that survived within the 5S gene pool of populations like those observed in Iran. Beech forests persisted throughout the entire Neogene in Iran (and in the Caucasus as well) although experiencing severe bottlenecks (Shatilova et. al., 2011; Dagtekin et al., 2020). The sharing of rare variants is consistent with this scenario as they link isolated populations and otherwise distinct species to a once common gene pool (e.g. Relict Lineage, ‘European O’, ‘Cross-Asia’ in ‘Crenata B2’; Figs 3–5). These variants are not part of the terminal subtrees, but instead reflect ancient, largely lost diversity that predates the formation of the modern species and possibly even the final split between Subgenus Engleriana (represented by *F. japonica*) and Subgenus Fagus *(crenata-sylvatica* lineage). 5S-IGS data from the North American species are needed in this context: do they have high-divergent, unique variants (as in non-pseudogenic ITS copies) or do they share lineages with certain Eurasian species (as in certain single-/low-copy genes)? Denk et al. (2002) reported for instance a pseudogenous ITS clone from a Georgian (Transcaucasus) individual sharing mutation patterns otherwise exclusive and diagnostic for the North American *F. grandifolia.*

In Europe, the Pleistocene fluctuations and Holocene expansion could have triggered the secondary sorting and homogenization of both A- and B-Lineage 5S-IGS arrays: ‘European A’ and ‘Western Eurasian B’ clades include relatively high proportions of shared (“ambiguous”) 5S-IGS variants in contrast to the highly coherent ‘European B’ clade (Figs 3, 4). Unhindered gene flow lasted much longer between Greek *F. orientalis* and *F. sylvatica* s.str. than between the latter and the more disjunct Iranian *F. orientalis* and/or Japanese *F. crenata*. A side effect of the higher genetic exchange between the western populations, a putatively larger active population size and more dynamic history, is the retention of ancient 5S-IGS variants that are likely relicts from a past diversification.

### Future research directions: From species trees to species networks

The rapidly increasing amount of genomic data demonstrate hybridization, reticulation in general, is the rule rather than an exception in plants (Yakimowski & Rieseberg, 2014). Genomes of numerous modern angiosperm lineages have preserved signals of multiple rounds of whole genome duplications; plant genome evolution includes complex processes of chromosome fractionation and rearrangement (Nieto Feliner et al., 2020). High-Throughput Sequencing opens up new avenues for investigating ancient hybridization/reticulation and the integration of distinct parental genomic traits into progenies (e.g. Ma et al., 2019; Cruz-Nicolás et al., 2021).

The 5S intergenic spacers provide a unique source of information to investigate these multifaceted evolutionary processes (see also Grimm & Denk, 2010; Volkov et al., 2017; Garcia et al., 2020; Vozárová et al., 2021). Despite our limited sample, our HTS data captured phylogenetic signals covering 55 million years of ancient and recent reticulation forming the gene pools of modern beeches. The O-Lineage and I/X-Lineage variants found in *Fagus japonica* signify hybrid (allopolyploid) origin, and the distribution of A-and B-lineage variants in the *crenata-sylvatica* lineage reflect speciation processes predating the establishment of modern species. In stark contrast to the standard dichotomous speciation model, modelled by phylogenetic trees, beech species perpetually form and fuse. Such reticulate speciation may be the norm in wind-pollinated Fagales *(Juglans:* Zhang et al., 2019; *Quercus:* Simeone et al., 2016; McVay et al., 2017; Hipp et al., 2020; *Nothofagus:* Premoli et al., 2012; Acosta & Premoli, 2018). The shift from insect-to wind-pollination revolutionized their speciation processes, maximizing the advantages gained from hybridization and introgression. Disentangling the key aspects of (ancient) hybrids’ genome architecture, discerning hybridization and introgression from incomplete lineage sorting to reconstruct the multi-species coalescent network (Wen & Nakhleh, 2018), should be the ultimate goal of future research (Mallet, 2016; Martin & Jiggins, 2017).

## Experimental procedure

### Plant material and HTS methodology

Figure 9 shows our sampling, experimental design, and workflow. Thirty beech individuals from three species (four recognized taxa) were collected in the wild, covering 14 total populations. Four samples contained five individuals each of *Fagus sylvatica* s.l. from single populations in Germany, Italy *(F. sylvatica* s.str.), Greece, and Iran *(F. orientalis);* individual DNA extracts from five populations of *F. crenata* from Japan (Subgenus Fagus) and *F. japonica* (Subgenus Engleriana; used as outgroup) were pooled as additional samples (details provided in Data S1, S2).

**Fig. 9.**
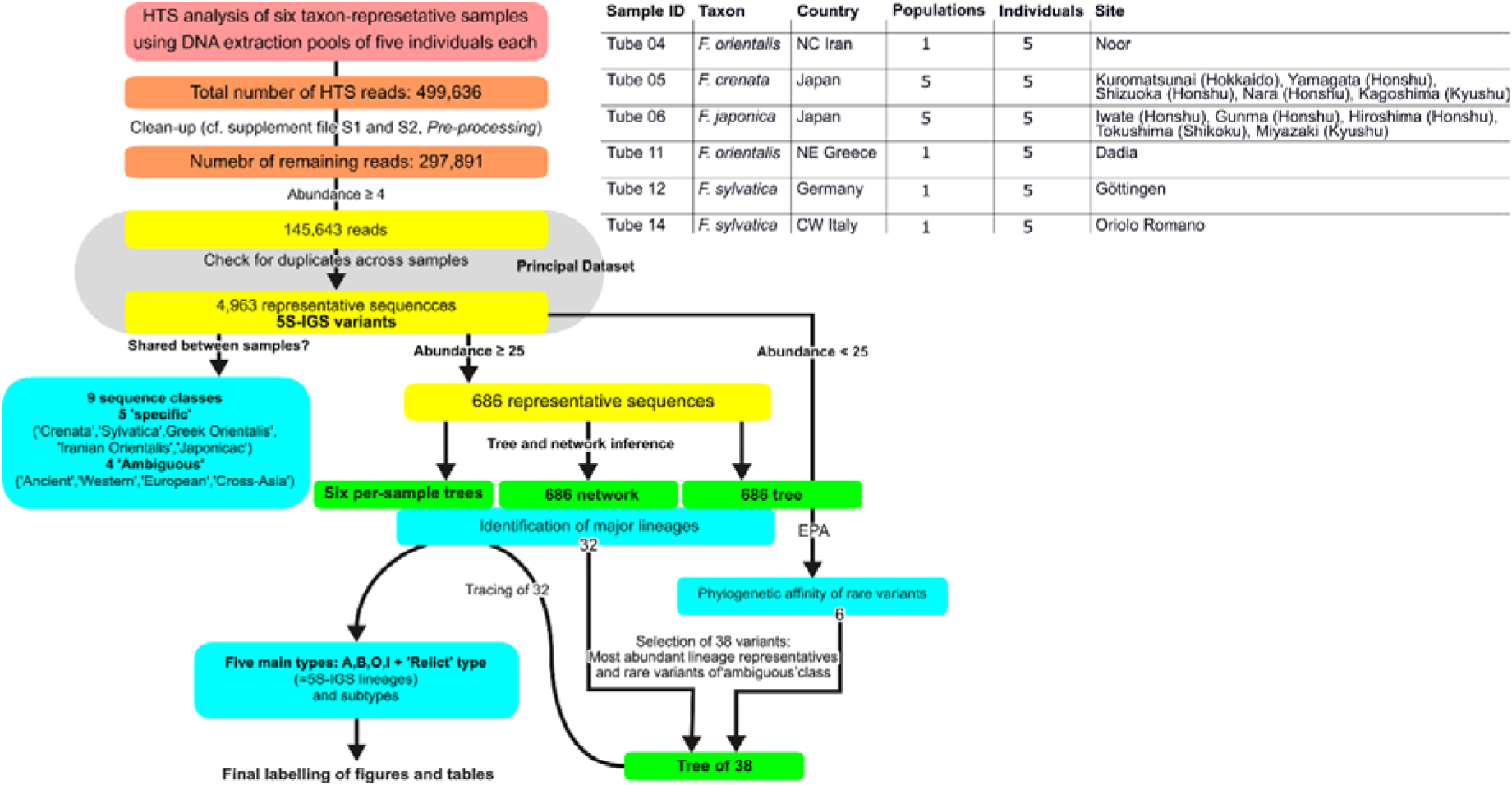
Analysis set-up and identification workflow.

Species and taxa were morphologically identified (Shen, 1992; Denk 1999a, b). DNA extractions were performed from silica-gel dried leaves with the DNeasy plant minikit (QIAGEN) and quantified with a NanoDrop spectrophotometer (TermoFisher Scientific). We prepared six artificial samples, consisting of pure species/taxa and/or populations, by pooling equal amounts of DNA of every individual up to a total of 20 ng per sample. Paired-end Illumina sequencing (2×300 bp) was performed by LGC Genomics GmbH using the 5S-IGS the plant specific primer pair CGTGTTTGGGCGAGAGTAGT (forward) and CTGCGGAGTTCTGATGG (reverse). Raw sequences were deposited in the Sequence Read Archive under BioProject PRJNA681175.

### Bioinformatics tools and workflow

Illumina paired-end reads (raw sequence data with adapters and primers clipped) were processed using mothur v. 1.33.0 (Schloss et al., 2009). Contigs between read pairs were assembled using the ΔQ parameter (Kozich et al., 2013) for solving base calls differences in the overlapping region and no ambiguous bases were allowed. Reads were dereplicated and screened for chimeras using UCHIME in *de-novo* mode (Edgar et al., 2011) within mothur and the unique (non-redundant) sequences (5S-IGS variants) with abundance < 4 were excluded from further analyses.

Given the absence of reference beech 5S rDNA sequences available in public repositories, the sequencing success of the target amplicon (5S-IGS) relied on blast searches (https://www.ncbi.nlm.nih.gov/; accessed on July 15th, 2020) within Fagales, with randomly chosen sequence reads, and reference to known literature (cf. Data S1, sections 2.2, 5).

Unique 5S-IGS sequence variants, selected based on abundance thresholds or per sample, were aligned using mafft v.7 (Katoh & Standley, 2013); MAFFT-generated multiple sequence alignments (MSAs) were manually checked and sequence ends trimmed using SeaView v.4.0 (Gouy et al., 2010). Sequence length and GC content percentage were calculated with jemboss 1.5 (Carver & Bleasby, 2003) and plotted with ggplot2 R package (Wickham, 2016). Occurrence and relative abundance of sequence reads in each sample were used for a preliminary identification of the obtained sequences (Data S1, section 2.3; Data S2). “Specific” classes included sequence reads (near-)exclusive (>99.95%) to one taxon/sample; sequences shared among taxonomically different samples were defined “ambiguous”. A graphical visualization of the links between the 5S-IGS variants in each sample and class assignation was performed by Circos plot (Krzywinski et al., 2009).

Variants with abundance ≥ 25 (686-tip set) were used in the phylogenetic analyses (trees and networks). Maximum likelihood (ML) analyses relied on RAxML v.8.2.11 (Stamatakis, 2014). Trees were inferred under a GTR + Γ substitution model using the ‘extended majority-rule consensus’ criterion as bootstopping option (with up to 1000 bootstrap [BS] pseudoreplicates; Pattengale et al., 2009). Trees’ visualization was performed in iTOL (www.itol.embl.de; Letunic & Bork, 2019) or Dendroscope 3 (Huson & Scornavacca, 2013). The Neighbour-Net (NNet) algorithm implemented in SplitsTree4 (Bryant & Moulton, 2004; Huson & Bryant, 2006) was used to generate a planar (equal angle, parameters set to default) phylogenetic network to explore incompatible phylogenetic signals and visualize general diversification patterns (cf. Hipp et al., 2020). The NNet used simple (Hamming) uncorrected pairwise (*p-*) distances computed with SplitsTree4. Intra- and interlineage diversity, between and within main 5S-IGS types (see *Results’),* was assessed with MEGA X (Kumar et al., 2018).

The phylogenetic position of variants with low abundance (<25) was evaluated with the ‘Evolutionary Placement Algorithm’ (EPA; Berger et al., 2011; Data S2, S3) implemented in RAxML. EPA was done by compiling MSAs for each sequence class and using the phylogenetic tree inferred from the 686-tip dataset; its outputs showed the multiple possible placement positions of each query sequence with different likelihood weights (probabilities) in all branches of the tree. Placement results (treejplace standard placement format) were visualized using iTOL. All phylogenetic and EPA analysis files, including used MSAs and a MSA collecting all 4693 sequences, are included in the Online Data Archive (http://dx.doi.org/10.6084/m9.figshare.16803481).

Following all phylogenetic analyses (per-sample trees, 686-tip tree and network, EPA assignation), we selected a set of 38 variants including the most abundant variants belonging to each identified lineage and the most abundant (taxon-)”specific” and “ambiguous” variants found in each sample, and additional strongly divergent and/or rare variants, for a further interpretation of the results. The autogenerated mafft MSA was checked and curated for all length-polymorphic regions using Mesquite v. 2.75 (Maddison and Maddison 2011). Phylogenetic tree inference for the selected data used RAxML included or excluded two generally length-polymorphic regions (see *Results’)* and 10,000 BS pseudo-replicates to establish support for competing splits. Data S4 documents the selection and includes a fully annotated, tabulated version of the 38-tip alignment.

## Accession numbers

All generated raw HTS sequences are deposited in the NCBI Sequence Read Archive (https://www.ncbi.nlm.nih.gov/sra) under BioProject PRJNA681175.

## Acknowledgements

EDS and GWG gratefully acknowledge the support of the German Centre for Integrative Biodiversity Research (iDiv) Halle-Jena-Leipzig funded by the German Research Foundation (FZT 118). The research was partially supported by MIUR (Italian Ministry for Education, University and Research), Law 232/2016, “Department of excellence”.

## Author contribution

GWG, ACP, JRPW, MCS conceived and designed the study

SC, RP, GWG, ACP, MCS performed research

AS, E-DS, PSS, YS, NT, JRPW contributed analytical tools and materials

SC, RP, TD, GWG, ACP, MCS analysed and interpreted data

All authors supervised and wrote the manuscript

## Data Accessibility

Used alignments and inference files are included in the Online Data Archive available at: http://dx.doi.org/10.6084/m9.figshare.16803481. Other relevant data are contained within the manuscript and its supporting materials. The original (reconstructed, phased haplotypes uploaded to gene banks) and revisited data (de-constructed individual sequences) of Jiang et al. (2021) have been included in an existing Fagaceae data collection (Grimm, 2020) and can be accessed at: https://doi.org/10.6084/m9.figshare.11603547.

## Supporting information

**Data S1** [PDF] Details about reasoning for using HTS data, sampling, 5S-IGS identification, pre-processing, amplicon length/GC content diversity within each sample/major type, and description of most prominent length-polymorphic patterns. Includes three supplement tables, 15 supplementary figures and two appendices. Appendix A is a summary of Data S2; appendix B includes violin and scatter plots sorted by sample and type.

**Data S2** [XLSX] Basic information about samples, obtained HTS reads, and completely annotated lists for the 4,693 variants with an abundance of ≥ 4, as-is and summarised as Pivot tables. Uses auto-filter, auto-generating and auto-formatting functionality; up-to-date version of Excel^®^ is recommended to properly view the file.

**Data S3** [PDF] Graphical representation of the EPA analysis of 4007 5S-IGS variants with a total abundance < 25 (full results provided in Online Data Archive, folder *EPA)*

**Data S4** [XLSX] Details about downstream in-depth analysis of 38 selected variants, including selection process and a fully annotated alignment in tabulated, graphically enhanced form.

**Data S5 [XLSX]** Tabulation and characterization of gene incongruence seen in, and phylogenetic information that can be extracted from, the 28-gene data of Jiang et al. (2021); including tree inference, bootstrapping and gene-wise model statistics.

