## Supplementary material for "5S-IGS rDNA in wind-pollinated trees (*Fagus* L.) encapsulates 55 million years of reticulate evolution and hybrid origins of modern species": Data S1

### *Data S1* – Additional Material & Methods, Results and Discussion

### 1. Rationale

#### *1.1. Using High-Throughput Sequencing (HTS) for assessing diversity in high-copy, multi-loci markers*

Assuming the occurrence of at least two functional, distinct, paralogous (or hom[o]eologous) 5S rDNA arrays in the studied beech taxa (Ribeiro et al., 2011), our data confirm the capacity of HTS to detect both of them (see **Section 4**). In addition, the HTS approach captured rare sequence types pointing to past divergences (speciation events) forming part of the 5S intergenic spacer (5S-IGS) sequence pool of the sampled populations. Although the HTS results cannot be generally considered quantitative (Lamb et al., 2019), two main sequence types were represented in all investigated taxa (**Appendix A**; cf. main-text fig. 6). Multiple loci and polyploid genomes can severely affect the generation of comprehensive phylogenetic data covering intra-genomic variation rendering traditional PCR-based direct sequencing impossible. Special attention to sequencing enough clones to capture the signal of all existing loci is therefore crucial (e.g. Volkov et al., 2017), especially when recruiting ITS data (Denk et al., 2002; Denk et al., 2005; Grimm et al., 2007a) for beech phylogenetics (four 35S rDNA loci; Ribeiro et al., 2011). For the 5S arrays, the required effort can be easily accomplished with HTS procedures, because of their optimal length for MiSeq analyses (i.e. < 400 bp; extremely conserved flanking regions). As for our work on *Quercus* (Piredda et al., 2021), the 5S-IGS has demonstrated to be a rapid and efficient tool for addressing difficult evolutionary questions and ecological applications. An obvious strength of the HTS method vs traditional cloning approaches is that even low-frequent copies are captured, including copies from potentially degrading arrays. These may indicate ancient links (e.g. Grimm & Denk, 2008) that would otherwise be overlooked because of ongoing concerted evolution and/or intragenomic silencing (cf. main-text fig. 8). The downside is clearly the amount of data that need to be analysed, which requires automated steps.

#### *1.2. Sampling & pooling*

So far, there is no 5S-IGS data for beeches. The samples used here  and data represent opportunistic research. Although HTS approaches are relatively cheap next-generation sequencing methods and do not require freshly sampled material and DNA yields as high as needed for, e.g., SNP-generating next-generation sequencing approaches, financial resources and access to relatively fresh material is the most limiting factor. At the time this research was initiated the price pre sample was ~30€ plus ~4500€ for set-up and ~500€ for sequencing of up to 192 samples. For our analysis of oaks (Piredda et al., 2021; work in progress), we were able to finance the HTS analysis of 250 samples in three sequencing batches. The number of samples was higher than originally planned,[1] hence, we could spare 10% for a pilot study into *Fagus* populations with a focus on the *F. sylvatica* s.str. – western *F. orientalis* hybrid zone in Greece.

However, to assess hybrids, it is necessary to first put up a data and phylogenetic framework for the focal species. First results demonstrated high complexity of the 5S-IGS pool in beeches (main-text fig. 2), surpassing the already complex situation in oak (Piredda et al., 2021).

Thus, for this pilot 5S-IGS study on beech, we selected four, taxonomically unambiguous, samples for sequencing:

- One sample of a *F. sylvatica* s.str. population north of the Alps, assumed to strongly have been affected by Pleistocene bottlenecks.
- One sample of *F. sylvatica* s.str. south of the Alps (C. Italy), a population growing in an area with known refuges for beech during the glacials.
- One sample each representing the westernmost (NE. Greece) and easternmost *F. orientalis* (N. Iran).[2]

For comparison, we included a sample of the supposed sister species of west-Eurasian *F. sylvatica* s.l., the Japanese *F. crenata*. *Fagus crenata* is widespread in Japan and is partly sympatric with the second Japanese species *F. japonica*, a distant relative. A sample of *F. japonica* was included here as an outgroup (cf. Denk et al., 2005 and main-text fig. 1 for inter-species phylogenetic relationships[3]). Provenance details are included in *Data S2*, sheet 'Sampling'.

As for oaks (Piredda et al., 2021), we extracted the DNA of five individuals per population and pooled the extracts into a single sample; together with the sequencing depths obtained, this ensures adequate coverage of both intra-genomic 5S-IGS diversity and inter-individual 5S-IGS

---

[1] When applying for the money, we calculated with a price of ~50€ per sample.

[2] Our sequenced beech samples unfortunately don't include any material from the Caucasus; otherwise such a sample would have been included, too.

[3] The near-resolved species tree in Jiang et al. (2021, fig. 2) is to an unknown degree an artefact. Some individuals/genes showed a high level of heterozygosity ("2ISPs"; cf. Potts et al., *Syst. Biol. 63*:1–16, 2014), hence, the authors de-phased their data to generate two (hypothetical) haplotypes per individual per gene, which were then (randomly?) concatenated and erroneously treated as independent tip taxa during all analyses. This data-naïve approach is highly problematic. As in case of much more divergent nuclear markers, the 2ISPs detected by Jiang et al. usually include the consensus nucleotide of the genus/subgeneric lineage and a species-specific fixed mutation. The concatenated sequence are thus random mixes of ancestral and species-specific artificial gene haplotypes. Inspection of Jiang et al.'s data (G.Grimm, June 2021) reveals that the reconstructed haplotypes are poorly sorted along the putative species tree and that individual genes provide little tree-like signal. The 2ISP patterns themselves exhibit a gradual shift from widely shared consensus gene sequences into species-specific intra-individual polymorphism.

divergence of the studied focal populations. The Japanese samples added as sister- and outgroups aimed at covering as much intra-species 5S-IGS diversity as possible, hence, we pooled individuals from five geographically distant populations (localities are included in **Fig. S3** in **Section 2.3**). While our data can be considered representative for each of the sampled western Eurasian populations, the Japanese samples should only be viewed being representative for their species across the covered range. Population-wise studies on Japanese beeches are likely to reveal further structuring of the 5S-IGS pools.

#### 2. Dataset Description and Classification of Sequence Reads

##### *2.1. Data curation (pre-processing steps)*

The applied HTS procedure produced 499,636 raw reads. Pre-processing (data curation) steps were performed with MOTHUR v.1.33.0 (Schloss et al., 2009). After merging the forward and reverse reads, we removed all sequences with ambiguous positions (N), those containing homopolymers longer than 30 nucleotides (nt), and any sequence longer than 400 or shorter than 200 basepairs (bp). Chimeric (artificial or naturally occurring recombinant) sequences were detected with the UCHIME algorithm (Edgar et al., 2011) in *de-novo* mode as implemented in MOTHUR. This method is optimized for detecting chimeric sequences in short, noisy sequences and produces a chimera-free database via an all-against-all pairwise sequence comparison and by exploiting the relative HTS abundance data. We then removed all sequences with abundance ≤ 3 per sample to remove possible contaminations, as in Piredda et al. (2021). The final dataset included 145,643 HTS reads. A large part of them were identical; a total of 4,693 unique 5S-IGS variants were identified (listed in *Data S2*).

In contrast to standard HTS data, 5S-IGS data may include reads with potentially artificial sequence patterns that are not filterable using standard curation procedures implemented in MOTHUR. Some of the here compiled sequences reads show imperfect ends (example provided in **Fig. S1**), which hinder the recognition and clipping of the ID-tag + primer sequence regions and may impede correct auto-alignment. Thus, for data completion, we manually checked and clipped 5’ and 3’ extensions in the 4,693-tip matrix. Folder *4693Data* in the Online Data Archive (ODA) includes both a block-aligned version of the raw data (after being processed and filtered by MOTHUR) and the cleaned alignment used for all analyses.

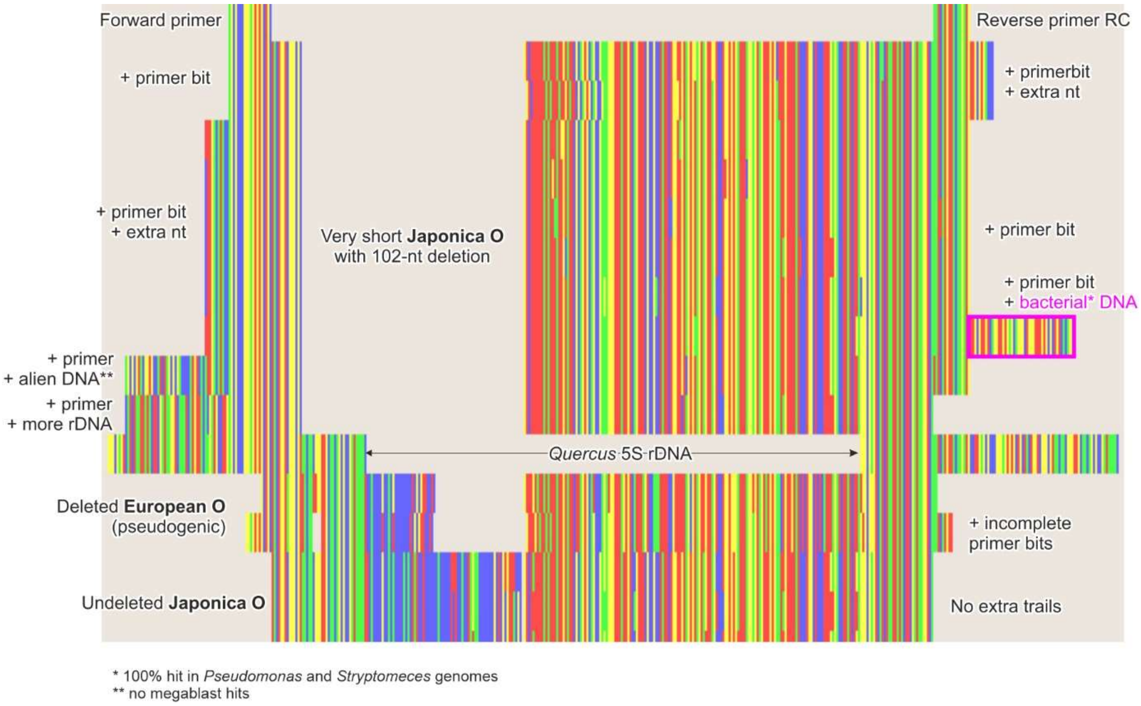

**Figure S1 | Bird's eye view of block-aligned data**; showing the imperfectly clipped ends in some sequences of short and normal-length variants of the O-Lineage (see **Section 4**). The 5S rRNA genes of *Quercus* ("Quercus 5S rDNA") and the used primer sequences (which usually are clipped automatically) are included for orientation. 'Alien' or bacterial DNA bits flanking the actual amplicons are probably ID-tags, standard pre-processing procedures failed to remove. Curious, but very rare, cases are reads where the primer bit is preceded by genuinely looking rDNA ("more DNA"), i.e. the read includes more rDNA than it should. Such patterns are occasionally encountered in cloned rDNA spacer data as well, and may point to amplification artefacts or pseudogenic, incomplete dimers.

#### 2.2. Cross-check with 5S rDNA data in gene banks

No beech 5S rDNA or intergenic spacer sequences are currently available in gene banks such as the NCBI nucleotide archive (https://www.ncbi.nlm.nih.gov/; last accessed on 25/11/2020). Randomly chosen sequence reads revealed 98–100% identity of the first 40–60 and last 30–50 sequenced bp with the 3' and 5' ends of cloned 5S-IGS sequences of various Fagales: *Quercus* spp. (oaks, Fagaceae; Simeone et al., 2018), *Alnus* spp., *Betula* spp., and *Carpinus* spp. (Betulaceae, same order; Forest & Bruneau, 2000; Forest et al., 2005), as well as with the 5S locus of an ongoing genome sequencing of *Juglans regia* (Juglandaceae, same order; BioProject: PRJNA350852). High (95–100%) identity values were also scored by these two subregions with several other eudicot nuclear-encoded 5S rRNA gene (5S rDNA) sequences, including *Morus alba*, *Punica granatum*, *Vitis vinifera*, etc. The overall mean genetic distances of the entire HTS dataset in these two sequence portions calculated as the number of base differences per site calculated by averaging over all sequence pairs with MEGA X (Kumar et al., 2018) were very low (0.01 and 0.02, respectively). We therefore concluded that the two regions belong to the 3' and the 5' portions of the highly conserved 5S rRNA gene; and confirmed this assessment visually (*Data S4*; *overview.nex* in ODA). In contrast, no high similarity BLAST scores were detected for the region comprised between these two portions, the 5S-IGS. This is not unexpected since *Fagus* is the most genetically distinct of all Fagaceae; in coding gene regions the sequence divergence between *Fagus* and the remainder of the family matches the situation found between families of core Fagales (**Fig. S2**). The ITS1 of the nuclear-encoded 35S cistron is partly unalignable (Denk et al., 2002; Denk et al., 2005).

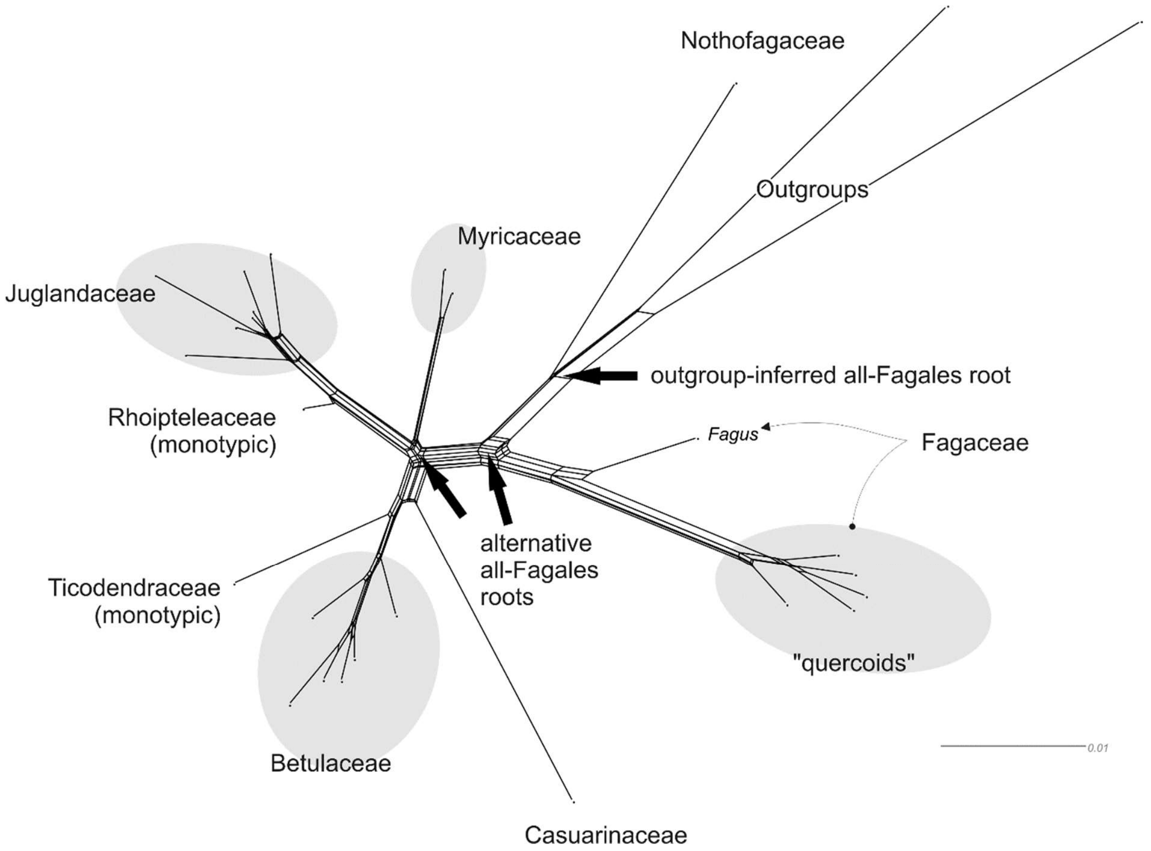

**Figure S2 | Neighbour-net for Fagales showing overall genetic divergence within the order** (after Grímsson et al., 2016, fig. 1). The graph is based on model-optimized genetic distances (scale give number of expected substitutions per site) inferred from the 6-gene matrix of (Li et al., 2004); for data see Grimm (2020). The genetic difference between *Fagus* and the remainder of the Fagaceae ("quercoids") is comparable with inter-family divergence in the rest of the crown Fagales (note: the current angiosperm classification APG IV includes the monotypic *Rhoiptelea,* Rhoipteleaceae, in the Juglandaceae).

The final, auto-generated multiple sequence alignment (MSA; included in ODA, folder *4693Data*) of the total dataset comprising 4,693 unique sequence variants was 468 bp long. A comparison with published *Quercus* 5S rDNA sequences (e.g. Tynkevich & Volkov, 2019) showed the obtained sequences comprise 44 and 34 bp of the upstream and downstream 5S coding region, respectively. Length heterogeneity was due numerous poly-nucleotide motives (mostly poly-T) and indels (insertions, deletions) of variable length occurring in the 5S intergenic spacer (5S-IGS). The gappyness of the auto-aligned MSA in the flanking upstream gene region is due to a peculiarity found in low-frequent variants involving a lack of a long portion including the end of the upstream 5S rRNA gene and the subsequent 5' part of the intergenic spacer ("short type O"; see below). A few of these 'short O' variants show a duplication (genuine or artificial) of the forward primer preceded by a non-5S rDNA four-nucleotide (nt) motif; in one variant the amplicon includes nearly two-thirds of the upstream gene while lacking the 4-nt motif (included in **Fig. S1**). The latter could be an indication that the primer duplication is not an amplification artefact but rather represents an imperfect dimer, where the IGS is eliminated as well as part of the genes.[4]

##### *2.3. Initial labelling and sample-exclusivity of sequence reads*

We categorized the obtained 5S-IGS dataset based on their sample distribution (**Fig. S3**; *Data S2*, sheet 'Representative seq_Reads'). Sequences were labelled as "specific" when exclusively found in the one or two of the samples representing the same taxon: "Japonica", "Crenata", "Iranian orientalis", "Greek orientalis", and "Sylvatica". Four 'Sylvatica' variants, including the overall most abundant one, can be found in Greek *F. orientalis* but with abundances ≤ 2 (<0.001%). Conversely, one 'Greek orientalis' variant was found in *F. sylvatica* from Germany. We identified four "ambiguous" classes, i.e. sequences shared among different species or taxa (with an occurrence of 1.3 to 42.7% per sample): 175 variants, classified as "European" and corresponding to 7,271 sequences are exclusively shared between *F. sylvatica* s.str. and Greek *F. orientalis*, while substantially rarer variants (three variants representing 21 sequences) shared across all western Eurasian samples were labelled as "Western". The last two shared sequence classes connect disjunct beech populations: "Ancient" is exclusively shared by *F. sylvatica* s.str. and *F. orientalis* from Iran (three variants representing 45 sequences); "Cross-Asia" is a variant corresponding to 73 *F. crenata* HTS reads and a single sequence of *F. orientalis* from Iran.

[4] Complete (perfect) dimers would have been filtered automatically during the pre-processing being > 400 bp.

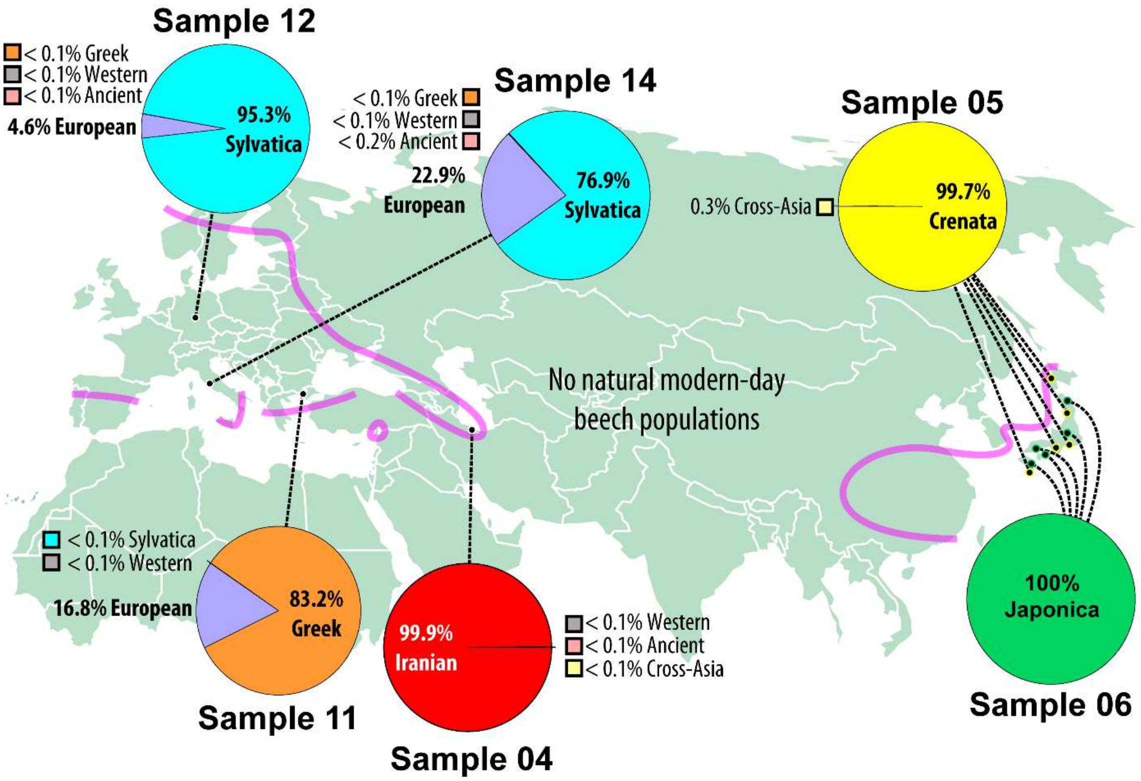

**Figure S3 | Per-sample distribution of "specific" (coloured) and "ambiguous" (grey) 5S-IGS sequence classes.** Based on the investigated dataset of 4,693 representative sequences, i.e. non-identical 5S-IGS variants with an abundance of ≥ 4. Purple lines give approximate natural boundaries of beech in western Eurasia and East Asia (after Denk & Grimm, 2009, fig. 3); see also https://www.gbif.org/species/2874875.

The *F. japonica* pooled sample included only exclusive sequences (100% 'Japonica'; **Fig. S3**); likewise, *Fagus crenata* and Iranian *F. orientalis* are near-exclusive (>99.5% "specific" sequences). In contrast, *F. orientalis* from Greece has only 83% "specific" sequences, whereas Italian *F. sylvatica* showed the highest percentage of shared ("ambiguous") variants (23%). The highest number of sequences were shared among *F. sylvatica* s.str. (both provenances) and Greek *F. orientalis* (class 'European'); only a few (0.003 – 0.01%) were shared between Iranian *F. orientalis* and *F. sylvatica* s.str., all western Eurasian beeches, or *F. crenata*. The amount of shared and exclusive ("specific") 5S-IGS variants clearly distinguish the geographically isolated Iranian populations of *F. orientalis* from the Greek one, the latter being much more interconnected with nearby *F. sylvatica* s.str.

#### 3. General Features of *Fagus* 5S-IGS

The obtained 5S-IGS sequences were highly variable in structure and length, both at the intra- and interspecific level (**Figs S4, S5**; **Appendices A, B**; *Data S2,* sheet 'GC content & length'), with an overall mean genetic distance of 0.1. The GC content of the amplicons ranged from

33.2 to 45.1% (23.9–44.8% for the spacer), and the length range was 166–307 bp (spacer: 88–229 bp). Such a structure and length variability can be found in different plant groups (e.g. Fulnecěk et al., 2002; Negi et al., 2002; Forest et al., 2005; Denk & Grimm, 2010; Grimm & Denk, 2010; Garcia & Kovařik, 2013; Mlinarec et al., 2016; Garcia et al., 2020).

The average GC content is lower than that found in these studies and in the only other genus of Fagaceae studied to date (typical GC content ~50%; Denk & Grimm, 2010; Simeone et al., 2018; Tynkevich & Volkov, 2019; Piredda et al., 2021). Interestingly, ongoing full genome sequencing projects of *F. sylvatica* s.str. and *F. crenata* (GenBank acc. no. QCXR00000000.1 and BKZX00000000.1; not yet annotated, accessed on 25/11/2020) reported very low preliminary GC content (34.9% and 32.9%, respectively). *Fagus* further differs from other Fagaceae in general, particularly oaks, by substantially longer ITS regions including more prominent AT-dominated sequence portions and, consequently. lower GC percentages, especially in the ITS1 (Denk et al., 2002; Denk et al., 2005; Denk & Grimm, 2010).

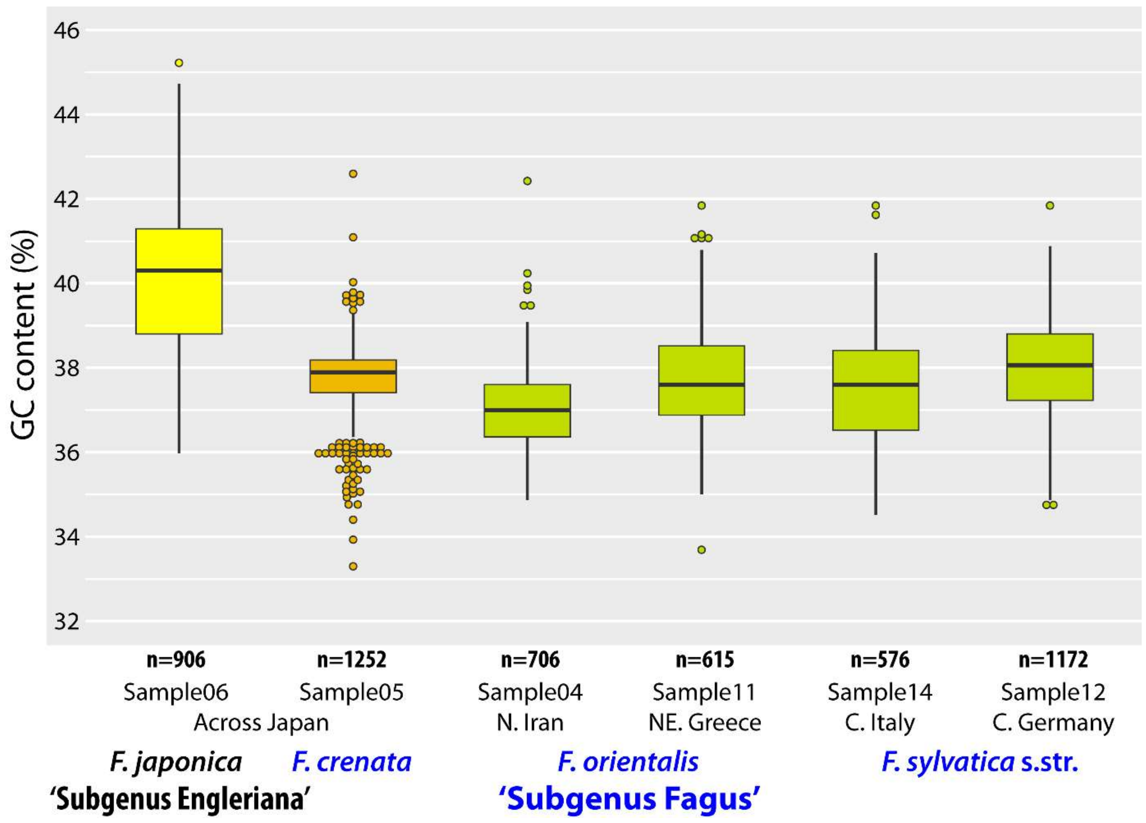

**Figure S4 | Boxplots of amplicon GC contents per sample.** Boxes give the median value and the 25th to 75th percentiles, whiskers refer to the entire range without outliers (dots) as defined by Tukey's formula (Tukey, 1949).

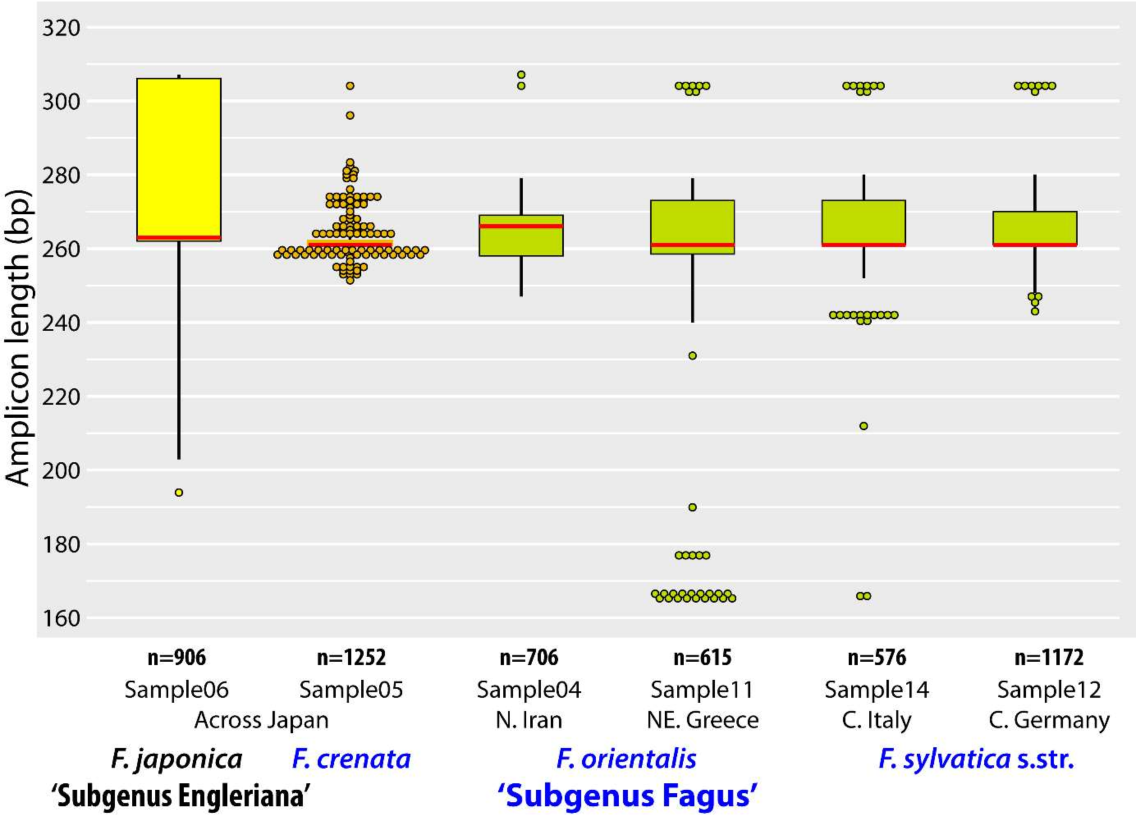

**Figure S5 | Boxplots of amplicon lengths per sample.** Boxes give the median value (in red for contrast) and the 25th to 75th percentiles, whiskers refer to the entire range without outliers (dots) as defined by Tukey's formula (cf. **Fig. S4**).

The GC content (**Fig. S4**) and length variation compiled per sample (**Fig. S5**) pointed to a structural difference between 5S-IGS of *F. japonica* (Subgenus Engleriana) and the other taxa (crown group of Subgenus Fagus), with numerous sequences showing over-average GC content and length. All other samples were, as a trend, more homogeneous in sequence structure. The GC range for the two combined *F. sylvatica* s.str. samples was 34.4–41.7% (median = 37.9%). The Iranian population of *F. orientalis* showed the lowest GC contents (median = 36.9%) but highest median length (266 bp).

##### *3.1. Relationship between GC content/amplicon length and abundance*

The GC content vs abundance scatterplot of the full dataset (**Fig. S6**) displays a bimodal distribution, with two peaks at 43.6% (*F. japonica*) and 37.9% (*F. crenata*) and abundances of 772 and 7730, respectively. The largest part of the 5S-IGS sequences (98.9% = 144,094 reads; representing 4,616 out of 4,693 non-identical variants) fell within a (>)34–(<)43% range, whereas a second group with higher GC content (>43%) included 74 variants with a total abundance of  1516 reads found exclusively in *F. japonica* (one frequent variant; remaining individual abundances ranging between 4–60). Three rare variants showed GC content <34% (**Fig. S6**). The length/abundance scatterplot (**Fig. S7**) shows four sequence clusters: (*i*) a relatively low-frequent group of very short sequences (<180 bp); (*ii*) a length-homogenous group (203–204 bp) with low to high abundances; (*iii*) a large cluster of sequences with an abundance maximum around ~260 bp; and (*iv*) a low- to high abundance, length-homogenous group centred around ~305 bp (303–307 bp).

In *F. japonica*, we found all four main length clusters, and a single sequence of 222 bp (with abundance of 16) missing 30 bp of the central IGS. The very short (≤ 170) and 204-bp cluster (102 5S-IGS variants represented by a total of 1865 reads; 5.8% of this species'/sample's total) share a 102 nt-long deletion involving 31 bp of the 3' end of the 5S rRNA gene (starting at invariable alignment pos. 14 and represented by "Ja_short" in *Data S4*, sheet 'Selected variants'); all but one belong to the group showing a GC content ≥ 44%.[5] The most length-homogeneous sample was *F. crenata* (**Fig. S5; Table S1**).

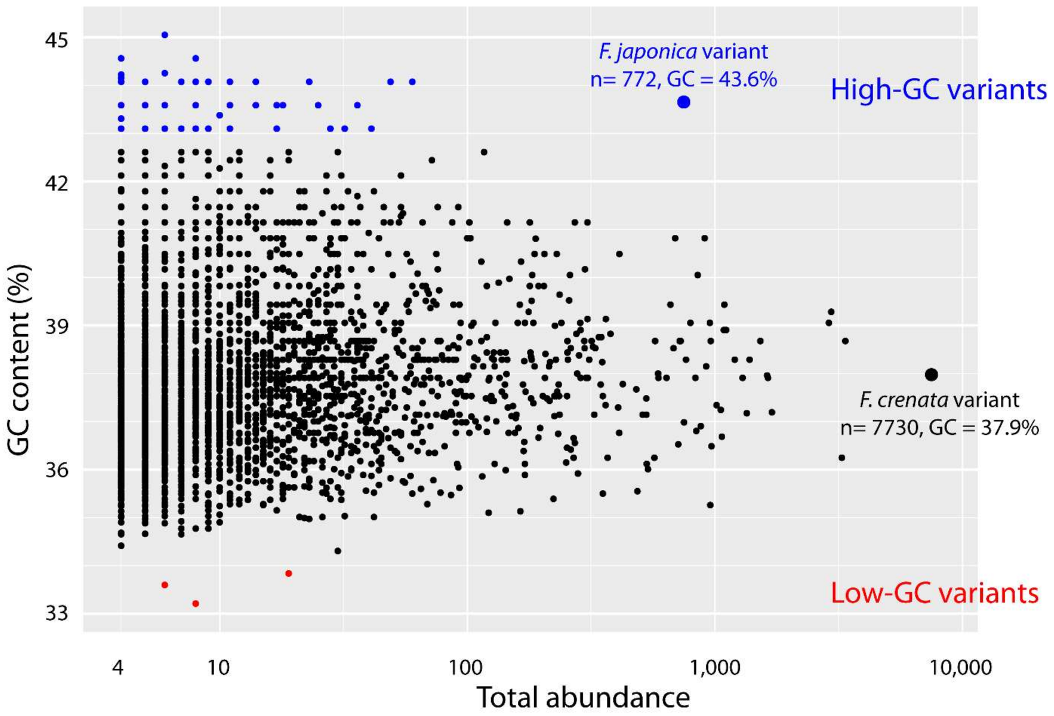

**Figure S6 | x-y plot of the GC content and abundance of the 4,693 5S-IGS sequences**. Shown is total abundance, i.e. the sum across all samples (cf. *Data S2,* sheet 'GC content & length').

[5] The sequence with lower GC content belongs to a different main 5S-IGS lineage ('Japonica I'; see **Section 4.3**)

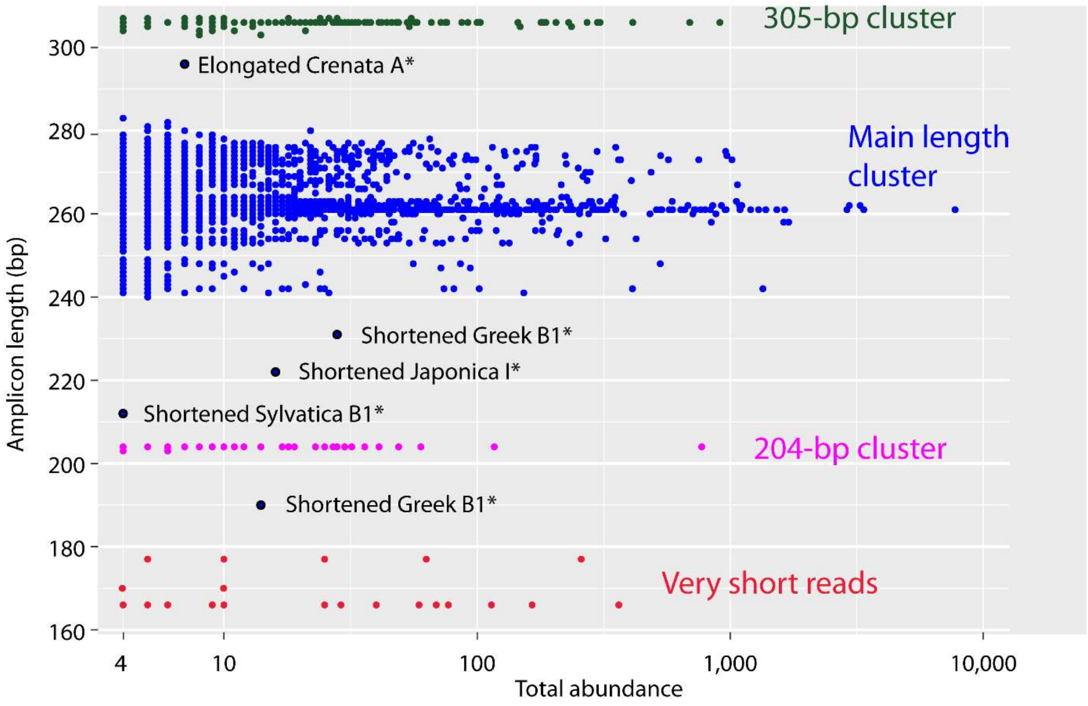

**Figure S7 | x-y plot of the length and total abundance.** Downstream phylogenetic analysis (cf. main-text figs 3–5; *Data S2*) shows that sequences with a length of ~280–300 bp either represent shortened versions of the 305-bp cluster or elongated versions of the main length cluster. * For labels see **Section 4** below, and main text.

The Iranian *F. orientalis* population showed a high diversity in sequence length and GC contents (**Figs S4, S5**). Most sequences of *F. sylvatica* s.str. from Italy (99.6%; 565 variants) and *F. orientalis* from Greece (92.4%; 583 variants; **Table S1**), the samples with the most pronounced length polymorphism (**Fig. S5**), fell within the main length cluster. Most of the observed length variation within the main length cluster relates to the co-occurrence of two main 5S-IGS lineages in samples representing *F. sylvatica* s.l. (cf. main-text fig. 7) of ~240–270 (B-Lineage) and ~260–280 bp (A-Lineage). Very few sequences of *F. sylvatica* s.l. (in total 10 variants with a sum-abundance of 100, i.e. ~1 ‰ of all reads) have a length ≥ 303 bp. The group of very short variants (23 “Greek-orientalis specific”, one “Sylvatica-specific”, one “ambiguous” variant, 166 or 177 bp long with abundances of 4–363) showed an identical ~95 nt-long deletion (represented by “OrG_short” in *Data S4*, sheet ‘Selected variants’). One variant belonging to the Greek sample of *F. orientalis* (length = 231 bp; GC content = 36.4%; abundance = 28) had a 30 nt-long deletion involving 29 bp of the 3’ end of the (upstream) 5S rRNA gene. As in the case of the short *F. japonica* variants (204-bp length cluster), the very short *crenata-sylvatica* lineage variants and the variant missing the end of the upstream 5S rRNA gene show no further evidence of sequence degradation. The downstream 5S rDNA bits are inconspicuous in showing the genus’/family’s consensus sequence.

**Table S1.** Number of non-identical sequence variants (NIV) and total abundance (TA) of length clusters (as defined in **Fig. S7**).

| | *F. japonica* | | *F. crenata* | | Iranian *F. orientalis* | | Greek *F. orientalis* | | *F. sylvatica* s.str. | |
|---|---|---|---|---|---|---|---|---|---|---|
| Length cluster | NIV | TA | NIV | TA | NIV | TA | NIV | TA | NIV | TA |
| Very short | 3 | 18 | – | | – | | 23 | 1345 | 2 | 12 |
| 203–204 bp | 99 | 1847 | – | | – | | – | | | |
| Main length cluster | 410 | 20785 | 1250 | 27364 | 704 | 20213 | 583 | 17100 | 1227 | 18303 |
| …range | 258–276 bp | | 252–283 bp | | 247–279 bp | | 240–279 bp | | 241–280 bp | |
| ~305 bp | 393 | 9559 | 1 | 4 | 2 | 14 | 7 | 20 | 9 | 66 |
| Total | 906 | 32225 | 1252 | 27375 | 706 | 20227 | 615 | 18507 | 1239 | 18385 |

#### 4. Phylogenetic Sorting

The subsequent sorting of the obtained dataset into five main phylogenetic lineages (O-, I-, and X-Lineage types in *F. japonica*; A- and B-Lineage in *F. crenata – F. sylvatica* s.l.; cf. main-text figs 3–5, *Data S2–S4*) revealed a structural homogeneity of the sequences included in each major lineage (**Fig. S8**; **Appendix A**; see also main-text fig. 7). The *crenata-sylvatica* A-Lineage variants have generally lower GC content and are shorter than those of the B-Lineage in each species/sample; the I-and X-Lineage variants (*F. japonica*) correspond to the B-Lineage of the other taxa regarding GC content and length ranges. O-Lineage variants (*F. japonica*) include longest 5S-IGS variants (typically ≥ 300 bp; 305 bp-class) and highest GC contents.

**(Following page) Figure S8 | x-y plots of GC content vs amplicon length**; sorted for the five main 5S-IGS lineages (A-, B-, I-, O-, X-Lineage) and include 5S-IGS variants forming the 'Relict Lineage'. Dots may represent more than one variant, since variants may have (near-) identical GC contents and lengths. The aberrant I-type ("SyG3" in *Data S4*) is a sequentially unique, weakly degraded variant obtained from the German *F. sylvatica* sample without affinity to any major sequence type and placed within the I-Lineage subtree by EPA (Evolutionary Placement Algorithm). * Low-frequent (total abundances ≤ 36) O-Lineage variants ('European O'; see **Section 4.3**) can be found in all samples of the *crenata-sylvatica* lineage.

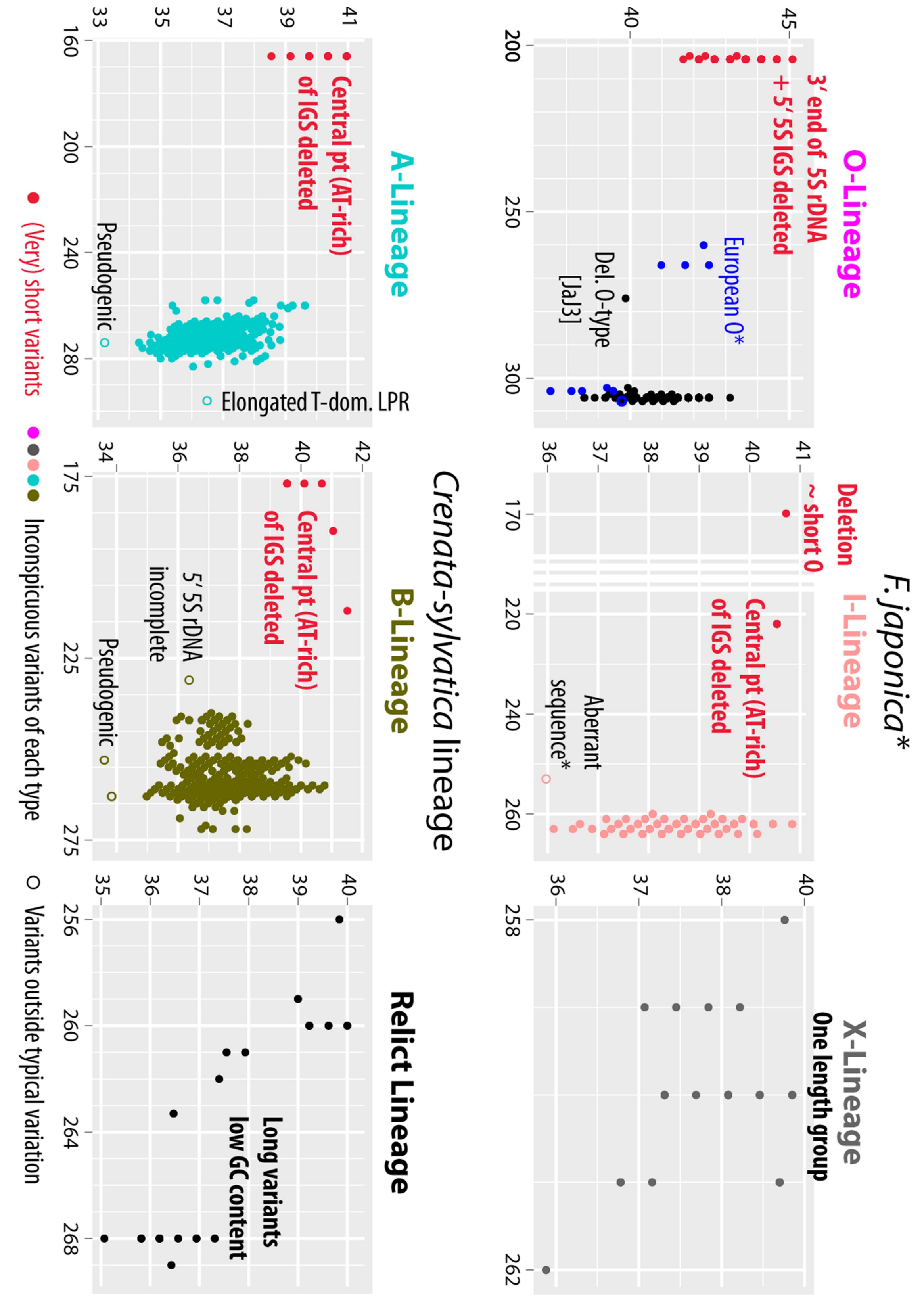
O-Lineage
3′ end of 5S rDNA + 5′ 5S IGS deleted
European O*
Del. O-type [JaJ3]
*F. japonica**
I-Lineage
Deletion ~ short O
Central pt (AT-rich) of IGS deleted
Aberrant sequence*
X-Lineage
One length group
A-Lineage
Central pt (AT-rich) of IGS deleted
Pseudogenic
Elongated T-dom. LPR
*Crenata-sylvatica* lineage
B-Lineage
Central pt (AT-rich) of IGS deleted
5′ 5S rDNA incomplete
Pseudogenic
Relict Lineage
Long variants low GC content
(Very) short variants
Inconspicuous variants of each type
Variants outside typical variation

##### 4.1. Sequence features of I-, O- and X-Lineage types of F. japonica

In *F. japonica*, the three distinct and conserved length classes (**Table S1**) sort phylogenetically. I- and X-Lineage variants have similar length (~260 bp; main length cluster) and GC contents; O-Lineage variants have higher GC contents and form the 204- and 305-bp length clusters (**Fig. S9**). The short (203–204 bp) sequences represent truncated O-Lineage variants ("Short O" in *Data S4*), missing 30 bp of the upstream 5S rRNA gene and the 5' end of the 5S-IGS up till and including all but one nucleotide of the T-dominated 5' length-polymorphic region (LPR; *Data S4,* sheet 'Motives'). While such a deletion is characteristic for pseudogenic, dysfunctional rRNA arrays/copies, the remainder of the sequences including the downstream 5S rDNA are inconspicuous, hence, the very short O-type sequences show the highest observed GC contents (**Fig. S8**).

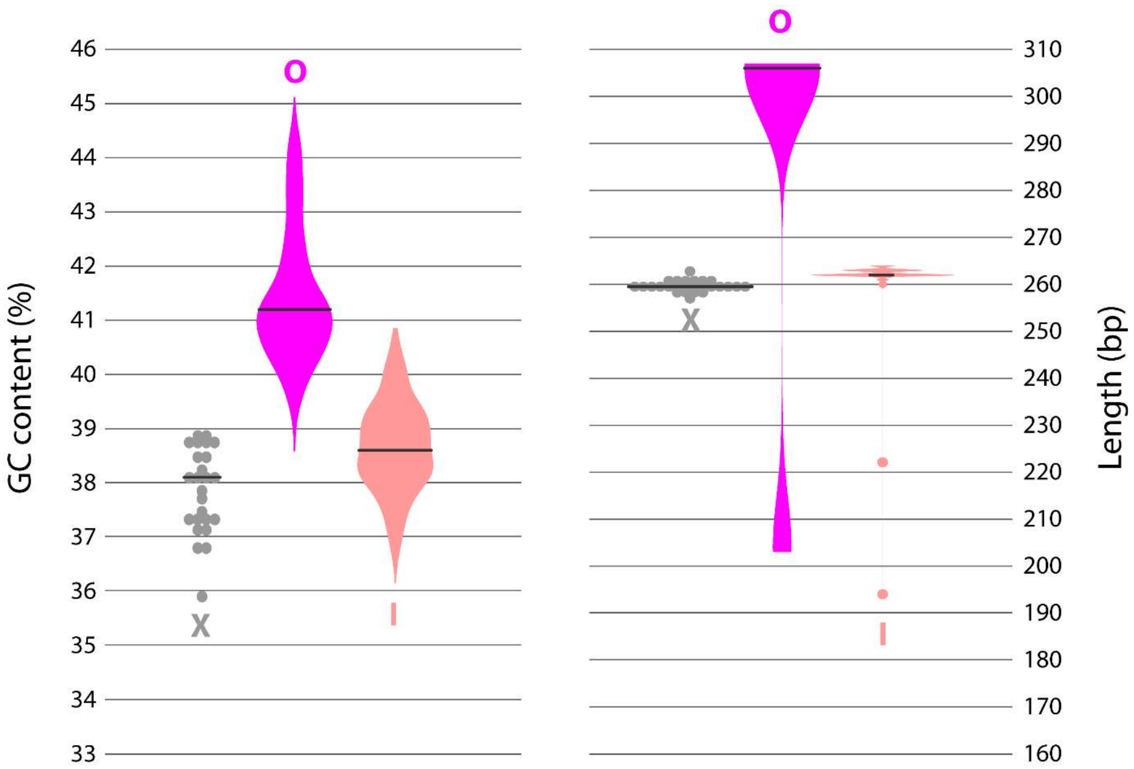

**Figure S9 | Violin and scatter plots of GC content and sequence length for three 5S-IGS lineages observed in *F. japonica*.** O-Lineage variants represent an outgroup 5S-IGS sequence type (see main-text), X-Lineage variants a indistinct ingroup type, and I-Lineage variants the sister lineage of *crenata-sylvatica* B-Lineage.

The most apparent diagnostic sequence feature of the O-Lineage is a much-elongated central length-polymorphic region, the “semi-homologous region”[6] defined in *Data S4*, and a generally high GC content in the length-homogenous portions of the spacer. The sequential difference to the ‘ingroup’ A-, B- and I-Lineage variants is striking, allowing to quickly identify O-type sequences when viewing (auto-generated) alignments in bird’s view[7]. The 3’ variable region following downstream of the semi-homologous region shows an even increased amount of point mutations, mostly transitions leading to a higher GC content than in A-, B-, I- and X-Lineage variants.

The I-Lineage variants, the most dominant of the *F. japonica* types (**Appendix A**), match in GC content and length diversity the *crenata-sylvatica* B-Lineage (**Appendix B**). Sequentially, it represents the *F. japonica* counterpart to the *crenata-sylvatica* B-Lineage as well, sharing several highly diagnostic sequence features differentiating between A-Lineage and B-Lineage (*Data S4*). For instance, its T-dominated 5’ LPR is sequentially most similar to that of putatively derived B-Lineage types such as ‘European B’ and ‘Iranian B1’ (**Fig. S10**). It differs from both types by several conserved point mutations, mostly transversions, in the lineage-discriminating central part of the 5S-IGS, in-between the T-dominated 5’ LPR and the semi-homologous region.

In contrast to the variants forming the O- and I-Lineage, the rare X-Lineage variants seem to lack discriminative sequence patterns obscuring its relationship with respect to the A-Lineage and B+I sister lineages (exemplified in **Fig. S11** for the 5’ T-dominated LPR). The 686-tip tree placed the X-Lineage as first diverging lineage within the ‘ingroup’ clade; its ambiguous, non-discriminate signal caused the collapse of branch support for deep (inter-lineage) relationships (main-text fig. 3). Sequentially, I-type sequences clearly represent an ‘ingroup’ type, which however lacks any clear diagnostic or phylogenetically informative feature in contrast to A- and B-/I-Lineage sequences. When deviating from the ingroup consensus, i.e. the modal consensus of A-, B-, I- and X-Lineage variants, it either shows sequence patterns diagnostic for the A- Lineage, the B+I clade or found as intra-lineage variation in either of them (**Table S2**).

---

[6] While position homology is straightforward to establish within each main-lineage, alternative alignments are possible between main lineages within this generally length-polymorphic and sequentially high-divergent part of the 5S-IGS. While the semi-homologous region (SHR) as a whole is homologous between “outgroup” type O and “ingroup” types, it is unclear which bit of the much-elongated SHR motif of type O corresponds to bits seen in “ingroup” types.

[7] The ODA includes a NEXUS version of the auto-aligned total 5S-IGS data, annotated with and optimised for viewing with MESQUITE, which provides a bird’s eye view option [Maddison, W. P., & Maddison, D. R. (2011). *Mesquite: a modular system for evolutionary analysis. Version 2.75*. In [new] https://mesquiteproject.wikispaces.com/ ]

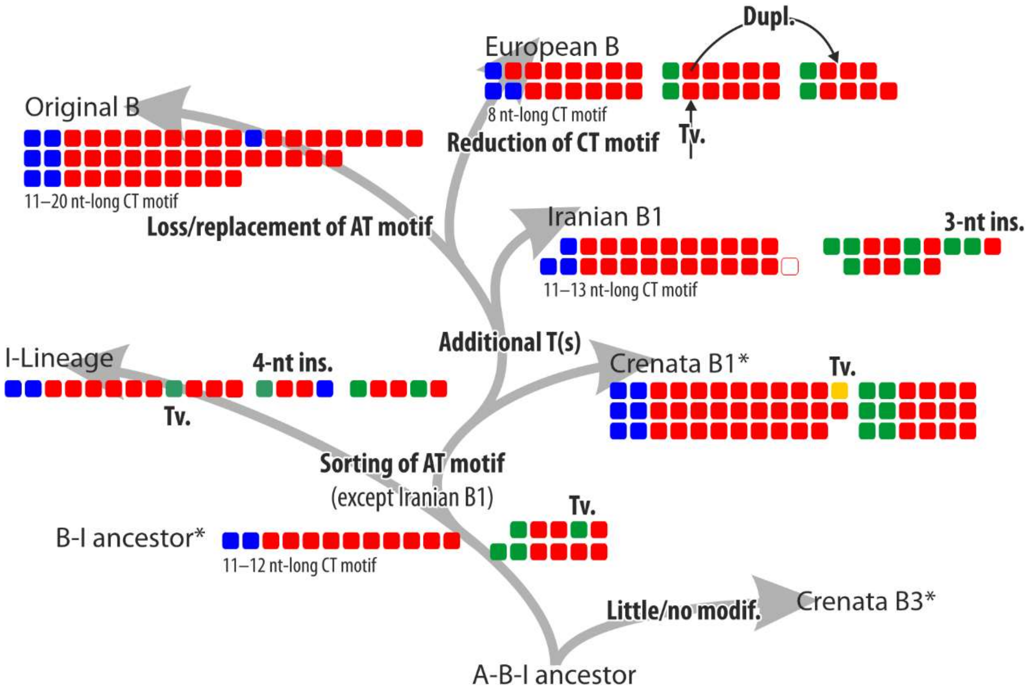

**Figure S10 | Differentiation of the 5′ T-dominated length-polymorphic region (5′ LPR) within the I-B lineage.** Coloured squares represent nucleotides: green = adenine (A), blue = cytosine (C), dark yellow = guanine (G), red = thymine (T); mixed colours uncertainties (hypothetical ancestral sequence) or polymorphisms. Empty squares reflect (here: 1-nt) length polymorphism.***** Crenata B1 variants, accounting for > 80% of *F. crenata* 5S-IGS reads, may show a potentially ancestral 5′ LPR oligonucleotide motif, while the 5′ LPR of very rare Crenata B3 variants lacks the otherwise shared, B-I-lineage-diagnostic features (cf. **Fig. S11**). Abbrev.: Dupl. = duplication; Ins. = insert; Ts. = transition; Tv. = transversion.

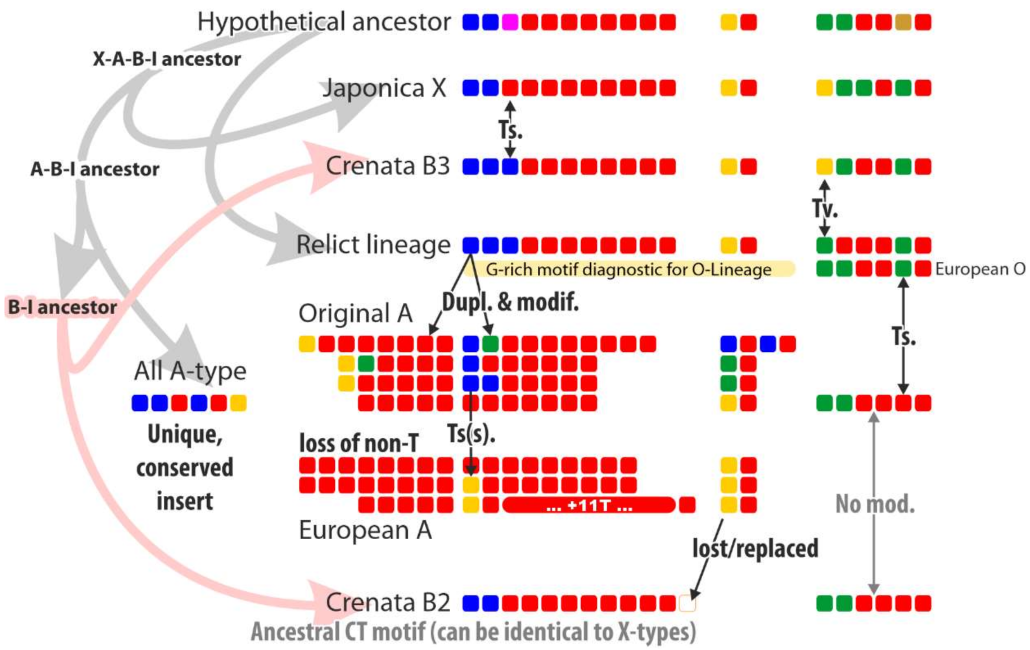

**Figure S11 | Hypothetical (early) evolution of the 5′ T-dominated length-polymorphic region in the 5S rDNA intergenic spacer of beeches**, leading to low-modified B-subtypes and diagnostic A-Lineage elongation (see also *Data S4,* sheet 'Motives'). The phylogenetic relationships between the shown types follow the result of the 686-tip (main-text figs 3, 4) and 38-tip dataset analyses (main-text fig. 5). Same colour coding and abbreviations as in **Fig. S10.**

**Table S2 | Phylogenetic ambiguity of X-Lineage variants**. Matches are highlighted by bold font and colour. IUPAC polymorphism codes used when there are co-dominant motives within each lineage (i.e., reflect plurality consensi; for complete documentation see *Data S4*)

| Lineage | Pos. 81ff | T-dom. LPR (96–218) | 135ff | 147f | 157 | SHR[a] (174–241) | 275 |
|---|---|---|---|---|---|---|---|
| Affinity of X | → B | → B | Outgroup | None | Ingroup[b] | Ingroup | Ambiguous[c] |
| **X** | TT**T**-TATA | **"Ancient B"** | **CAA**[d] | TT | **C** | **"Ancestral"** | **T** |
| **A** | TTG-TATA | "Long" | CTT | RA | **C** | **"Ancestral"** | A |
| | | | | | | "Asian types" | |
| **I** | TTG-TATA | "Ja intype" | CGA | GA | T | **"Ancestral"** | **T** |
| **B** | TT**K**-TATA | **"Ancient B"** | CGA | RA | **Y** | **"Ancestral"** | **T** |
| | | "Origianl B" | YTA | | | "Asian types" | |
| | | "European B" | TCG | | | Various derived types | |

[a] Semi-homologous region

[b] Usually candidate for an ancestral sequence feature within the ingroup; subsequently lost or modified in the ingroup sibling lineages.

[c] Highly conserved SNP: always T in 'Japonica O', B- and I-Lineage variants; consistently A in A- and Relict Lineage variants.

[c] Similar to O-Lineage types: AAA, **CAA** in 'European O'

The lack of decisive phylogenetic sequence patterns (**Figs S10, S11; Table S2**) is the main reason for the low support in the 486-tip tree (main-text fig. 3); thus, we did not include X-Lineage representatives in the select 36-sequence matrix (main-text fig. 5).

Without more data on other East Asian beech species, the (sequence-wise) rather primitive, underived *F. japonica* X-Lineage must remain an enigma. Possible evolutionary scenarios include:

- The X-Lineage comprises ancient 5S-IGS variants, originally shared by all lesser-evolved East Asian species (cf. *Data S5*[8]), i.e. left-over of incomplete lineage sorting.
- A common sequence type found in not sampled Chinese beech species, i.e. evidence for (past) reticulation.
- A rare relict variant exclusive of Subgenus Engleriana (possibly shared with the North American *F. grandifolia*[9]), largely replaced by I-Lineage variants in *F. japonica*.

##### *4.2. Sequence features of A- and B-Lineage types* of the F. crenata – F. sylvatica *s.l. lineage*

Downstream analysis of 38 select variants showed that much of the observed length variation relates to the co-occurrence of two main 5S-IGS lineages, A-Lineage (~265–275 bp) and B-Lineage (~240–260 bp; **Fig. S12**). The split between the two lineages is exemplarily reflected in the motives of the T-dominated LPR at the 5' end of the 5S-IGS (**Fig. S10)**. Conserved point mutations characteristic for the A-Lineage are concentrated directly downstream of the T-dominated 5' LPR. One 'Crenata A' sequence falls out of the usual length range (marked in **Fig. S8**) because of a much-elongated T-dominated LPR showing multiple repetitions of upstream $T_x$-C and downstream A-$T_y$ oligonucleotides ($x = 2–3$; $y = 4–7$) connected by five T. Sequences with ~300 bp belong to the same lineage as the *F. japonica* O-type variants ('European O'; see **Section 4.3**).

In addition, sample #11 (Greek *F. orientalis*) includes 18 variants ("OrG_short" in *Data S4*) with a similar-located 107-nt (A)/ 86-nt long (B) deletion in both A- and B-Lineage starting

[8] The East Asian clade in Jiang et al. (2021, fig. 2) is probably paraphyletic having near-zero character support (in total three near-conserved point mutations, scattered across three genes: P12, P21, and P54). It represents a data-/branching-artefact triggered by the use of reconstructed (including chimeric) haplotypes in a data situation where at least one member of the East Asian clade includes sequences that are ancestral to all other reconstructed haplotypes found in Subgenus Fagus.

[9] In several of the genes collected by Jiang et al. (2021), *F. grandifolia*, while being generally distinct from all other species, is genetically closer to the species of Subgenus Engleriana than the Eurasian members of Subgenus Fagus. See also Denk & Grimm (2009) for fossil links.

with the T-dominated 5' LPR and stopping before (B-Lineage types) or within (A-Lineage types) the semi-homologous region. As in the case of the short O-Lineage sequences, pseudogenic mutations are extremely rare in the flanking rDNAs. Since these short variants ('very short' length cluster; **Section 3.1, Fig. S7**) deleted most of the lineage-discriminating central part of the 5S-IGS, they are ± equally distant to both A- and B-Lineage variants, hence, collected in the central part of the neighbour-net splits graph (main-text fig. 4). Their assignation to A- and B-Lineage using EPA or detailed sequence-inspection is straightforward because of the lack (A-types) or occurrence (B-types) of lineage-specific, subtree-sorted point mutations in the non-deleted, length-homogenous but lineage-divergent 3' variable region and the undeleted part of the semi-homologous region (*Data S4*, sheet 'Motives').

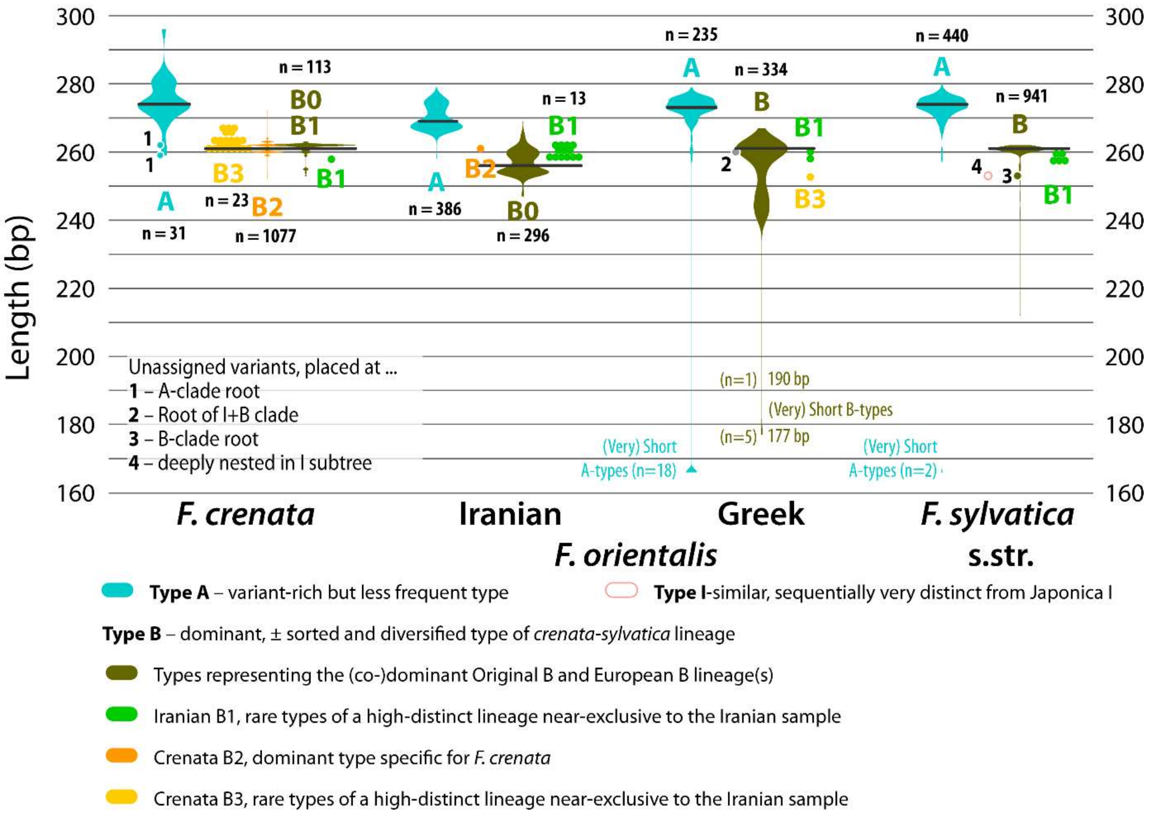

**Figure S12 | Violin and scatter plots of amplicon lengths;** shown are 5S-IGS sequence types of the A- (Type A) and B-Lineage (Type B) characteristic for the *crenata-sylvatica* lineage (**Section 4.2**).

B-Lineage variants show a higher GC content than A-Lineage variants (**Fig. S13**) and a much higher oligonucleotide motif diversity in the two generally length-polymorphic regions, the T-dominated 5' LPR (**Figs S10, S11**) as well as the semi-homologous region. Both length-polymorphic regions can be highly diagnostic for dominant subtypes such as 'European B' and 'Crenata B2'. In-depth analysis of the select set as well as visual inspection of the 4,693-sequence alignment (NEXUS-file included in the ODA, folder *4963Data*) indicates that A-Lineage variants include not only longer (always AT-rich) length polymorphic regions but also more variants with an increased proportion of point mutations that could be related to (beginning) pseudogeny, hence, the trend towards lower GC contents. This agrees with the overall abundance pattern of a very few, typically low-GC A-types in *F. crenata* and increasing dominance of B-type sequences, lowest in the easternmost Iranian and highest in the north-westernmost German population. Overall, the pattern points to an ongoing elimination (silencing) and replacement of (more primitive) A-Lineage by (more derived) B-Lineage arrays in the genomes of the *crenata-sylvatica* lineage (discussed in main text).

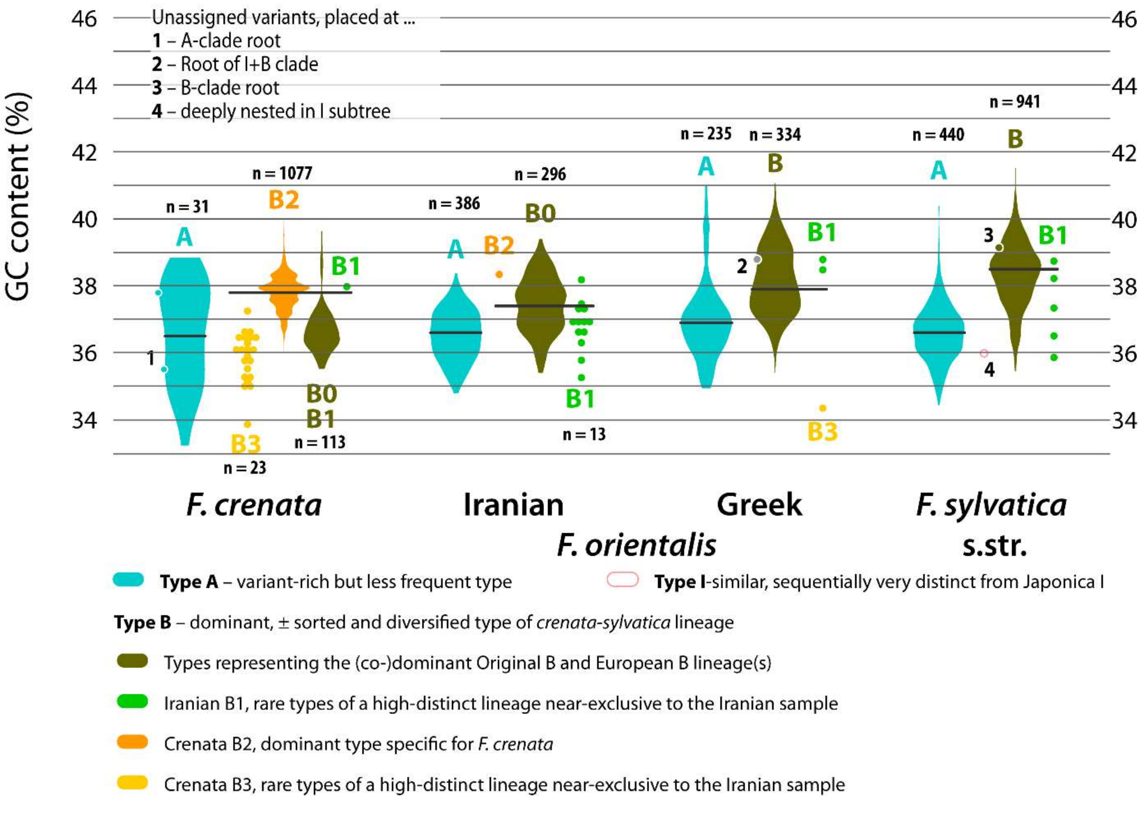

**Figure S13 | Violin and scatter plots of GC contents;** shown are 5S-IGS sequence types of the A- (Type A) and B-Lineage (Type B) characteristic of the *crenata-sylvatica* lineage.

##### 4.3. Relict sequence types and taxonomic mismatches

In addition to the main (co-dominant), phylogenetically and taxonomically ± sorted 5S-IGS types, rare variants can be found in all members of the *crenata-sylvatica* lineage that show a stronger affinity to the variants of the outgroup sample (*F. japonica*) and corresponding amplicon lengths and GC contents (**Fig. S14**).

***European O lineage*—**Variants labelled as "European O" show all distinctive characteristics of the *F. japonica* O-Lineage ('Japonica O') such as the much-elongated, GC-rich semi-homologous region (SHR). The SHR of 'Japonica O' and 'European O' variants is sequentially not identical but structurally corresponding (*Data S4*); highest number of distinguishing point mutations are concentrated in a 24 nt-long stretch at the 3' end of the SHR. All 'European O' variants show an increased amount of putatively pseudogenous transitions from G→A and C→T, in both the spacer and the flanking gene regions. Only a single 'European O' variant is relatively frequent (total abundance of 36; shared by both *F. sylvatica* s.str. samples; "D_ASOG" in the 38-tip dataset), hence, included in the 686-tip dataset, and resolved as sister to 'Japonica O' clade (cf. main-text figs 3, 5; assuming a midpoint root). 'European O' variants show a minor (~40 nt-long) length polymorphism: in the shortened variants (such as D_ASOG) the part directly upstream of the (shortened) T-dominated 5' LPR is deleted. 'European O' variants are more frequent and diverse in the European samples of the *crenata-sylvatica* lineage (**Fig. S14**), with only three variants found in Iranian *F. orientalis* (total abundance, TA, of 24; 0.12% of all post-processing reads; two of class "specific", one "ambiguous" shared with European samples) and *F. crenata* (TA = 30; 0.11%; all "specific").

***Ancestral polymorphism*—**The occurrence of a (largely?) degraded sister lineage of the 'Japonica O' type in the *crenata-sylvatica* lineage in addition to the I-B sister relationship is evidence for an ancient 5S rDNA polymorphism already present in the common ancestor(s) of all (Eurasian) beeches, i.e. the last common ancestor of Subgenus Fagus in Eurasia and Subgenus Engleriana (main-text fig. 8). 'European O' type arrays are probably much more common than covered in our data but not amplified because of high-pseudogenic gene regions with corrupted primer-attachment sequences; while their non-degraded counterparts in the *F. japonica* genome are much easier to capture using the here applied HTS approach (**Section 5.1**). According to fossil evidence and on the background of molecular phylogenies, this polymorphic ancestor can be considered to have lived at least ~35 myrs (cf. Grímsson et al., 2016) up to >50 myrs ago (Renner et al., 2016); *Fagus* 5S rDNA arrays have hence the capacity to retain independently evolving, alternative tribes of copies for tenth of millions of years.

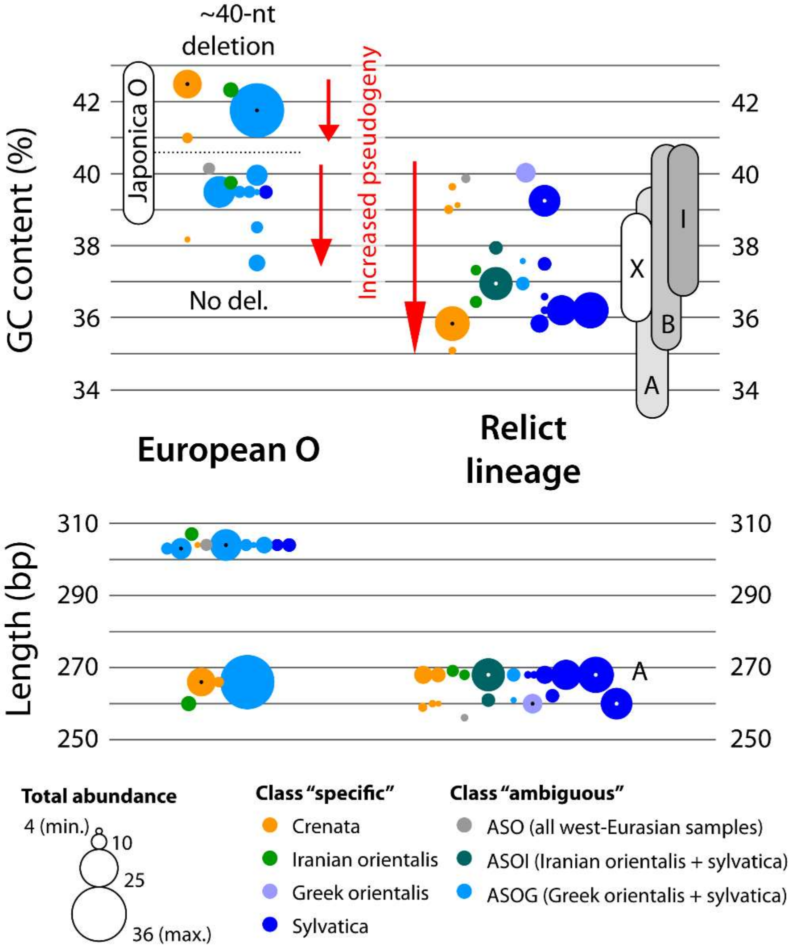

**Figure S14 | Scatter plots of GC content and sequence length of 'European O' and Relict Lineage variants.** Variants classified as "specific" are taxon-restricted, "ambiguous" are shared among studied samples. Bars give the GC range of normal-length variants (main length cluster, 305-bp length cluster in **Fig. S6**) for the major 5S-IGS types in *F. japonica* (representing I-, O-, and X-Lineage) and the *crenata-sylvatica* lineage (A- and B-Lineage).

The sole German-sample variant placed by EPA within the I-subtree (deeply nested; **Fig. S15**) is sequentially very distinct and lacks most (except for one point mutation) sequence patterns diagnostic for the I-Lineage ( "SyG3" in *Data S4*).[10] It includes sequence patterns otherwise

[10] This 'not-I' variant exemplifies the risk of mis-assignations when using automatic identification approaches such as EPA without post-analysis cross-check of surprising hits and visual inspection of results. The relatively low likelihood weight ratio for the best-placement alone is not conspicuous given the structure of our data and guide tree (many stochastically distributed mutations, flat terminal subtrees with high number of leaves); but in the case of the 'not-I' *sylvatica* variant, this it is coupled with a very long terminal branch in the EPA placement tree (all EPA placement trees are including in the *Online Data Archive*).

exclusive to the A- or B-Lineage, as well as shared, putatively underived oligo-nucleotide motives of the A- and I-Lineage. Subtracting point mutations due to beginning pseudogeny, the variant may well represent a leftover of the early diversification into I-, A- and B-Lineage or even an ancient recombinant.

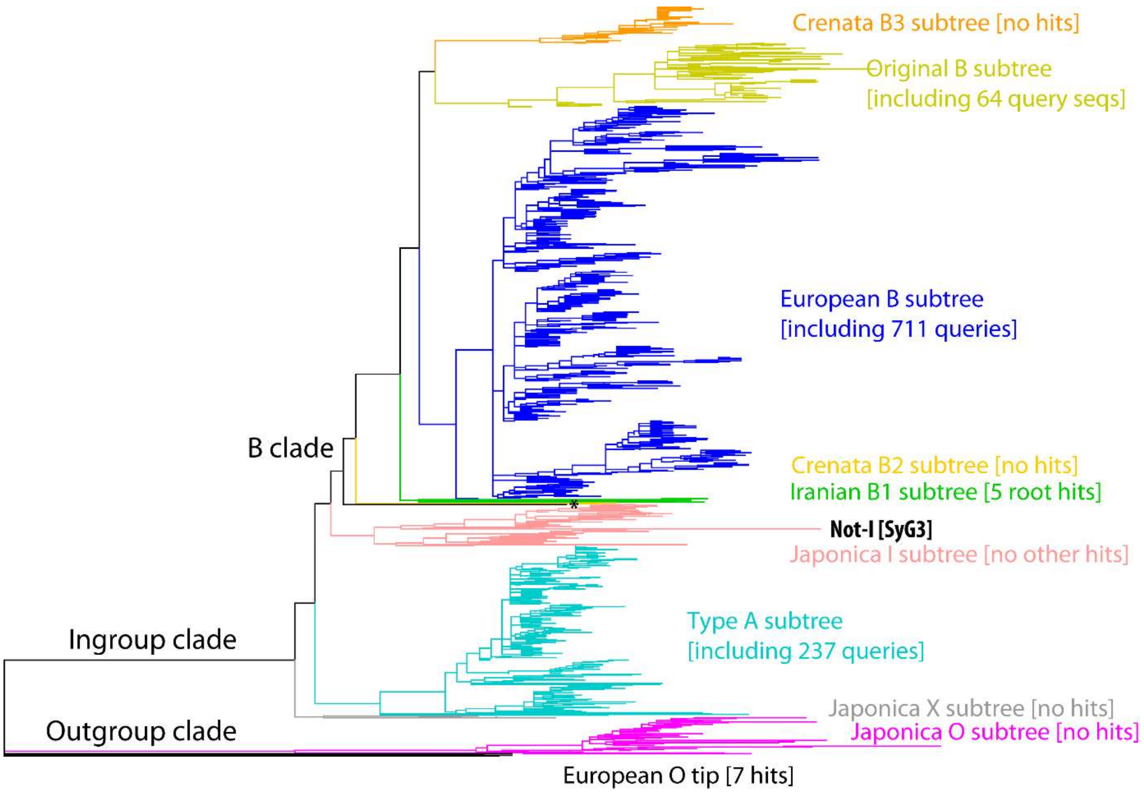

**Figure S15 | EPA placement tree for queries classified as 'Specific sylvatica'.** Shown is a phylogram, which gives the topology and branch lengths of the used reference tree (686-tip tree, cf. main-text fig. 3), and the tip-lengths of placed queries (cf. jPlace file included in the *Online Data Archive*). The distinctness of the single query placed within the I-Lineage subtree is obvious from such a graph; queries with much elongated tip branches may represent pseudogenic sequences, recombinants, or misplacements of unique sequence types not covered in the reference tree. * One query was placed at the root of the B-clade; with respect to the data and phylogenetic structure of the reference tree, such a placement can be indicative for an ancestral variant that lacks lineage-sorted and diagnostic sequence patterns.[11]

Ancestral polymorphism coupled with incomplete lineage sorting (and secondary mixing) is the only scenario to explain genetic relicts such as 'European O' and the not-I variant found in the German *F. sylvatica* s.str. sample. Sample contamination as alternative explanation can be ruled out insofar that the material was collected and DNA extracted at different times and the lack of identical variants in *F. japonica, F. crenata* and Iranian/ remaining European samples.

[11] When using EPA on short reads or queries with missing data, placements at roots can also indicate the query simply lacks the informative, lineage-discriminating sequence portions.

*Fagus japonica* is already rare in its native area, and extremely difficult to find in European arboreta or parks, and our west-Eurasian samples come all from natural (and old) beech stands. Hence, introgression because of cultivation can be ruled out as well. Notably, EPA did not recover any 'European O' variant in *F. japonica*, or 'Japonica O' and 'Japonica I' in any other sample.

***The Relict lineage*—**In addition, we captured rare variants that are sequentially intermediate between the 'outgroup' (O-Lineage) and the 'ingroup' types (X-, A-, B-, and I-Lineage) in all samples of the *crenata-sylvatica* lineage but not *F. japonica*. Two of these variants are shared by disjunct Iranian *F. orientalis* and *F. sylvatica* s.str. and samples (ASOI class in **Fig. S14**), one found in all west-Eurasian samples. Even more then the 'European O' variants, they can be enriched in potentially pseudogenous point mutations, in both the flanking gene regions and the non-coding, non-transcribed intergenic spacer. With respect to their phylogenetic position between 'outgroup' and 'ingroup', their pseudogenic tendency, and the oligonucleotide motives found in the 5' T-dominated LPR and the semi-homologous region, we refer to them as "Relict Lineage" (cf. main text). The pseudogenous mutations inflict long-branches in phylogenetic trees (long edges in networks). However, these stochastic patterns did not eliminate and obscure sequence patterns that appear to be conserved within and exclusive to this 5S-IGS shadow lineage (main-text fig. 5; *Data S4*). Hence, the high capacity of EPA to correctly identify members of this lineage based on only a single potential target in the 686-tip tree. Our hypothesis is that the Relict Lineage represents leftover imprints from early divergences or reticulations that much predate the formation of modern species, or involve species not sampled so far (continental and insular East Asian spp.; North American *F. grandifolia*) or extinct.

##### 4.4. Paralogy and possible homeology of 5S arrays in beech

The finding of two main sequence clusters in each sample (O- and I-Lineage in *F. japonica,* A- and B-Lineage in *crenata-sylvatica* lineage) coincides with the only available cytogenetic data of Ribeiro et al. (2011), showing *F. sylvatica* with two 5S rDNA pericentromeric loci (and four terminal 35S rDNA loci; arrays encoding for the 18S, 5.8S, 25S rDNAs). Two sequentially distinct, common and ± conserved length groups (**Section 3.1**) can point to genomic paralogy. In contrast to (topological) "paralogy" as used in phylogenetic literature, paralogy in a strict (genetic) sense implies that there are two (or more) 5S arrays per haplome. In polyploids and stabilized diploids, hom(o)eology (cf. Cronn et al., 2002), i.e. more than two 5S arrays per genome, has been observed as well in many plant groups (Yang et al., 2020; Piredda et al., 2021 and references therein), as well as in *Xenopus* (Ford & Southern, 1973; Cohen et al., 1999), bony fishes and elasmobranchs (Symonová, 2019). Homeologues are orthologues, i.e. they have the same locus (and function) but can evolve independently due to the lack of inter-haplome, inter-array concerted evolution. As consequence, homeologues inflict the same signal conflicts as paralogues, leading to the long-known issue of topological paralogy (e.g. Sanderson & Doyle, 1992; Ebach, 1999; Bailey et al., 2003). In beech, however, the situation is more complex than usually seen in (supposedly) diploid species and expected from cytology that identified two paralogous 5S rDNA arrays. Unless being a F1-hybrid or backcross, a diploid individual is expected to have two (more or less) homogenous haplomes. In the case of beech, diploid but with two loci per haplome, we thus would expect exactly two, potentially very distinct groups of 5S copies. In *F. japonica,* we observed two or three (X+I vs O), in the *crenata-sylvatica* lineage at least three (pseudogenous O-types, Relict Lineage, A+B).

In *F. japonica,* the longer, GC-rich and shorter GC-poorer length classes represent two highly divergent, distantly related lineages, the outgroup type O-Lineage and the ingroup type I-Lineage, close to the *crenata-sylvatica* lineage B-Lineage. Both are abundant; I- and O-Lineage variants are co-dominant. The basic hypothesis would be that *(i) F. japonica* has as well two paralogous 5S rDNA loci (per haplome), and (*ii*) each main lineage is linked to one of the loci. X-Lineage variants, representing a second 'ingroup' lineage of ambiguous phylogenetic affinity, may be rare but nevertheless exist and show no signs of sequence degradation, pointing either to a third locus, or intra-locus/intra-array mixing of 5S copies of different evolutionary sources (or representing different radiations). The data situation parallels cloned ITS data: ITS regions of *F. japonica* and its sisters *F. engleriana* (mainland China) and *F. multinervis* (Ulleung-do, Korea) show extreme length and sequence heterogeneity with up to three divergent main sequence types, while the ITS of the *F. crenata-sylvatica* lineage is more homogeneous and poorly sorted (Denk et al., 2005; Grimm et al., 2007a). This high ITS divergence corresponds with the cytological results of Ribeiro et al. (2011), who reported four NORs (Nucleolus Organizer Regions) for the diploid *F. sylvatica.*

In the 5S-IGS of the *crenata-sylvatica* lineage, the length and sequence differences are less pronounced with the longer A-Lineage variants being generally less derived and less abundant than the shorter much more abundant and more diverse B-lineage variants (*Data S5*). The samples exhibit increasing dominance of the B-Lineage over the A-Lineage: weakly developed in Iranian *F. orientalis*, increased in European samples following a (south)east/ (north)west gradient (Greece – Italy – Germany), and strongest in *F. crenata* (A-type sequences almost absent). In addition, putative relicts (cf. main-text fig. 5) can be found that lack/mix features of

A- and B-Lineage variants (*Data S5*) or are clearly related to the outgroup variants of the *F. japonica* O-Lineage ('European O', main-text figs 3, 4, 6; cf. **Section 4.3**). Hence, one can detect up to three major length classes referring to four principle 5S-IGS types of disparate phylogenetic affinity.

The divergence between the *F. japonica* ingroup (I-Lineage)/*crenata-sylvatica* (A-, B-Lineage) and outgroup (O-Lineage) variants would well fit with paralogy, i.e. independently evolving 5S rDNA array loci. Notably, although variants related to or with stronger affinity to the 'Japonica O'-type are occasionally found in the *crenata-sylvatica* lineage, they are extremely rare (cf. Table 2). A-Lineage and 'Original B' variants have been nearly eliminated in *F. crenata* and were replaced by three sequentially distinct, and *crenata*-specific B-Lineage types, mainly 'Crenata B2' (**Appendices A, B)**. Intragenomic silencing of paralogues (or homeologues, see below) leading to pseudogeny (reviewed in Volkov et al., 2007; see e.g. Volkov et al., 2017 for a case of an ancient allopolyploid) may cause the observed detection differences. The HTS primers bind to the highly conserved 5S rDNA. If these are strongly degraded, the intergenic spacers of such arrays will not be in our sample. Slightly degraded 5S array may be underrepresented.

While two paralogous 5S loci per haplome would facilitate retaining an ancient polymorphism (as the result of ancient polyploidisation/ hybridisation), i.e. O- vs X/I/A/B-Lineage, they cannot explain the diversity of the latter in the *crenata-sylvatica* lineage. In the light of the findings of Ribeiro et al. (2011), there are hence two possible scenarios: *(1)* the 'O' locus (or loci) is silenced in the *crenata-sylvatica* lineage; or *(2)* the A-/B-types replaced O-like arrays in both loci. Similarly, A-type arrays (or loci) were silenced or overprinted by B-types in *F. crenata* but not (yet) in its western Eurasian sisters; while in *F. japonica* a similar process eliminated X-Lineage variants in the I-X locus (loci). Based on the high structural and sequence diversity, counterbalanced by the large number of identical sequences detected in each sample, a combined effect of both concerted and birth-and-death evolution models must therefore be assumed for the 5S rRNA genes in beech (Nei & Rooney, 2005; Galián et al., 2014). Even if there are two (or more) loci in all species of *Fagus* (which are diploid, as far as studied), they may not be paralogous in a strict sense but act like homeologues, i.e. although they differ in position they do not differ in function and are affected by inter-array recombination and limited concerted evolution.[12] To date, little is known about the main features indicating 5S gene activity in non-model plants (e.g., copy number, GC content, secondary structure, promoter and terminator characteristics; Tynkevich & Volkov, 2019).

##### *4.5. Modern beech species as models for reticulate genetic mosaics*

including a re-evaluation of the 28-gene data compiled by Jiang et al. (2021)

Ancient reticulation, paralogy/homeology and sorting phenomena are also seen in other nuclear markers such as the internal transcribed spacers of the 35S rDNA (Denk et al., 2002, 2005; Grimm et al., 2007; see also graphs in Göker & Grimm, 2008, and Potts et al., 2014) and the 2ISP[13]-rich 28-gene data recently generated by Jiang et al. (2021; *Data S5*). To counter the high level of ambiguous base calls, Jiang et al. phased their data into a set of pseudo-OTU pairs ("…_1" and "…_2"; uploaded as independent accessions to gene banks) as basis for all analyses. The complexity of 2ISP patterns in their data – the full sequence details are provided in *Data S5* for both the uploaded phased reconstructed haplotypes (sheets '… rHT') and the de-phased original individual gene sequences (sheets '… 2ISP') – led to the reconstruction of a number of pseudo-haplotypes and artificial chimeric haplotypes (pseudo-recombinants; **Fig. S16**). In total, Jiang et al.'s (2021) reconstructed haplotypes include 16 cases, in which the two phased haplotypes of an individual belong to two different evolutionary lineages (genes P14, P21, P28, P48, P49[2 indiv.], F114[4], F202[5], F289). Furthermore, individuals of a species may have disparate phylogenetic affinities. Jiang et al. (2021) did not document the resulting intra-individual/intra-specific conflict as they collapsed all species' subtrees in their fig. 1.

[12] That we did not observe recombinant (chimeric) variants may be a sampling/data artefact. First, the pre-processing eliminates chimeric sequences. Second, 5S rDNA monomers are unlikely to show high proportions of chimeras because, in contrast to the ITS region, the non-transcribed intergenic spacer of the 5S rDNA has no sequentially highly conserved sequence patterns facilitating cross-over. In the ITS region, there are high-conserved sequence portions: the sequentially conserved ITS1 cleavage site, the extremely conserved 5.8S rDNA, and parts of the structurally conserved ITS2. Recombinant ITS clones were obtained e.g. in the case of densely sampled northern-temperate/-subtropical maple species (*Acer* sect. *Acer*; *Acer* sect. *Platanoidea*)
Grimm, G. W., Denk, T., & Hemleben, V. (2007b). Evolutionary history and systematic of *Acer* section *Acer* - a case study of low-level phylogenetics. *Plant Systematics and Evolution*, *267*, 215-253.
Grimm, G. W., & Denk, T. (2014). The Colchic region as refuge for relict tree lineages: cryptic speciation in field maples. *Turkish Journal of Botany*, *38*, 1050–1066.
[13] 2ISP = Intra-Individual Site Polymorphism, a general term introduced by Potts et al. (2014) for intra-individual sequence polymorphism irrespective of cause (heterozygosity, polyploidy, gene duplication) or mutation type (e.g. SNP – single-nucleotide polymorphism).

Phased data uploaded to gene banks: reconstruced haplotypes used by Jiang et al.

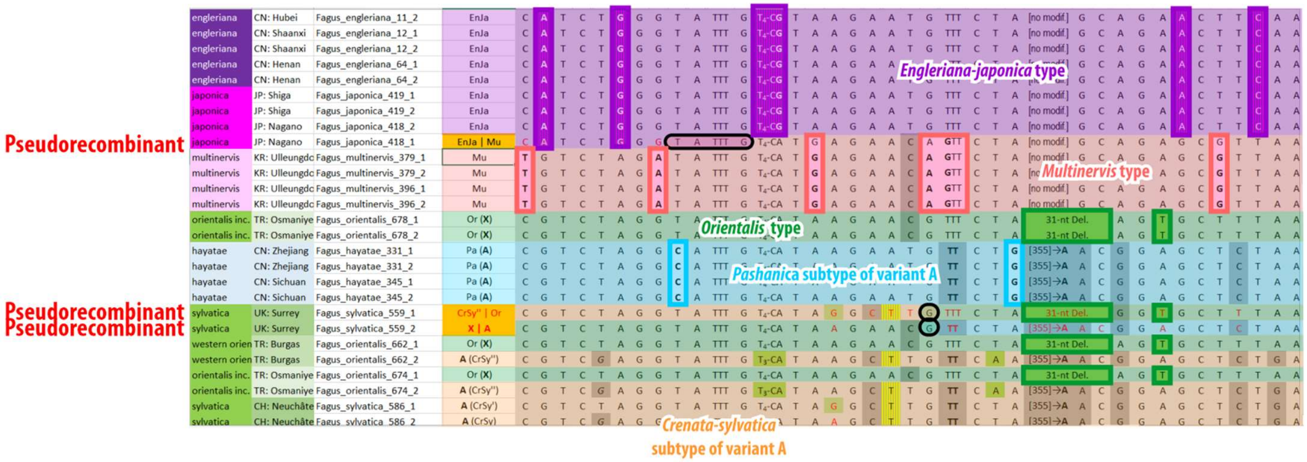

De-constructed original sequence data used for re-analysis,
generated by building the strict consensus of the two uploaded sequences per studied individual

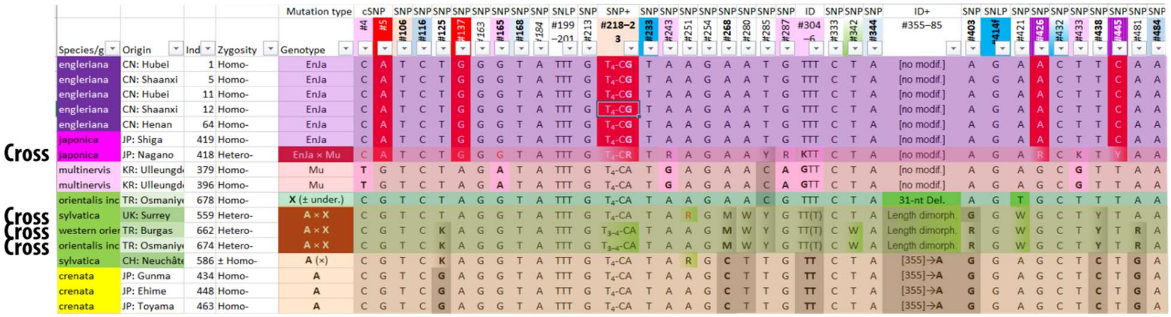

**Figure S16 | Example for pseudo-recombinants phased from 2ISP-rich heterozygotic/polymorphic individuals.** West-Eurasian individuals (green background) are characterized by a pronounced intra-individual gene polymorphism combining two major gene variants, Variant X close to the genus' consensus sequence, and the more evolved variant A shared across Eurasia; obviously, the west-Eurasian ancestor was polymorphic. One *japonica* individual (light purple) is a genetic cross between the *engleriana-japonica* and *multinervis* types. Part of the *multinervis* signature was not detected, or concerted evolution and backcrossing with pure *japonica* has started to homogenize the gene pool. By phasing such data one inflicts increased intra-specific divergence and artificially inflates terminal branch length. The concatenation (combination) with sorted data will however overprint such genes' internal conflict, masking it entirely in a combined tree (Jiang et al., 2021, fig. 1; cf. **Fig. S18** below).

The failure of algorithmic phasing is partly due to potentially paralogous gene copies (>2 gene variants per genome), i.e. 2ISP patterns that go beyond simple heterozygosity (only two variants, haplotypes, per genome). Evidence for within-lineage and incompletely sorted potential gene duplications can be found in genes P14, P67, F202 (**Fig. S17**), and F289. While the conflicting signals of potential paralogues are problematic for both classic and coalescence tree inferences, they can be an invaluable source to detect past reticulation (discussed in main text).

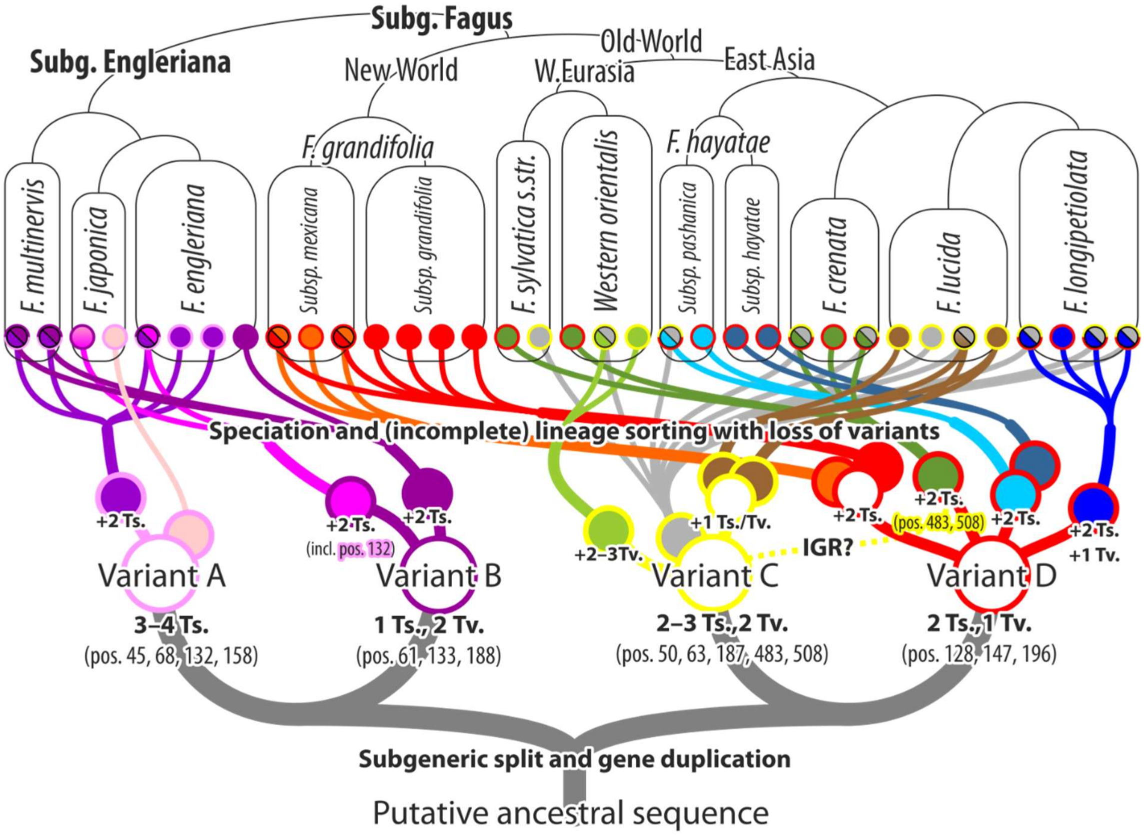

**Figure S17 | Example for potential gene duplication and subsequent partial lineage sorting involving gene loss in Jiang et al.'s gene F202.** Two main sequence variants characterize both subgeneric lineages (A+B: subg. Engleriana, C+D: subg. Fagus) and lead to high intra-generic diversity poorly sorted along the inferred combined-data species tree (top brackets; cf. **Fig. S18**). Abbrev.: IGR – possible intra-genomic recombination of Variants C and D in precursors of the *crenata-sylvatica* lineage; Ts. – transitions; Tv. – transversions.

Jiang et al.'s reconstructed haplotypes (or selected subset thereof) are problematic for inferring a species tree (Jiang et al., 2021, fig. 1) or a multi-species coalescence tree (Jiang et al., 2021, fig. 2). Nonetheless, the primary signals from their individual-level sample covering all but one species (Eastern *orientalis*) can provide additional insights into the complex genetic history of modern beeches, i.e. by allowing us to put our species-wise limited 5S-IGS data in a larger context (see main text). By overlapping the phased reconstructed haplotypes and building the strict consensus sequence using G2CEF (Göker & Grimm, 2008), we first deconstructed the original, 2ISP-rich gene sequence data (data files included in Grimm, 2020). This allowed us to identify and classify gene-polymorphic individuals (*Data S5*) and use the data to infer individual-based trees, i.e. using biological entities as OTUs instead of algorithmic constructs. In a second step, we used this tree to map individual gene differentiation patterns (**Fig. S18**). The overall divergence is relatively low in all genes (15–47 parsimony-informative sequence patterns[14]; median of 26), hence one could infer (full) median networks or median-joining networks for nearly all of the 28 genes (except for gene P67[15]), thus, identifying (still present) ancestral vs. derived (species- or lineage-specific) genotypes as well as heterozygotic individuals combining both genotypes (black-rimmed in **Fig. S18**).

Overall, while the combined tree finds the same inter-species relationships, it produces less biased branch-lengths and even higher overall support (only two branches above species-level with BS < 100) as polymorphic OTUs are treated as such and not divided into two tips, phased reconstructed haplotypes treated as independent taxa. Like the original tree based on the reconstructed haplotypes, the individual-based tree includes false positives, i.e. branches with high, (near-)unambiguous support that are gene-wise not unambiguous. Largely ignored in phylogenomic studies until today (but see Delsuc et al., 2005), the combination of incongruent genes can lead to topological artefacts. In extreme cases, strongly incongruent data, few tips, the combined tree may include clades not supported by a single gene/character in the underlying matrix (see e.g. example used in Schliep et al., 2017). Dominant genes (here: individual splits) may erase any conflict in the rest of the gene sample and can result in inflated branch support (as in the case of Jiang et al., 2021, data; see below). For instance, gene-wise analysis of complete angiosperm plastomes revealed that the high-supported, fully resolved complete plastome tree essentially receives its support from only very few of the plastid genes (mainly *rpoC2, matK*) and potentially masks disparate gene histories (Walker et al., 2019).

---

[14] Most parsimony-informative sequence patterns are SNPs or single-site 2ISPs. A few include linked mutations and oligonucleotide motives (incl. length-polymorphism).

[15] The reconstructed haplotypes documented by Jiang et al. (2021) are signal-wise chaotic, and de-phased into the likely original sequences show a highly diffuse 2ISP pattern with no apparent phylogenetic structure. It would be obligatory to clone the P67 PCR products to extract the actual gene variants (possible paralogues) leading to the found 2ISP pattern. Based on Jiang et al.'s method chapter, it is impossible to reconstruct whether detection or amplification issues may have underestimated the real number of 2ISPs in the sampled genomes. From the sequence-wise comparison, it appears that some 2ISPs remained undetected, especially in genes with potential paralogues/more than one major sequence variant per lineage/individual.

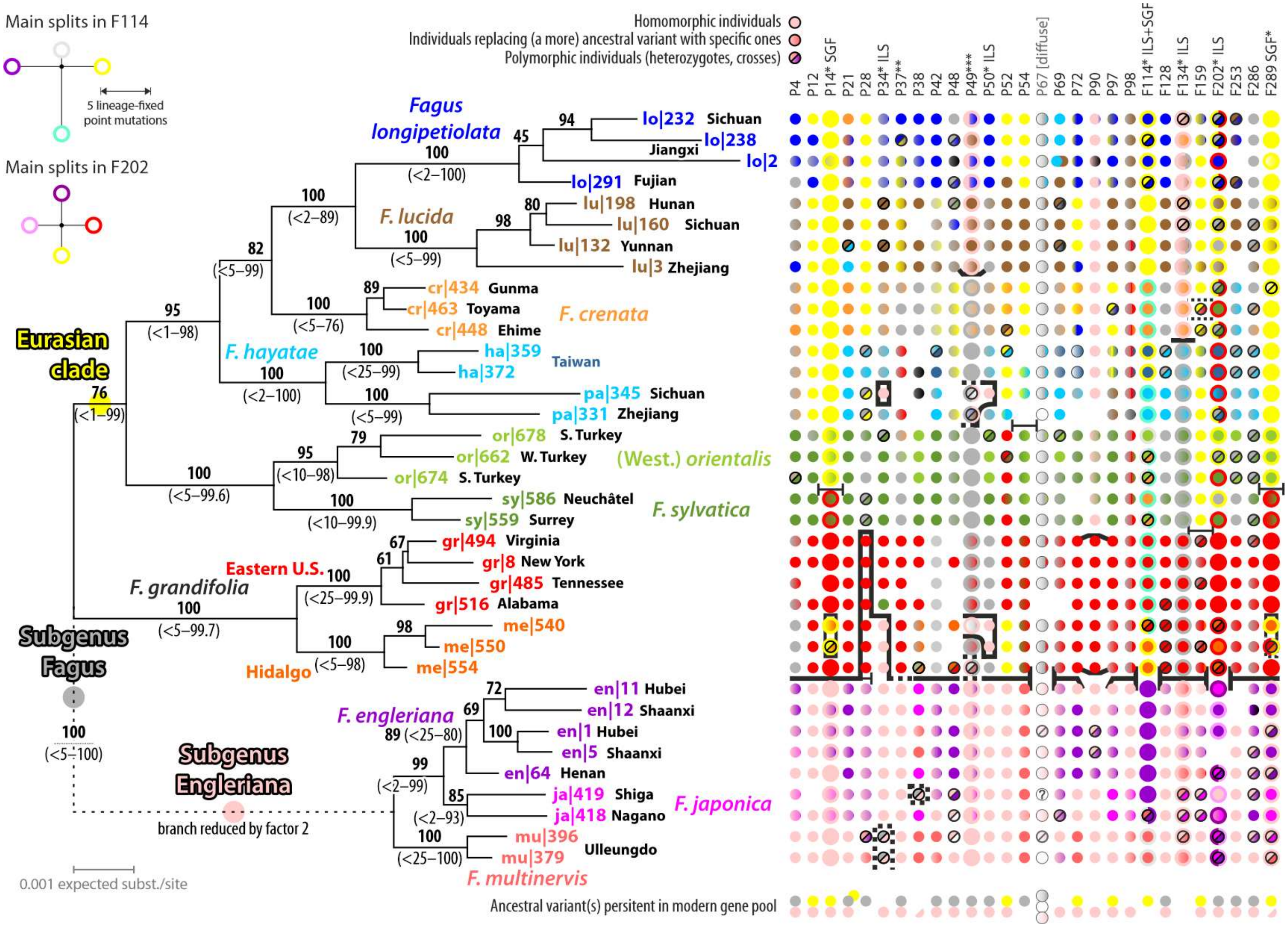

Main splits in F114
5 lineage-fixed point mutations
Main splits in F202
Homomorphic individuals
Individuals replacing (a more) ancestral variant with specific ones
Polymorphic individuals (heterozygotes, crosses)
P4 P12 P14* SGF P21 P28 P34* ILS P37** P38 P42 P48 P49*** P50* ILS P52 P54 P67 [diffuse] P69 P72 P90 P97 P98 F114* ILS+SGF F128 F134* ILS F159 F202* ILS F253 F286 F289 SGF*
Fagus longipetiolata
lo|232 Sichuan
lo|238 Jiangxi
lo|2
lo|291 Fujian
F. lucida
lu|198 Hunan
lu|160 Sichuan
lu|132 Yunnan
lu|3 Zhejiang
cr|434 Gunma
cr|463 Toyama
cr|448 Ehime
F. crenata
F. hayatae
ha|359
ha|372
Taiwan
pa|345 Sichuan
pa|331 Zhejiang
or|678 S. Turkey
or|662 W. Turkey
or|674 S. Turkey
(West.) orientalis
sy|586 Neuchâtel
sy|559 Surrey
F. sylvatica
gr|494 Virginia
gr|8 New York
gr|485 Tennessee
gr|516 Alabama
Eastern U.S.
F. grandifolia
me|540
me|550
me|554
Hidalgo
Eurasian clade
Subgenus Fagus
en|11 Hubei
en|12 Shaanxi
en|1 Hubei
en|5 Shaanxi
en|64 Henan
F. engleriana
ja|419 Shiga
ja|418 Nagano
F. japonica
mu|396
mu|379
Ulleungdo
F. multinervis
Subgenus Engleriana
branch reduced by factor 2
0.001 expected subst./site
Ancestral variant(s) persitent in modern gene pool

**Figure S18 (preceding page)| Species tree (left), inferred from the combined 28-gene data, and individual gene map (right);** using sampled individuals as tips (OTUs). Note the generally high, (nearly) unambiguous support for all branches, which contrasts the substantial inter-gene incongruence and sharing of ancestral (pink – ancestral within subg. Engleriana; grey – within subg. Fagus; yellow – within the Eurasian clade of subg. Fagus) or derived gene variants, which can be specific for sister species/lineages (all other colours). Lines in the gene map give the primary (typically subgeneric) and secondary phylogenetic split (two main gene variants in subg. Fagus) observed in each gene. Stippled lines bracket individuals with 2ISPs covering two main variants. Numbers at branches give ML(-A) BS support (cf. Potts et al., 2014) from the combined data and, in bracket, from gene-wise bootstrapping (ranges). * Genes with major sequence variants not sorted along the putative species tree (ILS – incomplete lineage sorting; SGF –secondary gene flow/ lineage-crossing); ** only the North American genotype (red) is substantially different from the ancestral (subgenus-consensus) sequence; *** gene with few but consistent mutations, largely decoupled from phylogeny.

Detailed sequence-wise re-inspection of Jiang et al.'s (2021) individual-level data (*Data S5*) shows that many 2ISPs patterns are lineage-diagnostic. All mutation patterns (mostly SNPs) involving 2ISPs include the generic (or subgeneric) consensus state in addition to, often species-unique, mutations. Adding to the lineage- and species-conserved mutation patterns and a generally low fixation rate, one can hence discern ancestral shared (pink, grey and yellow in **Fig. S18**) and potentially specific, derived genotypes (all other colours). The characteristic 2ISP patterns further allow pinpointing ancient and more recent crosses: individuals that are heterozygotic or polymorphic, in case of gene duplication, for distinct lineages. Mapping the so-identified gene variants on the tree demonstrates that some high-supported relationships seen in the combined tree are likely inference artefacts, resulting from forcing multi-dimensional non-treelike data, the result of reticulate evolutionary processes (discussed in main-text) into a single-dimension graph, a tree. Rather than approximating the coalescent, the combined tree merely represents the minimum-conflicting topology for an extremely complex data situation and genomic history (**Section 4.6**).

Mirroring what has been found for non-coding ITS (Denk et al., 2002, 2005; Grimm et al., 2007) and 5S-IGS data (main-text), the nuclear diversity and intra-genomic variation in Jiang et al.'s (2021) 28-gene data reflects gradual, population-scale as well as speciation and isolation processes: ancestral gene variants are (slowly) replaced by specific ones in the gene pool of modern *Fagus* species. Likewise, there is compelling, and easy to extract and visualize, evidence for ancestral and more recent (local) gene flow/gene exchange, in addition to incomplete lineage sorting involving potential gene duplications. Genes showing two or more major sequence types, potential paralogues, are sorted in some beech lineages/species while mixing in others indicating a complex history of gene duplication and loss in the evolution of the nucleome of modern *Fagus* species (**Fig. S19**).

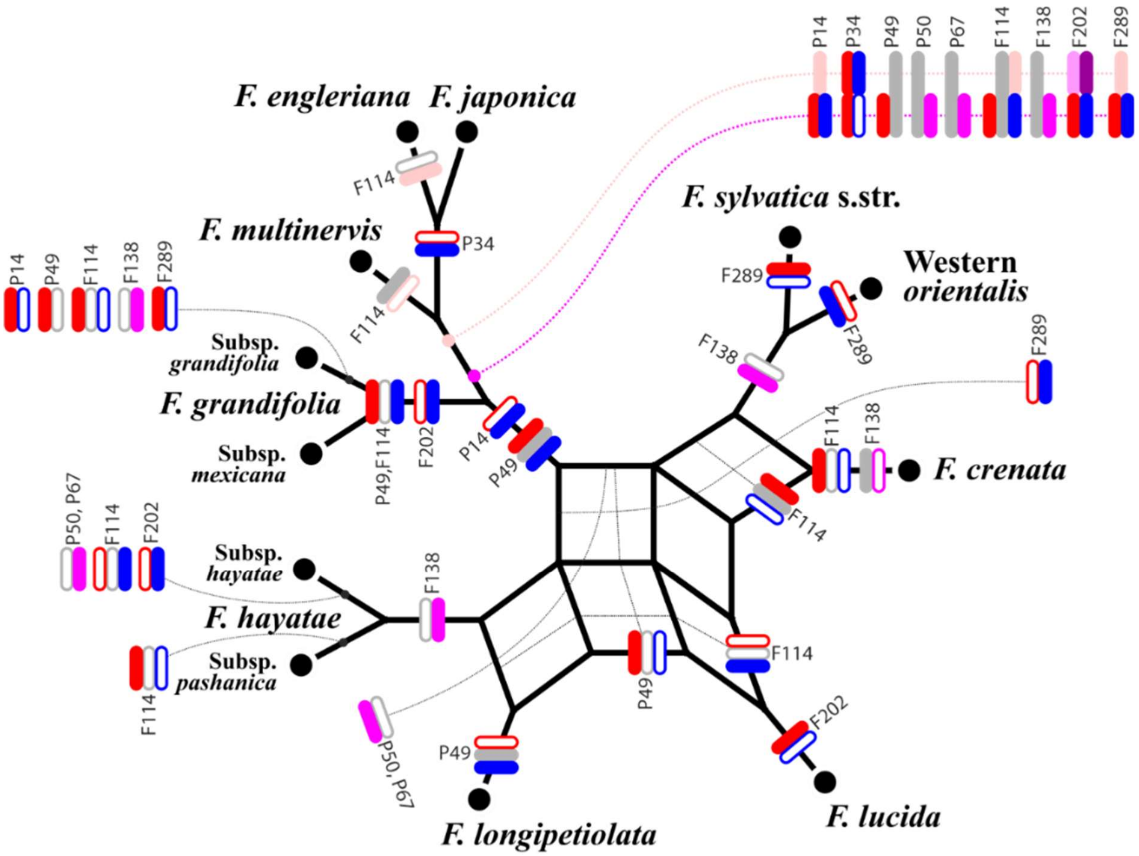

**Figure S19 | Gain and loss of major gene sequence variants mapped on the current synoptical species network (update of in-text fig. 1).** New major sequence variants may origin from gene duplication, or could have been captured during introgression/hybridisation. Full signature represents gain, open signature a loss of a major variant within a species' gene pool. Note that no intra-specific variation is shown; typically, if more than one variant is present, there are homozygotic/homomorphic individuals in addition to heterozygotic/polymorphic ones.

**(Following page) Figure S20| Gene incongruence in the 28-gene data of Jiang et al. (2021)**. Shown is a super-network of all 28 gene trees[16]; edge-lengths are sums of individual gene tree branch lengths. Numbers at selected edges give the range of gene-wise bootstrap support: the maximum and, in brackets, the median of all genes with BS > 25 for the according split (if supported by 6+ genes with BS > 25), the minimum[17], and the minimal averaged support across all 28 genes (splits with BS < 25 counted as 0). The light pink edge ("Artef.") is a reconstruction artefact (long-branch attraction between an aberrant lo|2 genotype and Subgenus Engleriana).

---

[16] Generated using 100 runs of the z-closure algorithm (Huson et al., 2004. Phylogenetic super-networks from partial trees. *IEEE/ACM Trans Comput Biol Bioinform* 1:151–158) implemented in SplitsTree. Being limited to maximal four dimensions, the graph does not show all conflicting splits in the gene trees.

[17] We only recorded splits with a BS ≥ 25 (*Data S5,* sheet 'GenewiseSplitSupport') based on accordingly filtered support consensus networks (cf. Schliep et al., 2017) of the bootstrap pseudoreplicate samples inferred for each gene. A minimum BS < 5, e.g., means that this alternative was rejected by at least one gene with a conflicting split receiving BS ≥ 95.

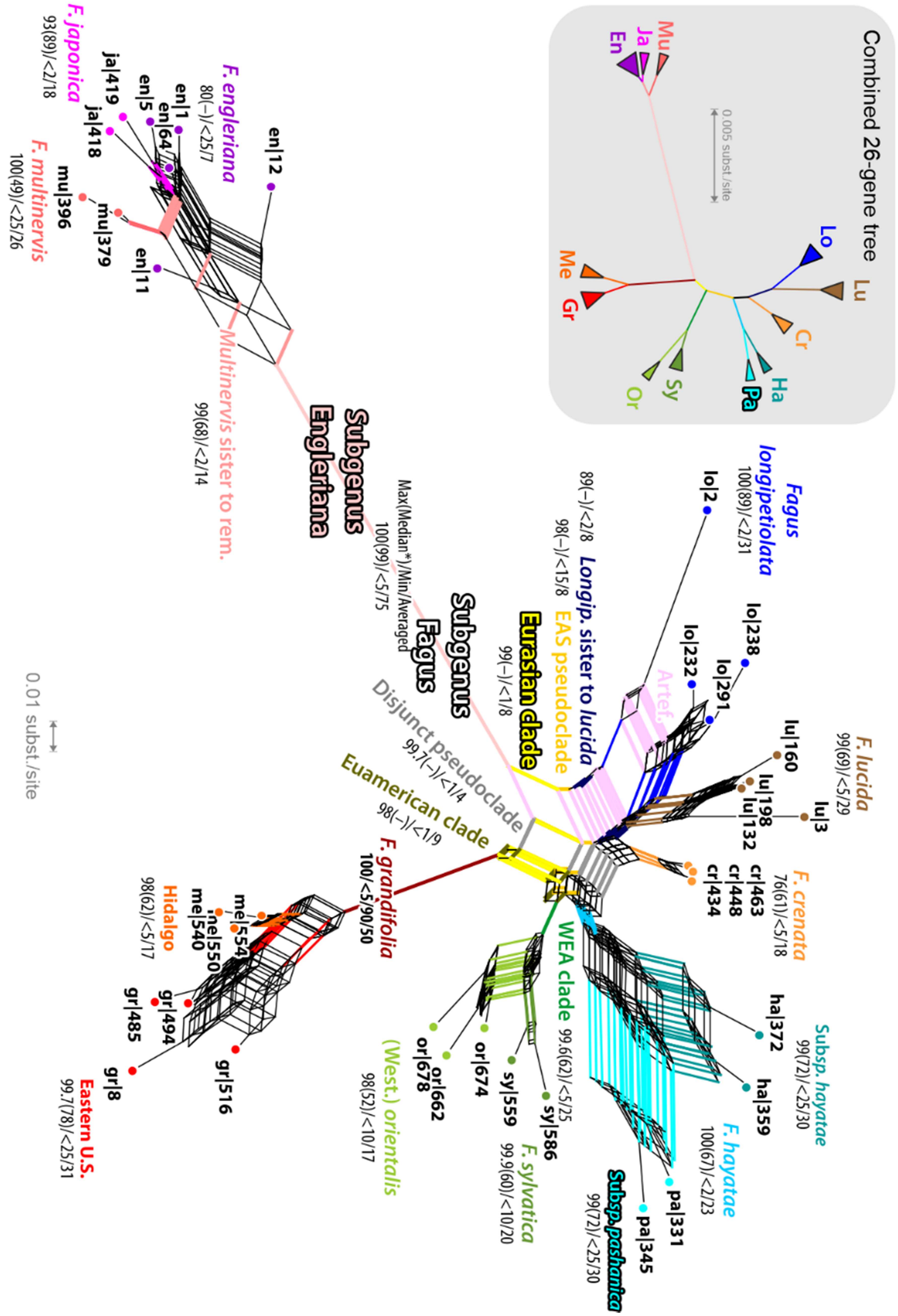

Combined 26-gene tree
0.005 subst./site
Mu
Ja
En
Lo
Lu
Cr
Ha
Pa
Sy
Or
Me
Gr
0.01 subst./site
Subgenus Engleriana
Multinervis sister to rem.
99(68)/<2/14
Max(Median*)/Min/Averaged
100(99)/<5/75
Subgenus Fagus
Fagus longipetiolata
100(89)/<2/31
89(–)/<2/8
98(–)/<15/8
Longip. sister to lucida
EAS pseudoclade
Eurasian clade
99(–)/<1/8
Artef.
Disjunct pseudoclade
99.7(–)/<1/4
Euamerican clade
98(–)/<1/9
F. grandifolia
100/<5/90/50
F. lucida
99(69)/<5/29
F. crenata
76(61)/<5/18
Subsp. hayatae
99(72)/<25/30
F. hayatae
100(67)/<2/23
Subsp. pashanica
99(72)/<25/30
WEA clade
99.6(62)/<5/25
F. sylvatica
99.9(60)/<10/20
(West.) orientalis
98(52)/<10/17
Hidalgo
98(62)/<5/17
Eastern U.S.
99.7(78)/<25/31
F. engleriana
80(–)/<25/7
F. japonica
93(89)/<2/18
F. multinervis
100(49)/<25/26

##### *4.6. A preliminary species network for beech trees*

The utility of phylogenomic data to infer a species tree for beeches (Jiang et al., 2021) is strongly limited because beech evolution is not a linear-dichotomous process but a complex reticulate one, and especially has been so in the genus' past. Moreover, the sorting of genes after speciation is strongly affected by population-level processes: phases of introgression and unhindered gene flow facilitated by wind-pollination and beech ecology (see main-text) alternate with phases of isolation and increased genetic drift. Beech species perpetually evolve and fuse. Consequently, individual gene genealogies may differ profoundly (**Fig. S20**; not documented in Jiang et al., 2021) being the product of incomplete lineage sorting *and* (past) lineage mixing. When examined in detail, the data can be used to extract a general picture of gene sorting and mixing among the precursors of modern beech species adding to initially generated, broad-sampled ITS (Denk et al., 2002, 2005) and newly generated 5S-IGS data (**Fig. S21**).

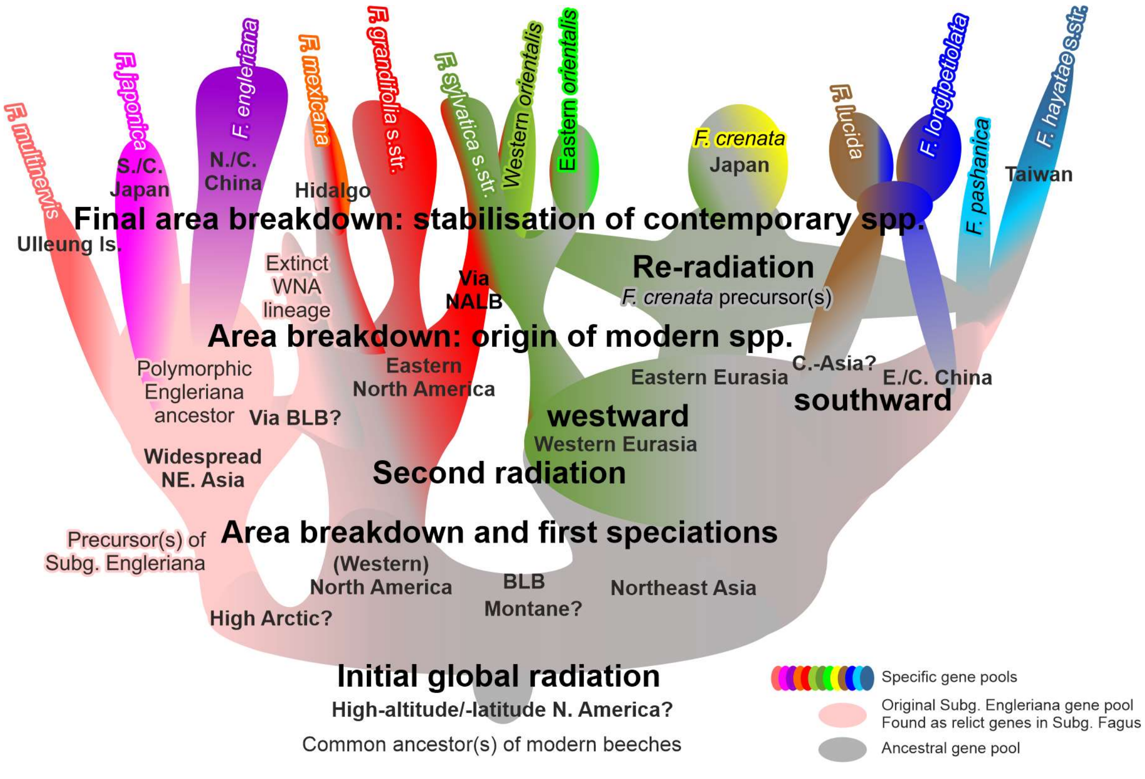

**Fig. S21 | Summarizing doodle depicting the evolution of the nuclear gene pool in *Fagus*** based on the 28-gene data of Jiang et al. (2021) and including evidence from broad-sampled ITS, 5S-IGS and available *LEAFY* intron 2 data (Denk et al., 2002, 2005; Grimm et al. 2007; Oh et al., 2016; Renner et al., 2016; Worth et al., 2021)

***Initial subgeneric split*****—**As in the case of ITS and the here reported 5S-IGS data, the subgeneric boundaries are well reflected in most genes of Jiang et al. (2021). The Japanese *F. japonica* shows Subgenus Engleriana-type sequences and *F. crenata* (typically slightly derived) Subgenus Fagus-type sequences, and there is no evidence for lineage mixing. Similarly, no derived sequence types are shared between *F. engleriana* and the Chinese species of Subgenus Fagus, despite overlapping distribution areas. Individual genes of the two subgenera differ by 1–11 subgenus-conserved mutations (median = 4; in total 112 mutations). Exceptions are genes P34, P49, P50, P67 (generally diffuse signals), P90, F114, F138, and F202 (**Table S3**). In these genes a potential gene duplication seems to have preceded the subgeneric split or the sampled gene region is simply too conserved to resolve any phylogenetic relationships at the genus-level and below. Being the second-most consistently supported split (min. averaged BS support of 75), the subgeneric split likely represents a very early phase of *Fagus* evolution (**Fig. S21**; cf. Renner et al., 2016). Subgenus Engleriana-similar genotypes found in members of Subgenus Fagus seem to represent relicts inherited from the genetically heteromorphic, widespread common ancestor(s) of all beeches, or gene flow at a very early stage in *Fagus* evolution (after the first speciation phase). All non-coding nuclear markers sequenced so far were always strictly sorted by subgenera and sequentially most distinct (see also **Section 4.4**). In the light of modern and past distribution, Subgenus Engleriana represents the tip of an old and stable high-latitude, Northeast Asian/Beringian lineage of beeches (cf. Denk & Grimm, 2009).

***New World beeches, a deep split*****—**Based on ITS and nuclear-intron data it has always been clear that the New World beeches represent an old, early evolved and largely isolated lineage within Subgenus Fagus (Denk et al., 2005; Renner et al., 2016). This is corroborated by Jiang et al.'s (2021) 28-gene data. The (eastern) North American individuals usually differ by several (1–5; in total 36) lineage-conserved mutations from the rest of Subgenus Fagus; in addition to mutations/ 2ISPs only found in some of the New World individuals or diagnostic (up to 3) for the two disjunct infraspecific taxa (subsp. *mexicana* in central Mexico, *F. grandifolia* s.str. in the Appalachian mountain range and adjacent lowlands). In some genes, the New World genotypes are substantially closer to Subgenus Engleriana than the rest of Subgenus Fagus (**Table S3**), in perfect agreement with the fossil record (Denk & Grimm, 2009).

**Table S3 | Most prominent splits in genes not sorted primarily by subgenera.** Split taxa – species, continental (subsp. *pashanica*) and insular (subsp. *hayatae*) subspp. of *F. hayatae* (*F. h.*), *F. sylvatica* s.l. – highlighted in red font.

<table>
<tr><th>Gene(s)</th><th>BS support</th><th colspan="4">Main splits</th></tr>
<tr><td>P14</td><td>94</td><td colspan="2">Subg. Engleriana + F. sylvatica</td><td colspan="2">All other subg. Fagus spp. incl. (western) orientalis</td></tr>
<tr><td></td><td>86 | 100 | 87 | 67</td><td>Subg. Engleriana</td><td>F. sylvatica</td><td>cr|434, F. hayatae, F. longipetiolata, F. lucida</td><td>cr|448, cr|436, (western) orientalis, N. American spp.</td></tr>
<tr><td>P28, F159</td><td>99, 34[a]</td><td colspan="2">Subg. Engleriana + N. American spp.</td><td colspan="2">Eurasian spp. of subg. Fagus</td></tr>
<tr><td>P34, P50</td><td>91, 95</td><td colspan="2">Subg. Engleriana incl. F. mexicana</td><td colspan="2">F. grandifolia s.str. + Eurasian subg. Fagus spp.</td></tr>
<tr><td>P38</td><td>94</td><td colspan="2">Subg. Engleriana incl. ja|418</td><td colspan="2">Subg. Fagus + ja|419</td></tr>
<tr><td>P49</td><td>85</td><td colspan="2">Subg. Engleriana, F. longipetiolata, F. lucida, me|540, pa|345</td><td colspan="2">F. crenata, F. h. subsp. hayatae, pa|331, west-Eurasian spp., N. American spp. (incl. me|550, me|554),</td></tr>
<tr><td>P67</td><td>93</td><td colspan="2">Subg. Engleriana incl. pa|331</td><td colspan="2">Subg. Fagus (incl. pa|345)</td></tr>
<tr><td>P90, F202</td><td>99, 87</td><td colspan="2">Subg. Engleriana mixed with Eurasian spp. of sg. Fagus</td><td colspan="2">N. American spp.</td></tr>
<tr><td>F114</td><td>30</td><td colspan="2">Subg. Engleriana incl. or|678</td><td colspan="2">Subg. Fagus (incl. or|662, or|674)</td></tr>
<tr><td></td><td>80 | 99 | 57 | 56</td><td>F. multinervis + or|678</td><td>F. engleriana + F. japonica</td><td>F. mexicana +{ F. h. subsp. hayatae, F. longipetiolata; F. lucida }</td><td>F. h. subsp. pashanica +{ or|662, or|674, F. crenata, F. grandifolia s.str., F. sylvatica }</td></tr>
<tr><td>F138</td><td>99.7</td><td colspan="2">Subg. Engleriana, F. crenata, F. lucida, F. longipetiolata</td><td colspan="2">N. American spp. + { west-Eurasian spp. + F. hayatae }</td></tr>
</table>

[a] The low support in case of gene F159 is due polymorphic patterns in one *F. crenata* individual (cr|463), being heterozygotic for the ancestral N. American and ancestral Eurasian genotypes. Accordingly it is placed as sister to the N. American spp. in the BS consensus network but included in the (overall less divergent) Eurasian clade in the 'best-known' ML gene tree. A Eurasian clade excluding cr|463 would receive BS = 88.
[b] A subgeneric split received BS = 51; ja|419 is, like cr|463 for gene F159 (→ footnote a), heterozygotic with 2ISPs covering the ancestral genotypes in both subgenera.

A Beringian (subg. Engleriana + ancestors of *F. grandifolia* s.l.)–Eurasian split (remainder of subg. Fagus) can be found in genes P28 and F159, supported by a total of seven mutations. In contrast, the East Asian clade discussed below only received support from one mutation each in three genes. The Mexican individuals (*F. mexicana*) are either closer to the consensus of the Eurasian Subgenus Fagus species or show conspicuous 2ISP patterns, pointing towards ancestral intra-genomic polymorphism (i.e. ancient hybridisation/ introgression) lost in the northern *F. grandifolia* s.str. Based on their geographic position, it is not unlikely that *F. mexicana* has also conserved genetic signatures from the extinct north-western North American beech species that must have been much closer to the ancestor(s) of the Eurasian spp. of Subgenus Fagus or all *Fagus*, and may have exchanged genetic material with the precursors of Subgenus Engleriana and other high-latitude Eurasian species across the Beringian land bridge.

***Gene flow via North Atlantic land bridge***—In three genes (P21, P54, F289; total of eight mutations; Euamerican clade in **Fig. S20**), the data point towards a direct cross-Atlantic connection: highly similar, putatively derived sequences are shared by (part of) North American and west-Eurasian individuals. Gene flow via the so-called North Atlantic Land Bridge has been possible at least until the Miocene (Grímsson & Denk, 2005, 2007; Denk et al., 2011). New World genetic signatures[18] may have intrograded into the *F. haidingeri* gene pool, the precursor species of all west-Eurasian species, via hybridisation with/ introgression from *F. gussonii* (see also Denk, 2004). A vector species of North American affinity like *F. gussonii* also would explain why there is no indication for specific west- or north-Eurasian gene variants having intrograded into the New World species.[19]

***The "East Asian Clade", a paraphylum***—Jiang et al.'s (2021) high-supported "East Asian Clade" (see also **Fig. S18**) draws its support largely from the generally less-evolved, less species-coherent gene sequences in the East Asian spp. compared to their west-Eurasian and North American siblings ('short-branch culling', a form of long-branch attraction, LBA).

[18] Cf. Denk et al. (2002), documenting a pseudogenous N. American ITS copy in a Caucasian beech

[19] The genetic links to Subgenus Engleriana and (part) of the west-Eurasian individuals are the reason the *BEAST multi-species coalescent preferred a cross-North Atlantic clade and placed *F. grandifolia* (s.l.) as sister to the west-Eurasian species. While in the combined tree, any such topology must be rejected because such splits are incompatible with the dominant differentiation patterns in most genes showing a general similarity of all Eurasian species, their substantial character-support (higher than for any other deep-split alternative) triggers the *BEAST-inferred topology, which is essentially based on single-gene optimisations (see also **Fig. S20**).

Notably, derived mutations are typically shared only between individual species pairs but never the entire clade:

- *F. crenata* + *F. hayatae* (continental and/or insular subspp.);
- *F. crenata* + *F. lucida,* in agreement with ITS polymorphism (Grimm et al., 2007);
- *F. hayatae* (both subspp.) + *F. longipetiolata* (cf. Denk et al., 2005); and
- *F. longipetiolata* + *F. lucida*; a split also seen in *LEAFY* intron data (Oh et al., 2016).

Direct support for an inclusive common origin is rare: the entire 28-gene data include three point mutations scattered across three genes (P12: A→T at pos. 317, missing in *F. longipetiolata* from Fujian; P21: G→T#643; P54: T→C#715), compared to seven/eight supporting conflicting splits not seen in the combined tree (**Table S4**).[20]

**Table S4 | Character support for major intra-generic splits seen in the combined tree and most consistently found alternatives**; sorted by total number of supporting mutation patterns.

| Split | Conflicting with combined tree | Number of genes with fitting mutations | Supporting, conserved mutations | |
|---|---|---|---|---|
| | | | Per gene | Total |
| Subgeneric split | No | 25 | ≤ 11 | 112 |
| *F. grandifolia* s.l. (N. American spp.) vs. rest | No | 24 | ≤ 5 | 36 |
| West-Eurasian spp. vs. rest | No | 11 | ≤ 6 | 18 |
| *F. hayatae* (continental + insular subspp.) vs. rest | No | 10 | ≤ 4 | 11 |
| Atlantic (N. American + west-Eurasian) \| Pacific | Yes | 3: P21, P54, F289 | 1 or 6 | 8 |
| *F. engleriana* + *F. japonica* vs. rest | No | 3: P21, F114, F289 | 1/2/5 | 8 |
| Cross-Beringia (subg. Engler. +*F. grandif.* s.l.) \| Eurasia | No | 2: P28, F159 | 3/4 | 7 |
| East Asian spp. vs. rest | No | 3: P12, P21, P54 | 1 | 3 |

Regarding the standard (cladistic) interpretation of molecular trees (clade = monophylum), the East Asian clade is obviously a false positive (hence, labelled 'East Asian pseudoclade' in **Fig. S20**). The inconsistency of the associations, changing from gene to gene, in addition to the observation that all four East Asian species can share (± ancestral) sequence types with at least some western Eurasian or Mexican individuals (cf. **Fig. S17**) points towards a complex interplay between genetic drift in the course of speciation, gene flow via secondary contact – the ancestors of *F. crenata* were probably much more widespread than today (cf. Denk & Grimm, 2009) – and stochastic sorting in shaping the modern gene pools of all Eurasian species. An inclusive common origin (monophyly fide Hennig; holophyly fide Ashlock) of the East

[20] Vice versa, these mutations would notably support the alternative of a high-latitude link between Subgenus *Engleriana*, *F. grandifolia* and the west-Eurasian species. In fact, any East Asian vs. rest split may be a left-over imprint from an early differentiation in high-latitude and mid-latitude species that predates the formation of all modern species (cf. Denk, 2004; Denk & Grimm, 2009); in particular with respect to upcoming *CRC, LFY* and plastome data (Worth et al., 2021)

Asian spp. can be ruled out. If their radiation occurred after the divergence between the west-Eurasian and East Asian members of Subgenus Fagus as inferred by Jiang et al. (2021) and our combined tree using their unphased data (**Fig. S17**; but see **Fig. S20**), we should find many more segregating mutations supporting an East Asian ingroup (crown group) vs outgroup (western Eurasian spp. + North American spp. + subg. Engleriana) split. Sequence types in both western Eurasian and East Asian spp., the mutual sister lineages, should be independently derived from a hypothetical ancestor, which should be genetically closer to the North American spp. than its western Eurasian and East Asian descendants. However, all genes that show high phylogenetic structuring comprise North American-unique (-specific) as well as the western Eurasian-unique (-specific) sequence types that can be directly derived from the subgeneric consensus, often still found in *F. crenata,* or individuals of (continental) *F. hayatae* (subsp. *pashanica*). Individual genes produce medium to near-unambiguous support (BS = 60–99.7) for splits connecting *F. crenata* (P14, F114), *F. hayatae* (F138, P37, P38), *F. longipetiolata* (F159) or several of the East Asian spp. (P49, P52, P72, F114) to the western Eurasian or (part of) the American spp. while excluding all others. The observed intra-individual gene polymorphism especially in *F. crenata* coupled with only few *crenata*-unique species-conserved mutations provides direct evidence for a polymorphic ancestor or multiple ancestors. The individual gene patterns furthermore show that there must have been several phases of gene flow between the precursors of the modern East Asian spp., especially between *F. lucida* + *F. longipetiolata* and *F. crenata*, while the insular *F. hayatae* (subsp. *hayatae*) has been more isolated (thus, placed as sister to the rest because of local LBA; note the 'Disjunct pseudoclade' in **Fig. S20**). ITS (Denk et al., 2005; Grimm et al., 2007), 5S-IGS (this study), *LEAFY* intron 2 (Oh et al., 2016; Worth et al., 2021), and Jiang et al.'s (2021) 28-gene data suggest that the nucleome of *F. crenata* is still relatively close to the likely polymorphic and widespread ancestor(s) of all modern Eurasian species of Subgenus Fagus (cf. Denk & Grimm, 2009), while especially the Chinese populations of *F. hayatae* (subsp. *pashanica*) represent genetic relics. Their insular, morphologically indistinguishable counterpart represents a genetically isolated and more derived (bottlenecks and small population size) lineage. Genetically, the morphologically consistent morphospecies *F. hayatae* is strikingly incoherent, and its continental (*F. pashanica*) and insular populations (*F. hayatae* s.str.) should be treated as distinct cryptic species (**Fig. S21**; see also **Table S3**). The putative sister species *F. lucida* and *F. longipetiolata* appear to represent two independently evolved lineages that underwent more recent gene flow (cf. **Fig. S18**): only four genes produce BS > 25 for a *longipetiolata-lucida* clade and individuals of either species may carry alien genotypes.

A best-fitting species tree should place *F. crenata* in-between *F. sylvatica* s.str. and the remaining East Asian species. The latter should form a soft 4-tip polytomy, as the inter-relationships between *F. hayatae* s.str., *F. longipetiolata, F. lucida,* and *F. pashanica* cannot be resolved using a dichotomous tree-model. The failure to infer such a more realistic tree instead of the fully resolved, high supported tree based on the combined data (Jiang et al., 2021), relates to secondary gene flow that had a partly homogenizing effect on the East Asian gene pools as well as local, gene-wise LBA/ 'short-branch culling'.

***Budding of western Eurasian beech***—The western Eurasian species share a direct common origin (fossil *F. haidingeri*) and evolved from an East Asian stock (fossil *F. castaneifolia*; cf. Denk et al., 2005; Denk & Grimm, 2009). Among all extant Eurasian species, *F. crenata* is still closest to the common ancestor and the closest living relative of the western Eurasian species. Overall, the 28-gene data, providing a larger species sample, are in very good agreement with the here observed 5S-IGS patterns despite the much lower signal amplitude (and potential detection artefacts) in Jiang et al.'s (2021) 28-gene data. Irrespective of its placement in the combined tree or the super-network (**Fig. S20**; see preceding section), *F. crenata* hence remains the most probable sister species of the west-Eurasian *F. sylvatica* s.l. (cf. in-text fig. 1). The evolutionary pathway, the transition from a *F. crenata*-close ancestor to *F. sylvatica* s.l., is gradual. Jiang et al.'s (2021) western (i.e. Western *orientalis*) and southern Turkish individuals (*F. orientalis* of unclear affinity, possibly hybrid; cf. Gömöry & Paule, 2010) are typically closer to the (ancestral) East Asian sequence variants than the western European individuals (*F. sylvatica* s.str.) The Eastern *orientalis* (represented by Iranian samples in our data) has yet to be screened at least for some of Jiang et al.'s gene sample. Based on recently assembled nuclear intron data (*Crabs Claw* and *LEAFY*; Oh et al., 2016; Renner et al., 2016; Worth et al., 2021), it can be expected that the Eastern *orientalis* are closer to the Eurasian common ancestor and *F. crenata*, show no (or very little) North American influence (discussed above), and that one may find more links to discrete East Asian species in individual genes.

The 28-gene data compiled by Jiang et al. (2021) fully supports the early, originally only ITS-based hypothesis (Denk et al., 2002, 2005), now corroborated by first nuclear intron and 5S-IGS data, that the modern species are genetic mosaics and the product of a complex reticulate history involving several alternating phases of

- gene flow via hybridisation/ introgression, facilitated by wind-pollination; and
- genetic drift during local isolation and speciation caused by climate-driven area fragmentation.

Reflecting this dynamic past, beech genomes are highly polymorphic and show high capacity to not only retain ancestral and derived gene copies but also to carry and pass on genetic signatures from different sources, and possibly involving extinct lineages. While fast-evolving, single-copy gene regions may be homogenised by later processes (e.g. latest episode of contact and gene flow), some slow-evolving gene regions may still reflect gradual replacement processes or ancient hybridisation/ introgression events (see mutation patterns tabulated for each gene in *Data S5*, sheets '… 2ISP'). Therefore, analysing nuclear data of beeches is challenging; inter-relationships of the modern species coalesce to a species network and not a species tree. However, this can easily be overlooked when data merely are combined. In the case of *Fagus,* any analysis using multiple (nuclear) genes producing a fully resolved, high-supported tree must be extremely biased and will be incomprehensive. Some topological details of the inferred tree may be strongly misleading (e.g. East Asian pseudoclade, sister-relationship between *F. longipetiolata* and *F. lucida*). Cloudograms and coalescent species tree analyses may be more comprehensive regarding alternative phylogenetic links but cannot discriminate between lack of signal, coexistence of ancestor-descendant gene copy pairs, and signal conflict, disparate gene histories. A comprehensive exploratory data re-analysis with focus on intra-individual (intra-genomic) variation is indispensable (**Figs S16–S21**; *Data S5*).

#### 5. Comparison with Other Fagaceae/Fagales

With two 5S loci (and four NORs), *Fagus* is unique within Fagaceae. The 27 *Quercus*, seven *Castanopsis*, four *Lithocarpus*, four *Castanea* and one *Trigonobalanus* species investigated so far by fluorescent in-situ hybridisation (FISH) largely showed a single pericentromeric 5S rDNA locus (www.plantrdnadatabase.com; accessed 15/08/2020; Chokchaichamnankit & Anamthawat-Jonsson, 2015). Only single individuals showed an additional locus (*Castanea mollissima*) or odd numbers of (unpaired) loci (*Lithocarpus vestitus, Quercus suber*), as a likely result of inter-specific hybridisation or autopolyploidisation (Chokchaichamnankit et al., 2008; Ribeiro et al., 2011). No comprehensive data are currently available for a comparison with other families within Fagales with the exception of *Corylus*, Betulaceae, showing a single 5S locus and a much lower intra-individual, intra-specific divergence (Forest & Bruneau, 2000).

Our data revealed five main phylogenetic lineages within the 5S-IGS gene pool of *Fagus*: O-, I-, and X-Lineage in *F. japonica*; A- and B-Lineage in the *crenata-sylvatica* lineage. In addition, we recovered copies from a likely pseudogenic lineage, the Relict Lineage. Intra-specific to intra-genomic 5S-IGS sequence polymorphism (5S-IGS "paralogy"; see preceeding section) is a common and long-known feature of Fagales (*Corylus*, Betulaceae: Forest and Bruneau 2000; *Quercus petraea*, *Q. robur,* Fagaceae: Muir et al. 2001). Cloned sequence data demonstrating the extent of 5S-IGS polymorphism are further available for all species of western Eurasian oaks (*Quercus*; Denk & Grimm 2010, >900 sequences; complemented by Simeone et al. 2018; see Piredda et al. 2021 for first HTS data) and numerous species of Betulaceae (Forest et al., 2005). The sequence divergence (*Data S4*) observed in our beech sample largely matches that of oaks and corresponds to inter-generic divergence in Betulaceae. The divergence between 'outgroup' O-Lineage and 'ingroup' X-I-A-B lineage is higher than between genera of the same Betulaceae subfamily; it approaches the divergence found between oak subgenera and sections, lineages with roots in the Eocene-Oligocene (Hubert et al., 2014; Hipp et al., 2020).

The divergence between A- and B-Lineage variants exceeds inter-species differentiation in oaks (cf. Denk & Grimm, 2010, fig. 2) and matches inter-generic differences found between *Corylus* and other members of the Coryloideae (cf. Forest & Bruneau, 2000, fig. 2). The species studied in this pilot study typically have large and stable population sizes, and are widespread in parts of temperate to boreal Eurasia (e.g. Peters, 1997). Beeches are wind-pollinated and disperse short-range via seeds (jaybirds being probably the most effective dispersal vector; e.g. Ridley, 1930; Johnson & Adkisson, 1985) and including mast years with extreme seed production (see e.g. Hilton & Packham, 2003 for a historical review). Within their climax climates, they are the dominant tree species including relict areas such as the Caucasus (see e.g. Denk et al., 2001 for information on Georgian relict forests). Thus, one cannot expect extreme levels of genetic drift and can conclude that the main 5S-IGS sequence types reflect deep splits (much) predating the formation of extant species.

#### 5.1. Evidence for sequence degradation (pseudogeny)

In some cases, increased 5S-IGS diversity has been linked to pseudogeny. Little is known about the GC content in functional 5S repeat units and its non-coding intergenic spacers (Symonová, 2019). The 5' and 3' part of the 5S rRNA gene is highly conserved and identical in *Fagus* (our data) and *Quercus* (reference accessions AJ242950, AJ242948). Increased numbers of mutations that may be detrimental for the functioning of the 5S rRNA genes appear to be confined to (very) rare variants (cf. *Data S4*; ODA subfolder *4693Data*). Exceptions are 'European O', Relict Lineage and 'Crenata A' variants, which commonly show signs of sequence degradation, especially also in the flanking 5S rRNA genes (**Table S5** gives an overview). As a general trend, potentially pseudogenic transitions (consensual C→T, consensual G→A) are more often found in *F. japonica* O-type than in its I-type variants (note the higher spread of GC contents for types of the O- vs. I-Lineage; **Section 4.1, Fig. S9**); a similar observation can be made for *crenata-sylvatica* A-Lineage (more common) and B-Lineage types (rare; see also **Section 4.2**, **Fig. S13**).

**Table S5 | Number of visibly pseudogenic sequence variants**; variants identified by compilation of sequence patterns (*Data S4*) and/or showing markedly deviating flanking gene regions in bird's eye view of the block-aligned data (NEXUS-file included in ODA). NIV = Number of variants, TA = total abundance (sum of all variants); percentages refer to the total of the given lineage/type. NIV and TA represent minimum approximates.

| Lineage | Class (NV) | NV | TA |
|---|---|---|---|
| Japonica O | Specific japonica | 14 | 226 |
| European O | Ambiguous Greek orientalis-sylvatica (3)<br>Specific crenata (2)<br>Specific Iranian orientalis (1)<br>Specific sylvatica (2) | 8 | 108 |
| Relict Lineage | Ambiguous west-Eurasian (1)<br>Ambiguous Iranian orientalis-sylvatica (2)<br>Ambiguous Greek orientalis-sylvatica (1)<br>Specific crenata (2)<br>Specific Iranian orientalis (2)<br>Specific sylvatica (5) | 13 | 164 |
| Unique variant ('not-I') | Specific sylvatica (1) | 1 | 5 |
| A-Lineage | Ambiguous west-Eurasian (1)<br>Ambiguous Greek orientalis-sylvatica (4)<br>Specific crenata (8)<br>Specific Greek orientalis (3)<br>Specific sylvatica (1) | 17 | 1147[a] |
| B-Lineage | Specific crenata (1)<br>Specific Iranian orientalis (3)<br>Specific Greek orientalis (3)<br>Specific sylvatica (5) | 12 | 160 |

[a] High number due to one variant shared by Greek *F. orientalis* and *F. sylvatica* s.str., the most common "ambiguous" variant of the *crenata-sylvatica* lineage ("ASOG" in the 38-tip set; TA = 1016)

Since the HTS approach is amplicon-based, highly pseudogenic 5S rDNA arrays will not be captured, or with much-decreased efficiency. While pseudogeny can hinder phylogenetic tree inference because of the conflicting signal from pseudogenic mutations and risk of long-branch attraction, pseudogenic nuclear spacer data have proven to be highly informative regarding past reticulations and identification of parentage in stabilized allopolyploids (e.g. Hugall et al., 1999; Manen, 2004; Won & Renner, 2005; Grimm & Denk, 2008; Vierna et al., 2013; Volkov et al., 2017).

#### 6. List of Included Appendices

**Appendix A | Summary of *Data S2.*** Amplicon GC content and length ranges for major sequence types; total abundance and relative proportion.

**Appendix B | Violin and scatterplots of amplicon GC content and length**; sorted by samples and main 5S-IGS lineages.

**Appendix A.** General structural features (amplicon length and GC content), number of non-identical variants (NIV), and (proportional) total abundance (TA/PA) of main types per sample**.** Dominant and co-dominant (regarding both diversity and abundance) types highlighted by grey shading.

| Sample | Type | GC content [%] Median | GC content [%] Range | Length [bp] Median | Length [bp] Range | Sum[a] NIV | Sum[a] TA | Sum[a] PA | Shared[b] 'Original A/B' NIV | Shared[b] 'Original A/B' PA | Specific 'Iranian B1' NIV | Specific 'Iranian B1' PA | Specific 'Crenata B' NIV | Specific 'Crenata B' PA | Specific 'European A/B' NIV | Specific 'European A/B' PA |
|---|---|---|---|---|---|---|---|---|---|---|---|---|---|---|---|---|
| 06 – *Fagus japonica* | O | 40.9 | 38.6–42.5(–44.3) | 306 | 262–307 | **398** | 9593 | **30%** | *Japonica*- specific | | | | | | | |
| | Short O | 43.6 | 41.7–45.1 | 204 | 203–204[e] | 97 | 1854 | 6% | *Japonica*- specific | | | | | | | |
| | I | 38.6 | 36.1–40.8 | 262 | 197–264 | **386** | 20116 | **62%** | *Japonica*- specific | | | | | | | |
| | X | 38.1 | 35.9–38.9 | 260 | 258–262 | 26 | 662 | 2% | *Japonica*- specific | | | | | | | |
| 05 – *F. crenata* | O | – | 38.2; 41.0; 42.6 | – | 266; 304 | 3 | 30 | 0.1% | Sister lineage of 'Japonica O' | | | | | | | |
| | A | 36.5 | 33.2–38.8 | 274 | 259–283(296) | 31 | 199 | 0.7% | 29 | 0.7% | – | – | – | – | 0 | 0% |
| | B[c] | 37.8 | 33.8–39.9 | 261 | 252–272 | 1215 | 27098 | 99% | 10 | 16.8% | 1 | 0.03% | **1204** | **82%** | 0 | 0% |
| | Relict lin. | – | 35.1–39.6 | – | 259–268 | 5 | 48 | 0.2% | Rare, derelict type of the *crenata-sylvatica* lineage | | | | | | | |
| 04 – Iranian *F. orientalis* | O | – | 39.7; 40.1; 42.3 | – | 307; 304; 260 | 3 | 24 | 0.1% | Sister lineage of 'Japonica O' | | | | | | | |
| | A | 36.6 | 34.8–38.4 | 269 | 258–279 | 388 | 8466 | 42% | **386** | **42%** | – | – | – | – | 0 | 0% |
| | B | 37.4 | 35.3–39.4 | 256 | 247–269 | 310 | 11706 | 58% | **296** | **52%** | 13 | 5% | 1 | 0.005% | 0 | 0% |
| | Relict lin. | – | 36.4–39.8 | – | 256–269 | 5 | 31 | 0.2% | Rare, derelict type of the crenata-sylvatica lineage | | | | | | | |
| 11 – Greek *F. orientalis* | O | – | 37.5–41.7 | – | (266)303–304 | 8 | 26 | 0.1% | Sister lineage of 'Japonica O' | | | | | | | |
| | A | 36.8 | 34.9–39.6 | 273 | 258–279 | 235 | 4932 | 27% | 30 | 2% | – | – | – | – | **216** | **25%** |
| | Short A | 39.8 | 38.6–41.0 | 166 | 166 [constant] | 17 | 978 | 5% | 0 | 0% | – | – | – | – | 17 | 5% |
| | B[d] | 37.9 | 33.6–41.1 | 261 | (177–)231–267 | 334 | 12546 | 68% | 149 | 23% | 2 | 0.1% | 1 | 0.03% | **180** | **45%** |
| | Relict lin. | – | 36.9–40.0 | – | 256–268 | 4 | 25 | 0.1% | Rare, derelict type of the crenata-sylvatica lineage | | | | | | | |
| 14 – *F. sylvatica* s.str. (Italy) | O | – | 37.5–40.1(41.7) | – | 303–304(266) | 9 | 33 | 0.3% | Sister lineage of 'Japonica O' | | | | | | | |
| | A | 36.6 | 34.4–39.6 | 274 | 258–280 | 241 | 4142 | 38% | 10 | 1% | – | – | – | – | **228** | **37%** |
| | Short A | – | 40.4 | – | 166 | 2 | 12 | 0.1% | 0 | 0% | – | – | – | – | 2 | 0.1% |
| | B | 38.3 | 35.7–41.5 | 261 | 212–262 | 312 | 6605 | 61% | 57 | 12% | 1 | 0.04% | 0 | 0% | **254** | **49%** |
| | Relict lin. | – | 35.8–39.8 | – | 256–268 | 10 | 28 | 0.3% | Rare, derelict type of the crenata-sylvatica lineage | | | | | | | |
| 12 – *F. sylvatica* s.str. (Germany) | O | – | 37.5–40.1(41.7) | – | 303–304(266) | 8 | 63 | 0.2% | Sister lineage of 'Japonica O' | | | | | | | |
| | Not I[e] | – | 36.0 | – | 253 | 1 | 5 | 0.01% | Ingroup relict type of unclear affinity | | | | | | | |
| | A | 36.6 | 34.7–39.1 | 274 | 258–280 | 308 | 6176 | 17% | 7 | 0.1% | – | – | – | – | **298** | 17% |
| | B | 38.5 | 35.4–40.8 | 261 | 243–262 | 844 | 30142 | 83% | 18 | 0.6% | 5 | 0.1% | 0 | 0% | **820** | **82%** |
| | Relict lin. | – | 35.8–39.8 | – | 256–268 | 11 | 90 | 0.2% | Rare, derelict type of the crenata-sylvatica lineage | | | | | | | |

[a] Sum includes variants placed at the root branch of the respective clades.

[b] Far the most 'Oriental A' types are specific for Iranian *F. orientalis* (see Figs 2, 3)

[c] Most variants are of type 'Crenata B2' (total abundance = 21609)

[d] Includes one variant with an abundance = 9 (GC content = 38.1%; length = 260) placed at the I-B-clade root.

[e] See *Supplementary file S4* ("SyG3")

[f] Corrected for wrongly clipped 5' ends found in four 'short O' variants (see Supplementary files S1, S2)

**Appendix B | Violin and scatterplots including all 5S-IGS types**

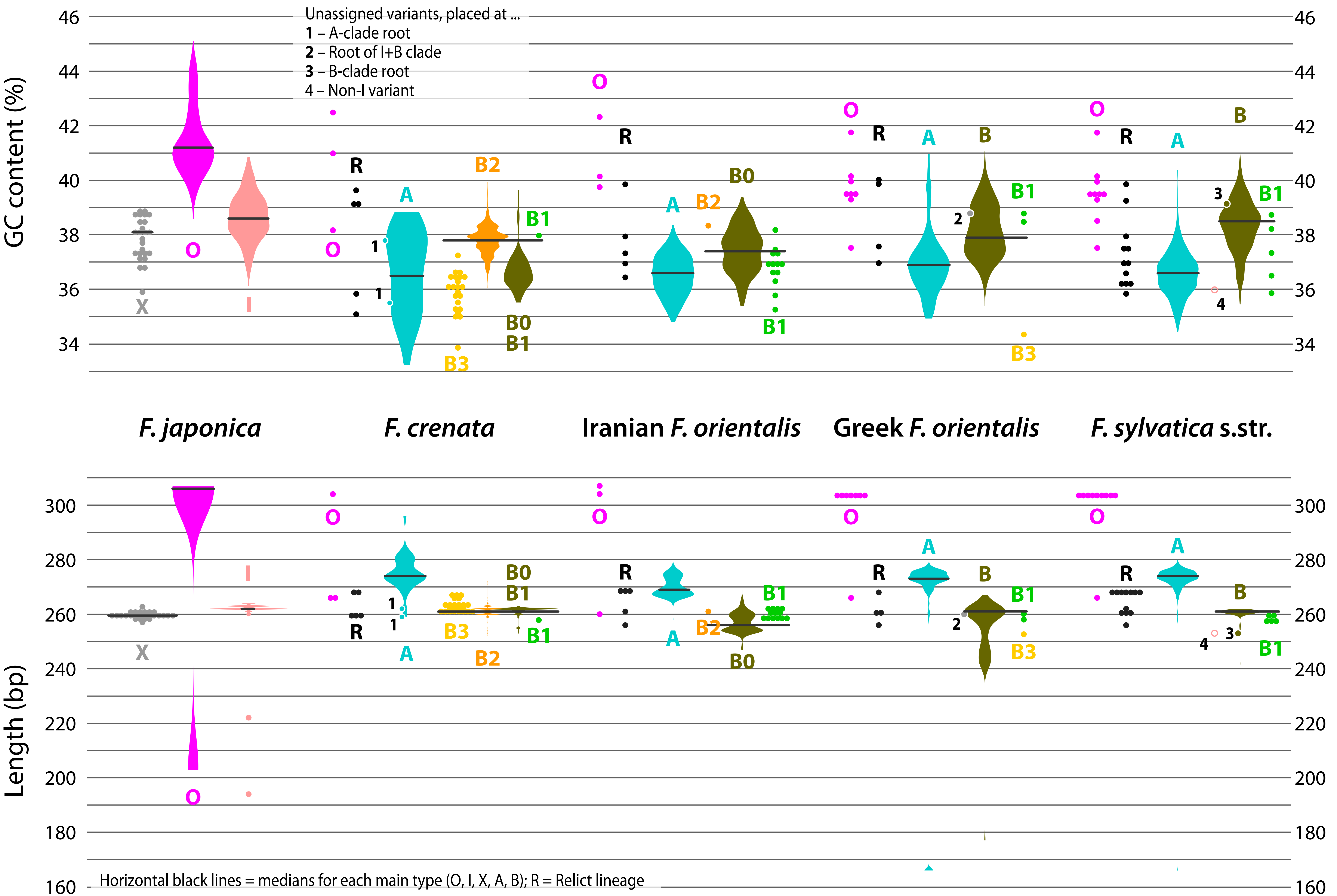
