## Supplementary material for "5S-IGS rDNA in wind-pollinated trees (*Fagus* L.) encapsulates 55 million years of reticulate evolution and hybrid origins of modern species": Data S3

EPA (Evolutionary Placement Analysis) of the "Ambiguous" sequence reads with abundance < 25; 144 total variants with a cumulative abundance of 1304 reads, shared by *F. sylvatica* and *F. orientalis* samples. These sequences can be subdivided in three classes ("Western", "Ancient", "European")

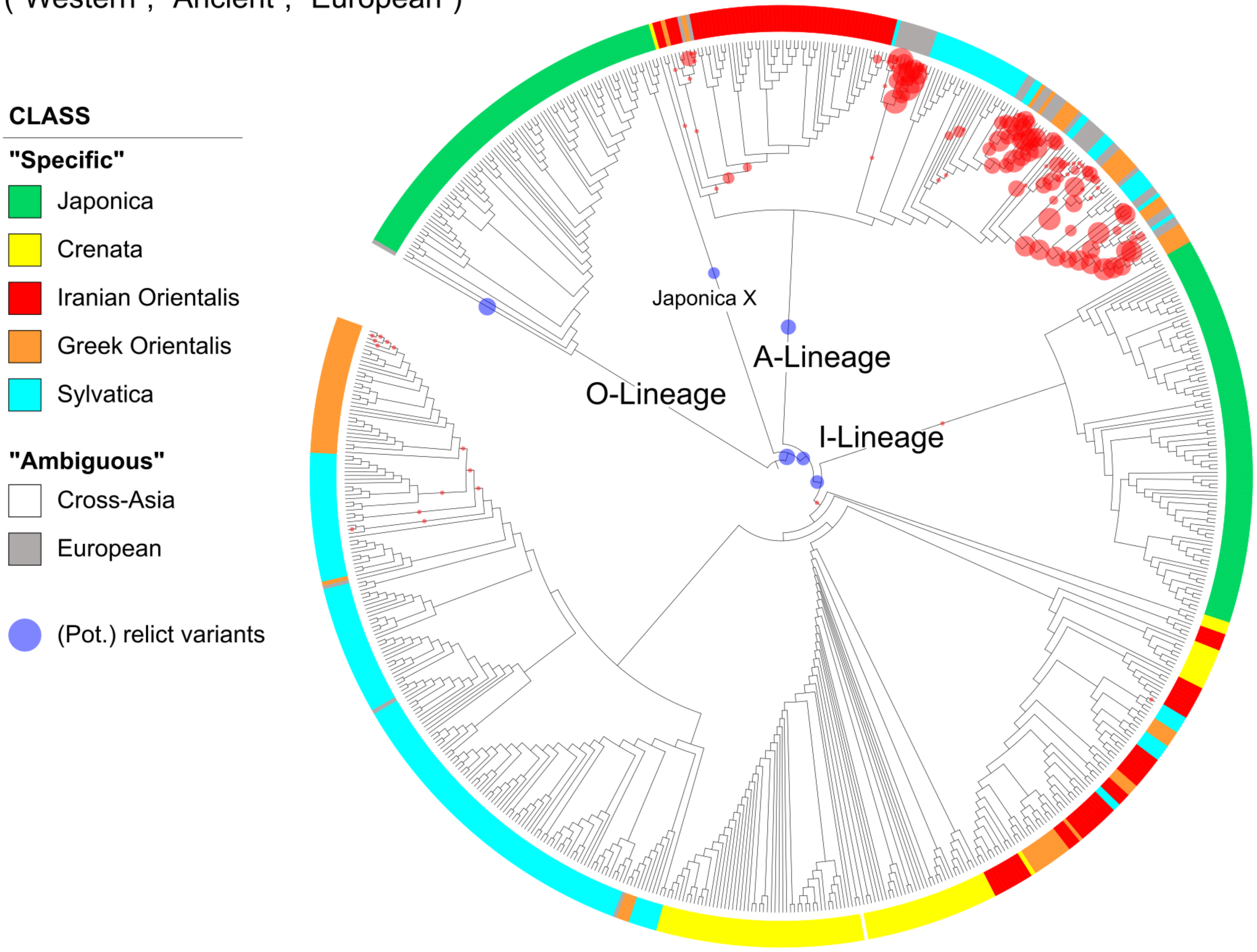

EPA (Evolutionary Placement Analysis) of sequence reads with abundance < 25
a.1) Class "Western" (three variants with a cumulative abundance of 21);
shared across all *F. sylvatica* s.l. samples.

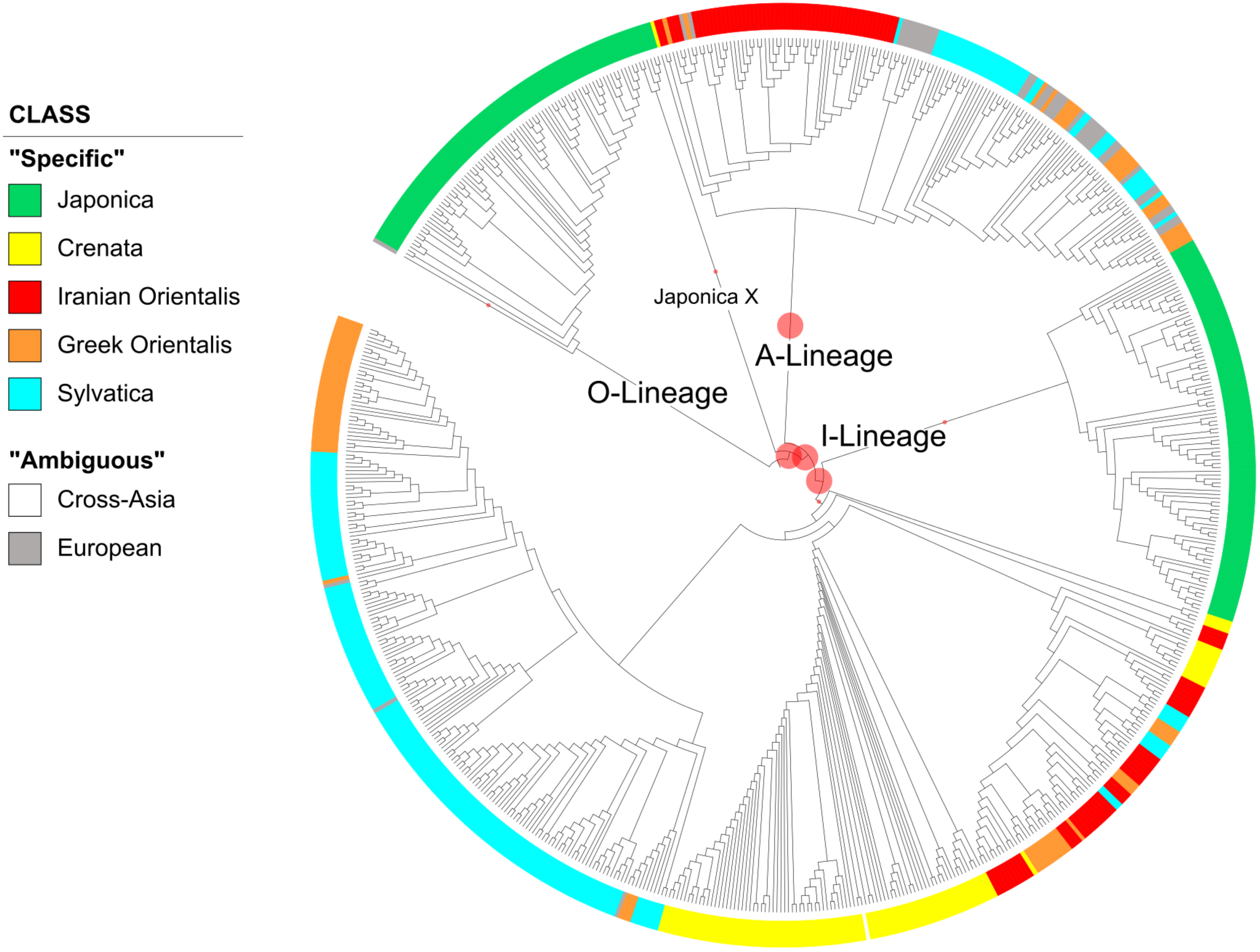

EPA (Evolutionary Placement Analysis) of sequence reads with abundance < 25
a.2) Class "Ancient" (three variants with a cumulative abundance of 45);
shared by *F. sylvatica* s.str. and Iranian *F. orientalis*

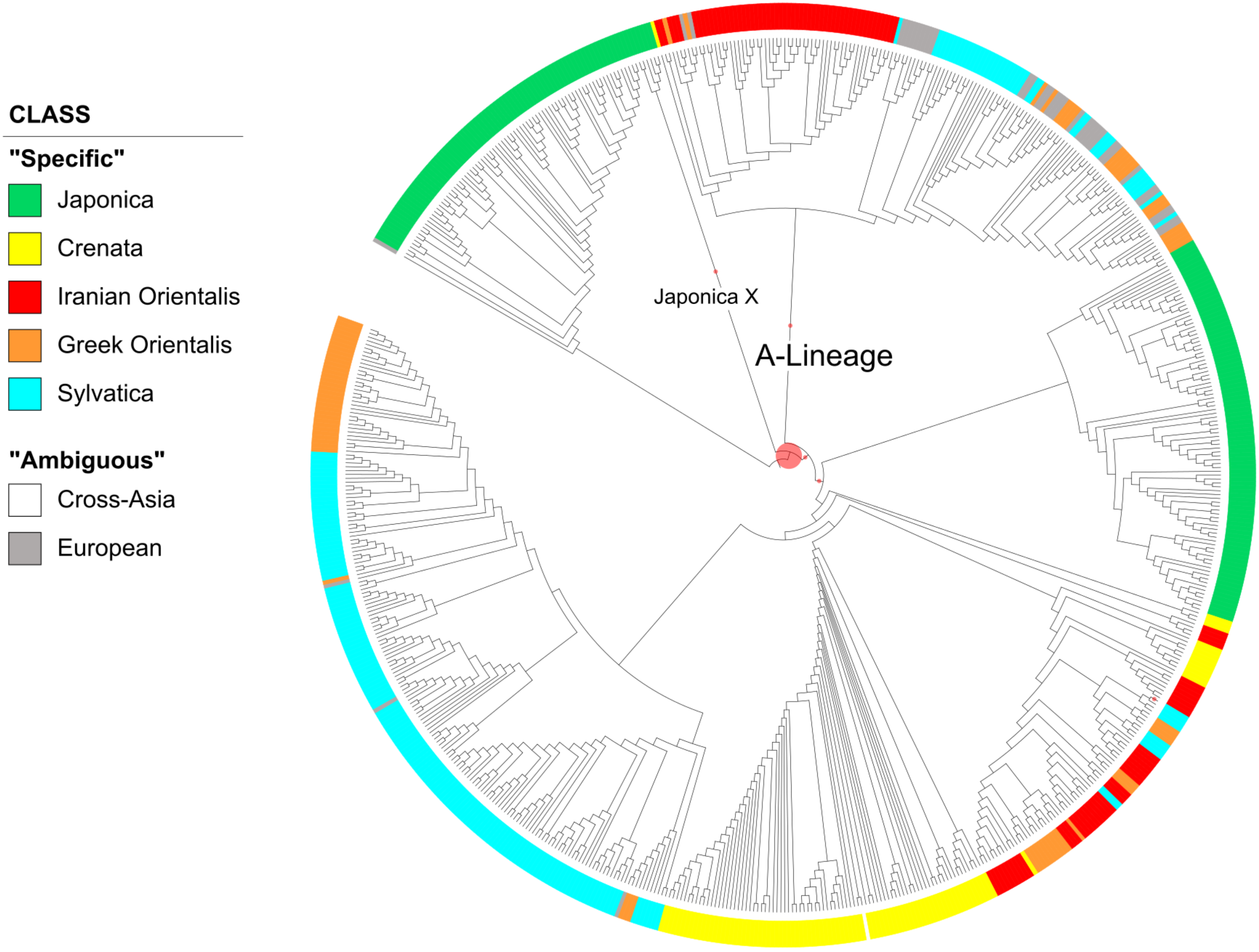

EPA (Evolutionary Placement Analysis) of sequence reads with abundance < 25
a.3) Class "European" (138 variants with a cumulative abundance of 1238);
shared by *F. sylvatica* s.str. and Greek *F. orientalis*.

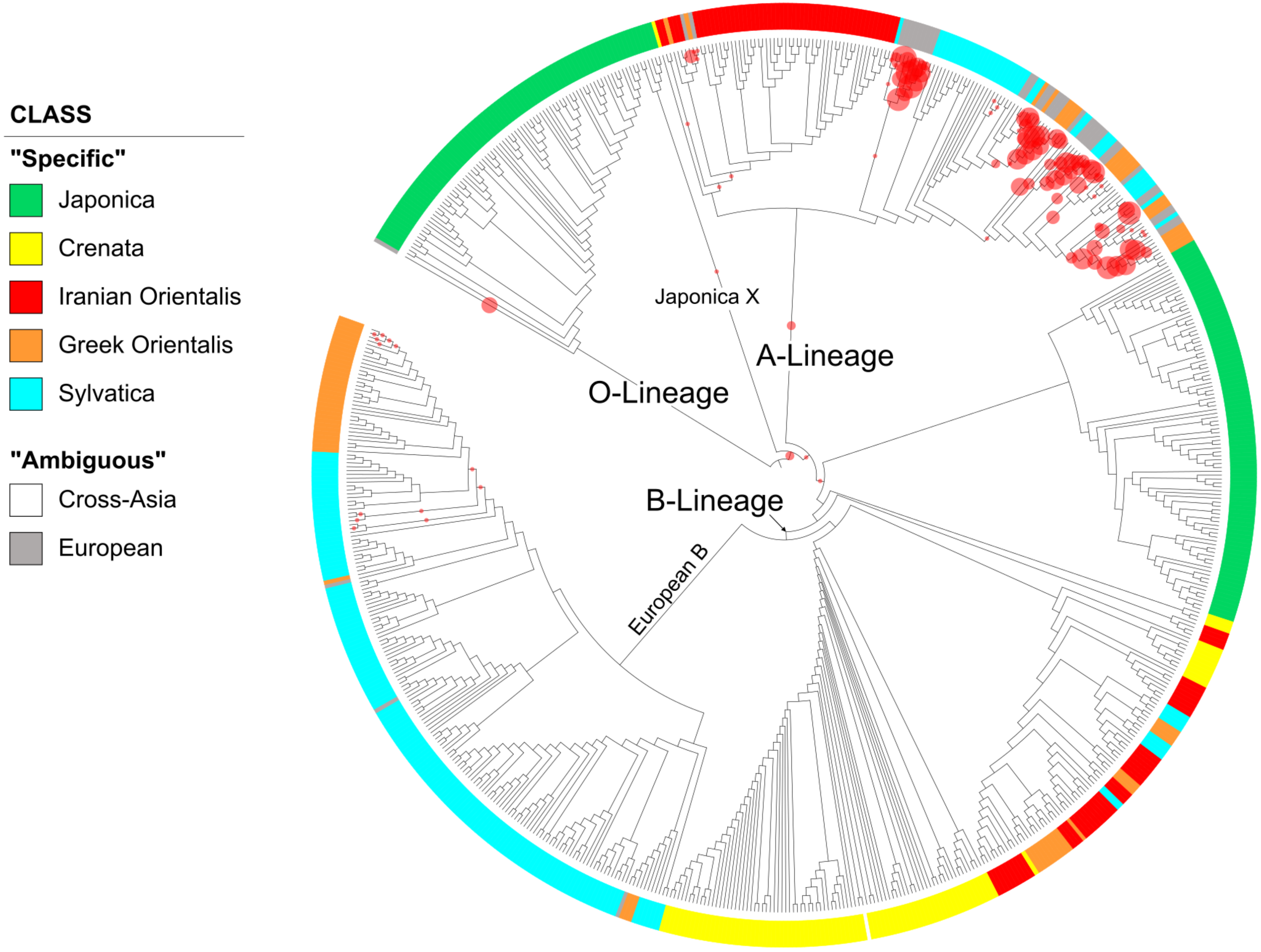

EPA (Evolutionary Placement Analysis) of sequence reads with abundance < 25
b) Class "Crenata" (1154 variants with a cumulative abundance of 7840);
only found in the *F. crenata* sample

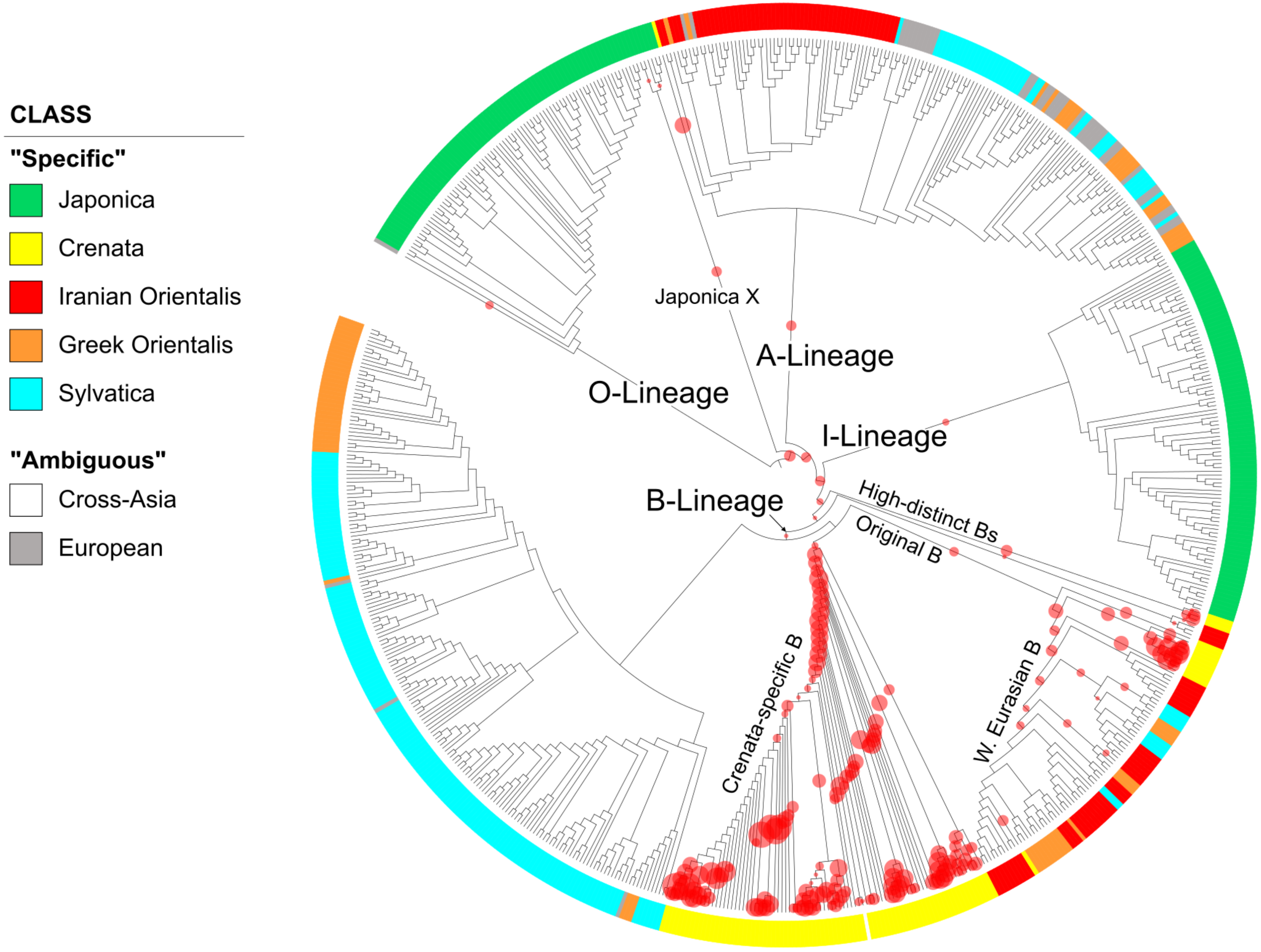

EPA (Evolutionary Placement Analysis) of sequence reads with abundance < 25
c) Class "Japonica" (727 variants with a cumulative abundance of 5560);
only found in the *F. japonica* sample.

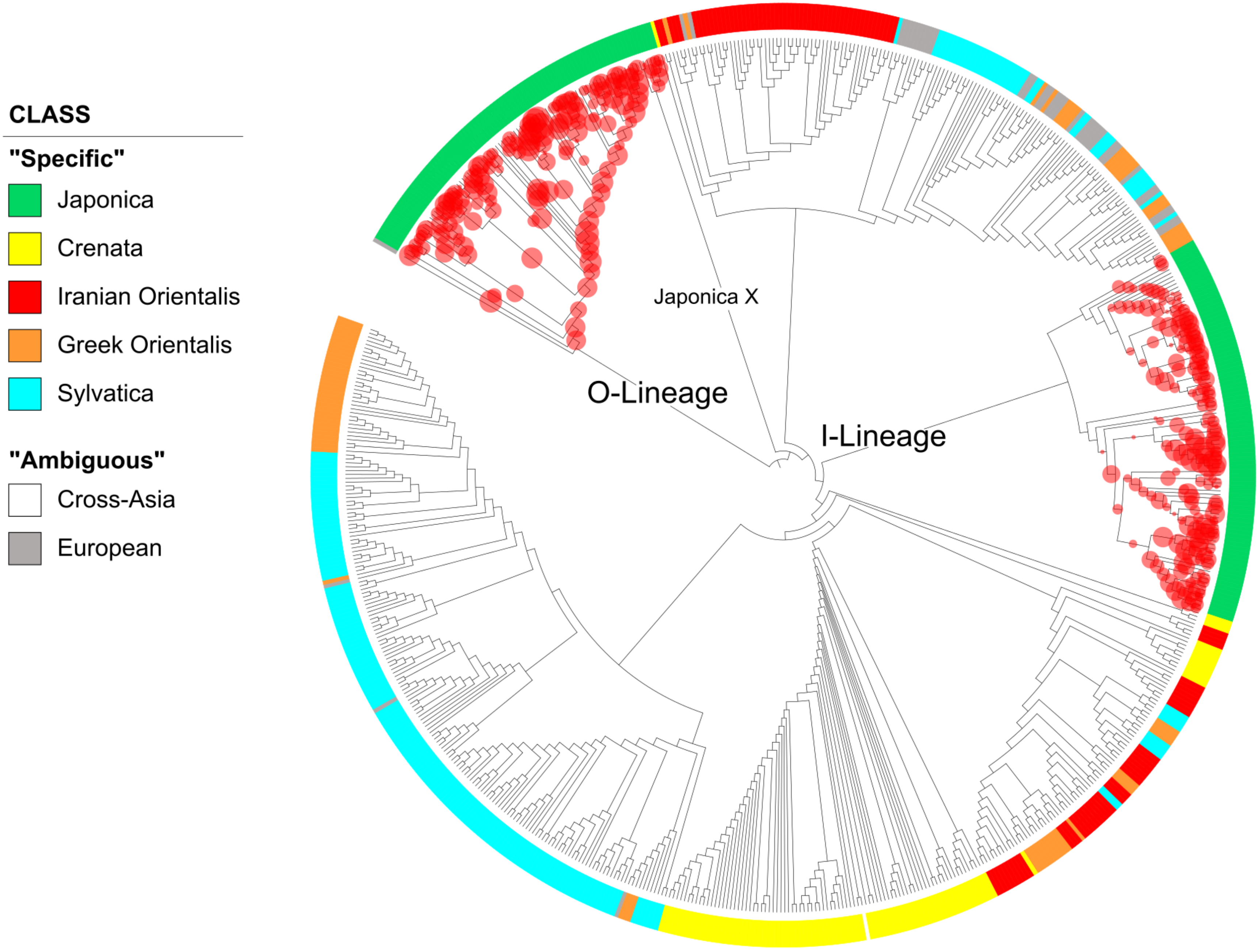

EPA (Evolutionary Placement Analysis) of sequence reads with abundance < 25
d) Class "Greek Orientalis" (356 variants with a cumulative abundance of 2566); only found in the Greek *F. orientalis* sample.

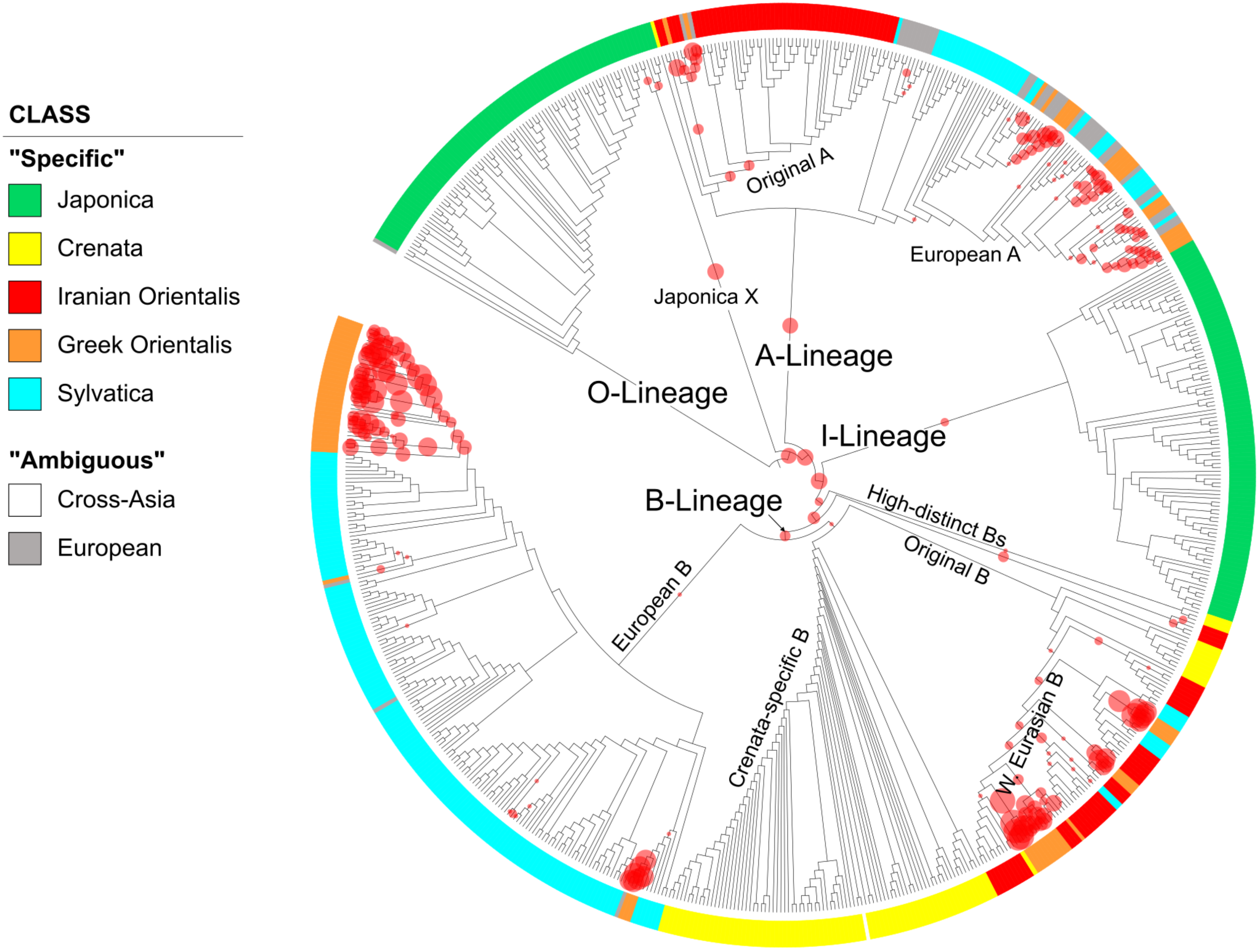

EPA (Evolutionary Placement Analysis) of sequence reads with abundance < 25
e) Class "Iranian Orientalis" (598 variants with a cumulative abundance of 4371); only found in the Iranian *F. orientalis* sample.

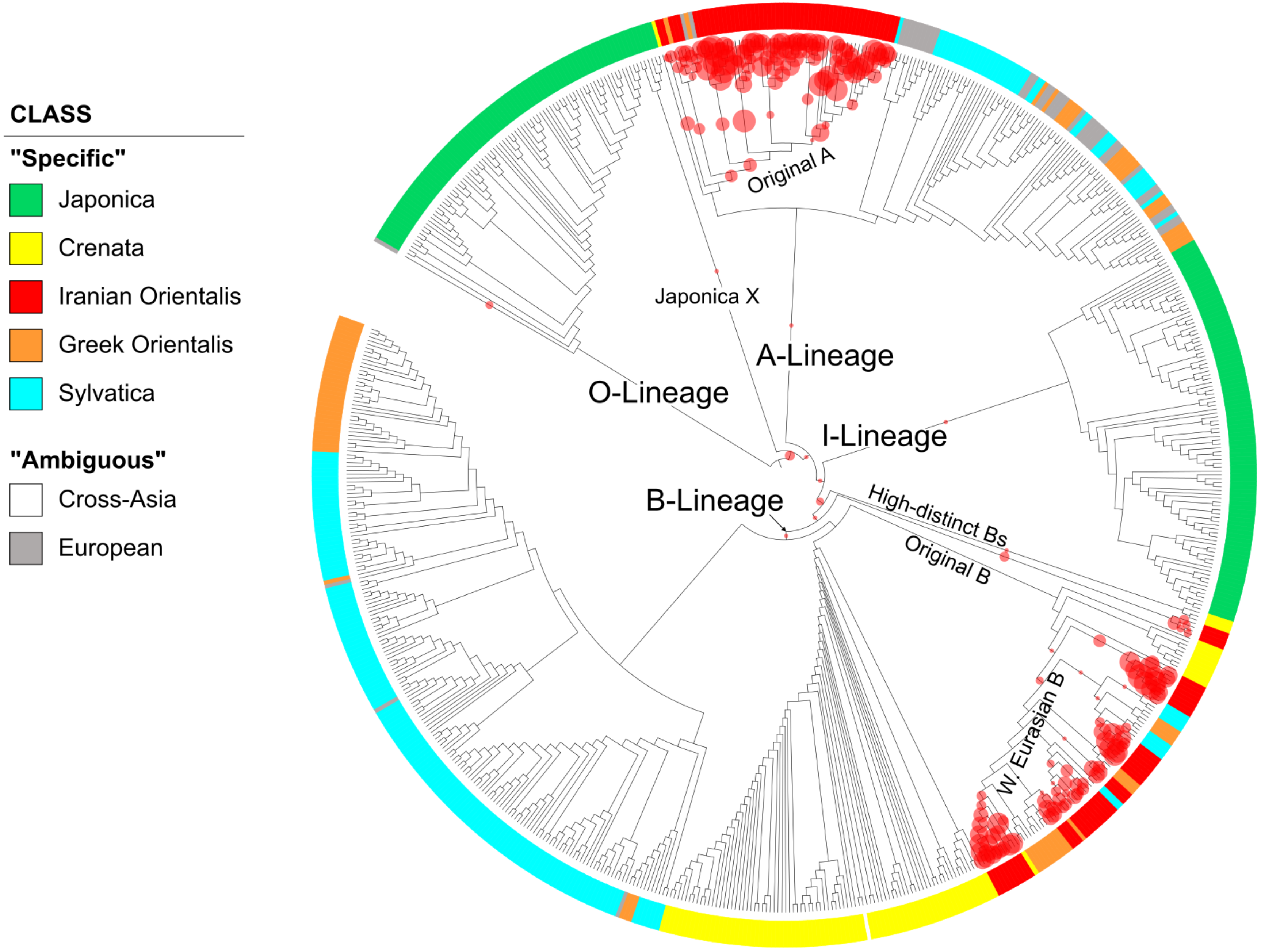

EPA (Evolutionary Placement Analysis) of sequence reads with abundance < 25
f) Class "Sylvatica" (1028 variants with a cumulative abundance of 8229);
only found in *F. sylvatica* s.str. samples.

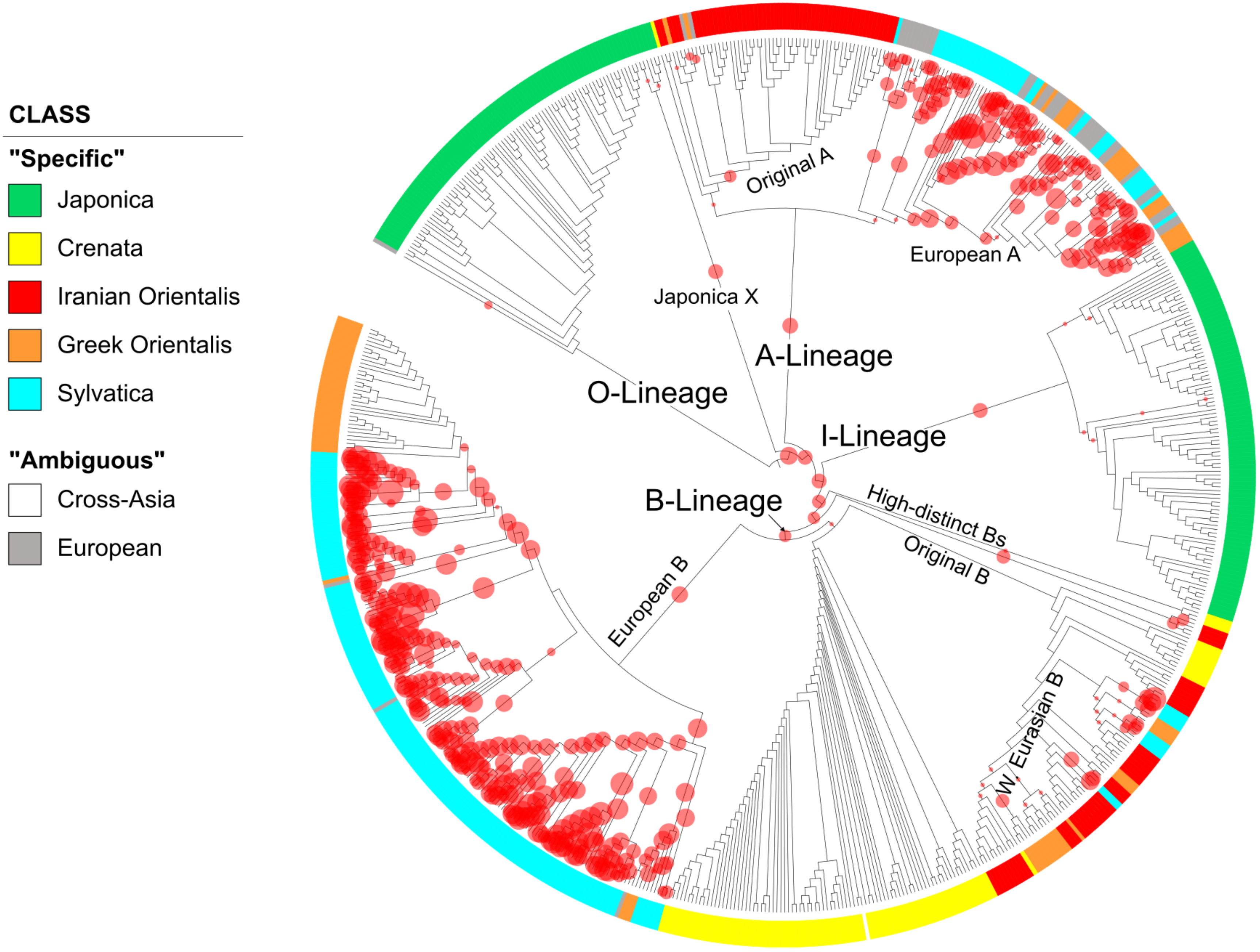
